# *Botrytis cinerea* identifies host plants via the recognition of antifungal capsidiol to induce expression of a specific detoxification gene

**DOI:** 10.1101/2022.05.11.490027

**Authors:** Teruhiko Kuroyanagi, Abriel Bulasag, Keita Fukushima, Takamasa Suzuki, Aiko Tanaka, Maurizio Camagna, Ikuo Sato, Sotaro Chiba, Makoto Ojika, Daigo Takemoto

## Abstract

The gray mold pathogen *Botrytis cinerea* has a broad host range, causing disease in over 400 plant species, but it is not known how this pathogen evolved this polyxenous nature. *B. cinerea* can metabolize a wide range of phytoalexins, including the stilbenoid, resveratrol, and the sesquiterpenoids capsidiol in tobacco, and rishitin in potato and tomato. In this study, we analyzed the metabolism of sesquiterpenoid phytoalexins by *B. cinerea*. Capsidiol was dehydrogenated to capsenone which was then further oxidized, while rishitin was directly oxidized to epoxy-or hydroxy-rishitins indicating that *B. cinerea* has separate mechanisms to detoxify structurally similar sesquiterpenoid phytoalexins. RNAseq analysis revealed that a distinct set of genes were induced in *B. cinerea* when treated with capsidiol or rishitin, suggesting that *B. cinerea* can distinguish structurally similar phytoalexins to activate appropriate detoxification mechanisms. The gene most highly upregulated by capsidiol treatment encoded a dehydrogenase, designated *Bccpdh*. Heterologous expression of *Bccpdh* in a capsidiol-sensitive plant symbiotic fungus, *Epichloë festucae*, resulted in an acquired tolerance of capsidiol and the ability to metabolize capsidiol to capsenone, while *B. cinerea Δbccpdh* mutants became relatively sensitive to capsidiol. The *Δbccpdh* mutant showed reduced virulence on the capsidiol producing *Nicotiana* and *Capsicum* species but remained fully pathogenic on potato and tomato. Homologs of *Bccpdh* are not found in taxonomically distant Ascomycota fungi but not in related Leotiomycete species, suggesting that *B. cinerea* acquired the ancestral *Bccpdh* by horizontal gene transfer, thereby extending the pathogenic host range of this polyxenous pathogen to capsidiol-producing plant species.

**Significance Statement:** *B. cinerea* can metabolize a wide range of phytoalexins, however, the extent to which phytoalexin detoxification contributes to pathogenicity is largely unknown. In this study, we have shown that *B. cinerea* recognizes structurally resembling sesquiterpenoid phytoalexins, rishitin and capsidiol, to activate appropriate detoxification mechanisms. We identify *Bccpdh*, encoding a dehydrogenase for capsidiol detoxification, which is upregulated in *B. cinerea* exclusively during the infection of capsidiol producing plant species, and is required to exert full virulence. Analysis of the *Bccpdh* locus implicates that the gene was acquired via horizontal gene transfer. This work highlights that the polyxenous plant pathogen *B. cinerea* can distinguish its host plants by its anti-microbial compounds, to activate appropriate mechanisms for enhanced virulence.

## Introduction

The antimicrobial secondary metabolites produced in plants during the induction of disease resistance are collectively termed phytoalexins (1, 2). Several hundred phytoalexins of diverse structures have been identified from a wide range of plant species, which include terpenoids, flavonoids and indoles (3, 4). Many of these phytoalexins are considered to exhibit their antimicrobial activity by targeting the cell wall or cell membrane of pathogens (5, 6), but their mechanisms of action remain largely unknown.

In plants belonging to the Solanaceae family, the major phytoalexins are sesquiterpenoids, such as capsidiol in *Nicotiana* and *Capsicum* species and rishitin in *Solanum* species (2, 7, 8). In *Nicotiana* sp., production of capsidiol is strictly controlled by regulating the gene expression for capsidiol biosynthesis, such as *EAS* (5-*epi*-aristolochene synthase) and *EAH* (5-*epi*-aristolochene dihydroxylase), encoding the enzymes dedicated to the production of capsidiol (9, 10). In *N. benthamiana*, expression of *EAS* and *EAH* genes is hardly detected in healthy tissues, but their expression is rapidly induced during infection by pathogens or the treatment with elicitor protein (11, 12). Produced phytoalexins are secreted locally via ABC (ATP-binding cassette) transporters at the site of pathogen attack (13, 14). For plants, such temporal and spatial control of phytoalexins is probably critical as the toxicity of phytoalexins is often not specific to microorganisms but is also harmful to plant cells (13, 15, 16). For example, rishitin affects the permeability of plant liposomal membranes and disrupts chloroplasts (17). Therefore, timely production and efficient transport of phytoalexins to the site of pathogen attack are important for plants to apply these double-edged weapons effectively.

It has been reported that various phytopathogenic fungi can metabolize and detoxify phytoalexins (18). Although many studies have shown an approximate correlation between the ability of strains to detoxify phytoalexins and their virulence (19–21), the importance of phytoalexin detoxification for pathogen virulence is largely unproven for most plant-pathogen interactions. The best-studied phytoalexin metabolizing enzyme is pisatin demethylase (PDA) of *Nectria haematococca*. Deletion of the *PDA* gene resulted in reduced virulence of *N. haematococca* on pea, directly proving the importance of PDA for the virulence of this pathogen (22). It has also been reported that sesquiterpenoid phytoalexins are metabolized by pathogenic fungi. Capsidiol is metabolized to less toxic capsenone via dehydration by pathogens such as the gray mold pathogen *Botrytis cinerea* and *Fusarium oxysporum* (23). *Gibberella pulicaris*, the dry rot pathogen of potato tubers, oxidizes rishitin to 13-hydroxy rishitin and 11,12-epoxyrishitin (24). However, the enzymes involved in the detoxification of sesquiterpenoid phytoalexins have not been isolated to date, and their importance for pathogenicity has not been demonstrated.

In this study, we first investigated some pathogens isolated from Solanaceae and non-Solanaceae plants on their tolerance of capsidiol and rishitin, as well as their ability to detoxify/metabolize these phytoalexins. Among the pathogens that can metabolize both capsidiol and rishitin, we chose *Botrytis cinerea* for further analysis to investigate the importance of its ability to detoxify capsidiol for the pathogenicity on plant species that produce capsidiol.

## Results

### Metabolization of sesquiterpenoid phytoalexins capsidiol and rishitin by fungal plant pathogens

Four oomycetes and 12 fungal species were evaluated for their ability to detoxify/metabolize sesquiterpenoid phytoalexins capsidiol and rishitin, produced by Solanaceae plant species. Four *Phytophthora* species isolated from Solanaceae host plants, including potato late blight pathogen *P. infestans, P. nicotiana* isolated from tobacco, *P. capsici* isolated from green pepper and *P. cryptogea* isolated from nipplefruit (*Solanum mammosum*), are all sensitive to capsidiol and rishitin. The amount of capsidiol and rishitin after the incubation with these oomycete pathogens did not decrease, indicating that they are neither capable of metabolizing nor tolerant to capsidiol and rishitin (Fig. 1 and Supplemental Fig. S1). Among 12 fungal plant pathogens (eight isolated from Solanaceae plants), seven and eight fungal strains can metabolize capsidiol and rishitin, respectively, and showed increased resistance to phytoalexins. In some cases, pathogens can metabolize phytoalexins that are not produced by their host, indicating that the ability to detoxify phytoalexins does not always correlate with their host range (Supplemental Figs. S1-3 and Supplementary Notes 1 and 2). *Botrytis cinerea* and *Sclerotinia sclerotiorum* are well-known as pathogens with a wide host range and are capable of metabolizing both capsidiol and rishitin. In this study, *B. cinerea* was selected as the pathogen to further investigate the role of detoxification of phytoalexins on the pathogenicity of this polyxenous pathogen.

**Fig. 1.**
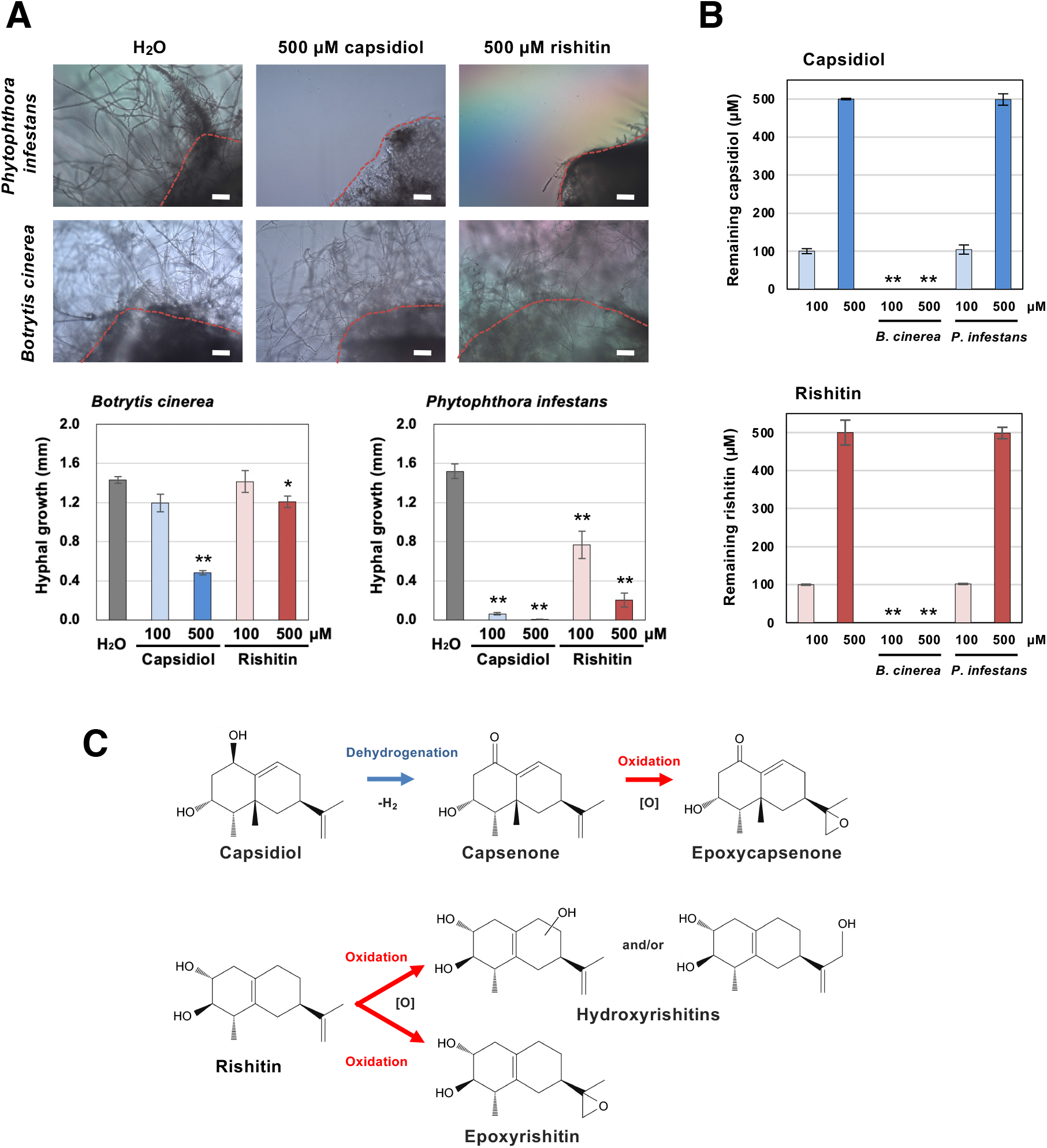
Sensitivities and metabolic capacities of *Botrytis cinerea* and *Phytophthora infestans* to sesquiterpenoid phytoalexins. **(A)** Mycelial blocks (approx. 1 mm^3^) of indicated pathogen were incubated in 50 µl water, 500 µM capsidiol or 500 µM rishitin. Outgrowth of hyphae from the mycelial block (outlined by dotted red lines) was measured after 24 h of incubation (n = 6). Bars = 100 µm. **(B)** Residual capsidiol and rishitin was quantified by GC/MS 2 days after the incubation. **(C)** Predicted metabolism of capsidiol and rishitin by *B. cinerea*. Note that the structure of oxidized capsenone was determined in this study. Data marked with asterisks are significantly different from control as assessed by the two-tailed Student’s *t*-test: **P < 0.01, *P < 0.05.

### Metabolization of capsidiol and rishitin by *B. cinerea*

Metabolization of capsidiol and rishitin by *B. cinerea* were evaluated by LC/MS. Under the experimental conditions of this study, capsidiol was metabolized to capsenone by dehydrogenation within 12 h, and oxidized capsenone was detected at 24 h, whereas oxidized capsidiol was not detected (Fig. 1C and Supplemental Figs. S4). In contrast, rishitin was directly oxidized within 6 h and at least four different forms of oxidized rishitin were detected within 24 h (Fig. 1C and Supplemental Figs. S5), indicating that despite the structural similarity of these phytoalexins, *B. cinerea* detoxifies/metabolizes capsidiol and rishitin by different mechanisms (Fig. 1C).

### Unique sets of genes are upregulated in *B. cinerea* treated with capsidiol, rishitin or resveratrol

To identify *B. cinerea* genes upregulated during the detoxification of sesquiterpenoid phytoalexins, RNAseq analysis was performed for mycelia of *B. cinerea* cultured in minimal media supplemented with capsidiol or rishitin. A stilbenoid phytoalexin, resveratrol, produced in grape was also used for comparison. Mycelia of *B. cinerea* were grown in liquid CM media supplemented with either 500 µM rishitin, 500 µM resveratrol, or 100 µM capsidiol, as 500 µM capsidiol caused significant growth defects in *B. cinerea* (Fig. 1A, Supplemental Fig. S6). The mycelial tissue was then used to perform an RNAseq analysis. Among 11,707 predicted *B. cinerea* genes (25), 25, 27 or 23 genes were significantly upregulated (Log2 fold change >2, p <0.05) by the treatment with capsidiol, rishitin or resveratrol, respectively. Unexpectedly, distinctive sets of genes were upregulated even between *B. cinerea* treated with capsidiol and rishitin which resemble each other structurally (Fig. 2A, Supplemental Table S1-3), indicating that *B. cinerea* can either distinguish the structural difference of capsidiol and rishitin or the damage caused by these sesquiterpenoid phytoalexins. For instance, two genes Bcin08g00930 and Bcin12g01750 encoding hypothetical proteins containing a motif for dehydrogenases were specifically induced by capsidiol, while the treatment of rishitin significantly induced Bcin07g05430, encoding a cytochrome P450 gene (Fig. 2B). Expression of *BcatrB* encoding an ABC transporter involved in the tolerance of *B. cinerea* against structurally unrelated phytoalexins resveratrol, camalexin and fungicides fenpiclonil and fludioxonil (26, 27), is upregulated by treatment with rishitin and resveratrol, but not by capsidiol. In contrast, Bcin15g00040, encoding a major facilitator superfamily (MFS)-type transporter was upregulated specifically by capsidiol (Fig. 2B). Interestingly, treatment of *B. cinerea* with phytoalexins also activated genes predicted to be involved in pathogenicity to plants. For example, capsidiol treatment induced expression of a hydrophobin gene Bcin06g00510.1 (Bhp3), while rishitin treatment activated the expression of Bcin01g00080.1 (Bcboa8), encoding an enzyme for biosynthesis of a phytotoxin, botcinic acid (Supplemental Tables S1 and S2). Given that capsidiol is metabolized to capsenone by a dehydrogenation reaction in *B. cinerea* (Fig. 1C, Supplemental Fig. S4), the function of Bcin08g00930 and Bcin12g01750, both encoding a predicted short-chain dehydrogenase/reductase (SDR) were further analyzed in this study.

**Fig. 2.**
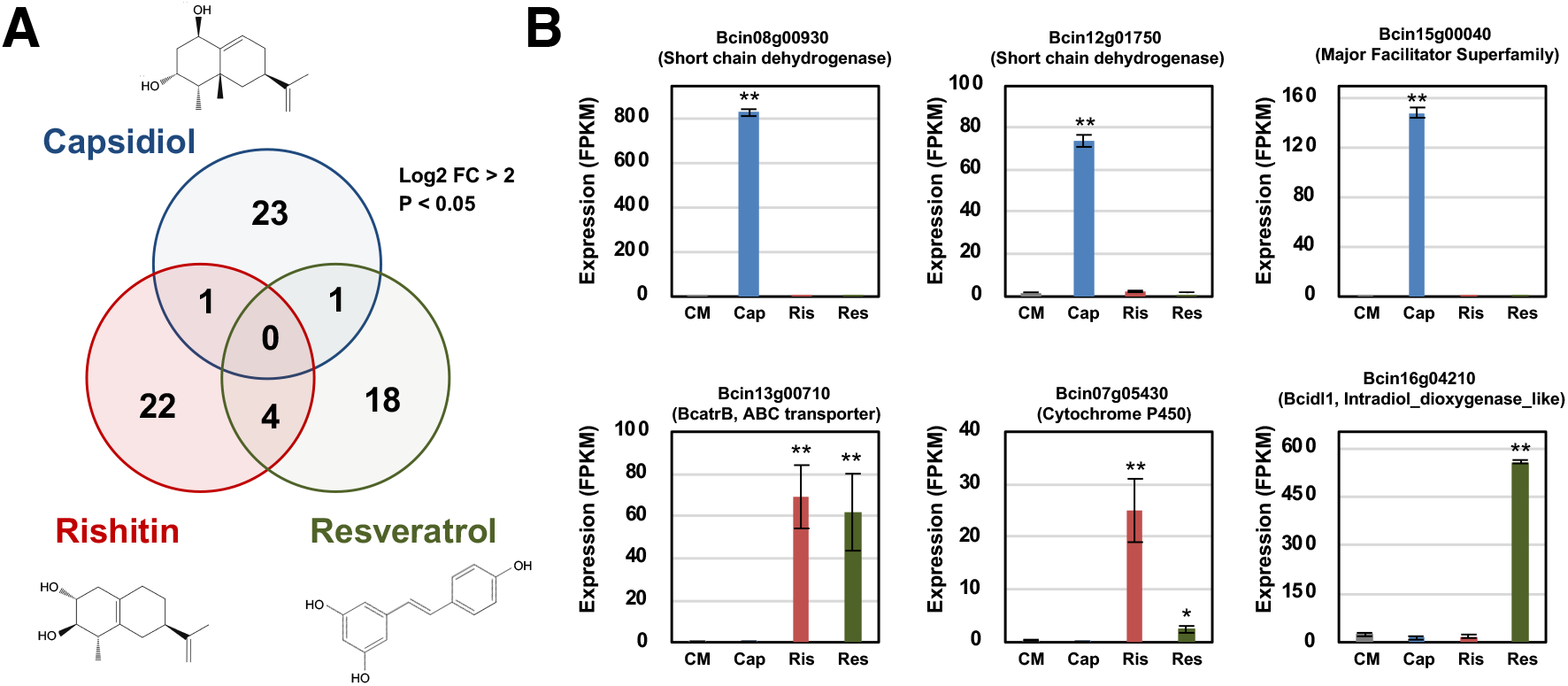
Unique set of genes are upregulated in *Botrytis cinerea* treated with capsidiol, rishitin and resveratrol. **(A)** Venn diagram showing genes upregulated in *B. cinerea* cultured in CM media containing 100 µM capsidiol, 500 µM rishitin or 500 µM resveratrol for 24 h. The numbers of significantly upregulated genes by phytoalexin treatment with Log2 FC > 2 compared with control (CM without phytoalexin) and P value < 0.05 are presented. **(B)** Expression profiles of representative genes upregulated by the treatment with capsidiol, rishitin or resveratrol. The gene expression (FPKM value) was determined by RNA-seq analysis of *B. cinerea* cultured in CM media containing 100 µM capsidiol, 500 µM rishitin or 500 µM resveratrol for 24 h. Data are mean ± SE (n = 3). Asterisks indicate a significant difference from the control (CM) as assessed by two-tailed Student’s *t*-test, **P < 0.01, *P < 0.05.

### Bcin08g00930 encodes a short-chain dehydrogenase for the detoxification of capsidiol

To investigate the function of SDR genes induced by capsidiol treatment, an endophytic fungus *Epichloë festucae* was employed for the heterologous expression of these genes. Bcin08g00930 and Bcin12g01750 were expressed in *E. festucae* under the control of the constitutive *TEF* promoter (28). The growth of wild type and control *E. festucae* expressing DsRed was severely inhibited in the 100 µM capsidiol (Fig. 3A and B). *E. festucae* transformants expressing Bcin12g01750 also hardly grew in 100 µM capsidiol, however, *E. festucae* became tolerant to capsidiol by the expression of Bcin08g00930 (Fig. 3A and B). The amount of capsidiol was not altered by wild type, DsRed or Bcin12g01750 expressing *E. festucae*, while capsidiol was metabolized to capsenone by the *E. festucae* transformant expressing Bcin08g00930 (Fig. 3C). These results indicated that Bcin08g00930 encodes a dehydrogenase that can detoxify capsidiol, thus the SDR encoded by Bcin08g00930 was designated as BcCPDH standing for *B. cinerea* capsidiol dehydrogenase.

**Fig. 3.**
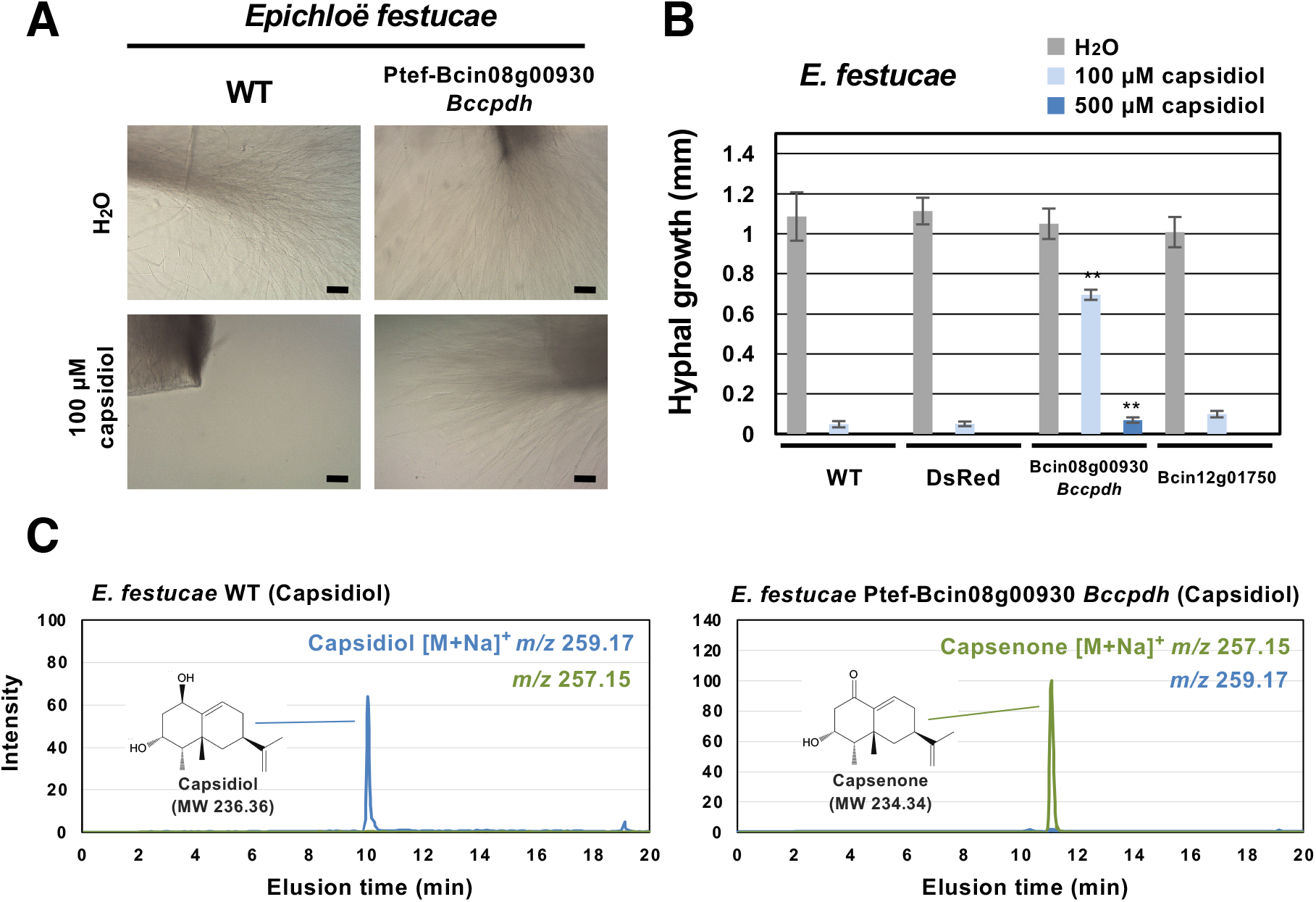
Bcin08g00930 encodes a capsidiol detoxification enzyme, capsidiol dehydrogenase BcCPDH. **(A)** Mycelia of *Epichloë festucae* wild type (WT) or a transformant expressing Bcin08g00930 were incubated in water or 100 µM capsidiol and outgrowth of mycelia was observed 7 days after inoculation. Bars = 100 µm. **(B)** Hyphal outgrowth of *E. festucae* WT, transformants expressing DsRed, Bcin08g00930 (BcCPDH) or Bcin12g01750 in water, 100 µM or 500 µM capsidiol was measured after 24 h of incubation. Data are mean ± SE (n = 6). Asterisks indicate a significant difference from WT as assessed by two-tailed Student’s *t*-test, **P < 0.01. **(C)** Mycelia of *E. festucae* WT or transformant expressing Bcin08g00930 (*Bccpdh*) were incubated in 100 µM capsidiol or for 48 h. Capsidiol and capsenone were detected by LC/MS.

### Expression of *Bccpdh* Is specifically induced by capsidiol

To further investigate the expression pattern of *Bccpdh, B. cinerea* was transformed with a reporter construct for GFP expression under the control of the 1 kb proximal *Bccpdh* promoter (P_*Bccpdh:GFP*). P_*Bccpdh:GFP* transformants showed no obvious expression of GFP in water, but a significant increase of GFP fluorescence was detected in 500 µM capsidiol (Fig. 4A). The P_*Bccpdh:GFP* transformant was incubated with other anti-microbial terpenoids produced by Solanaceae species, including rishitin, debneyol (29), sclareol (30) and capsidiol 3-acetate (31). Among the terpenoids tested, treatment of capsidiol and capsidiol 3-acetate induced expression of GFP in the P_*Bccpdh:GFP* transformant. This result further indicated that *B. cinerea* specifically reacts to capsidiol and its derivative capsidiol 3-acetate. As the *Bccpdh* promoter was activated by capsidiol 3-acetate, we investigated whether BcCPDH could metabolize capsidiol 3-acetate to capsenone 3-acetate. However, capsidiol 3-acetate was not metabolized by *E. festucae* expressing *Bccpdh* (Bcin08g00930) (Supplemental Fig. S7A). Consistently, capsidiol 3-acetate was directly oxidized by *B. cinerea* and at least two different forms of oxidized capsidiol 3-acetate, but not capsenone 3-acetate, were detected (Supplemental Fig. S7B), indicating that capsidiol and capsidiol 3-acetate were metabolized in *B. cinerea* by distinctive pathways.

**Fig. 4.**
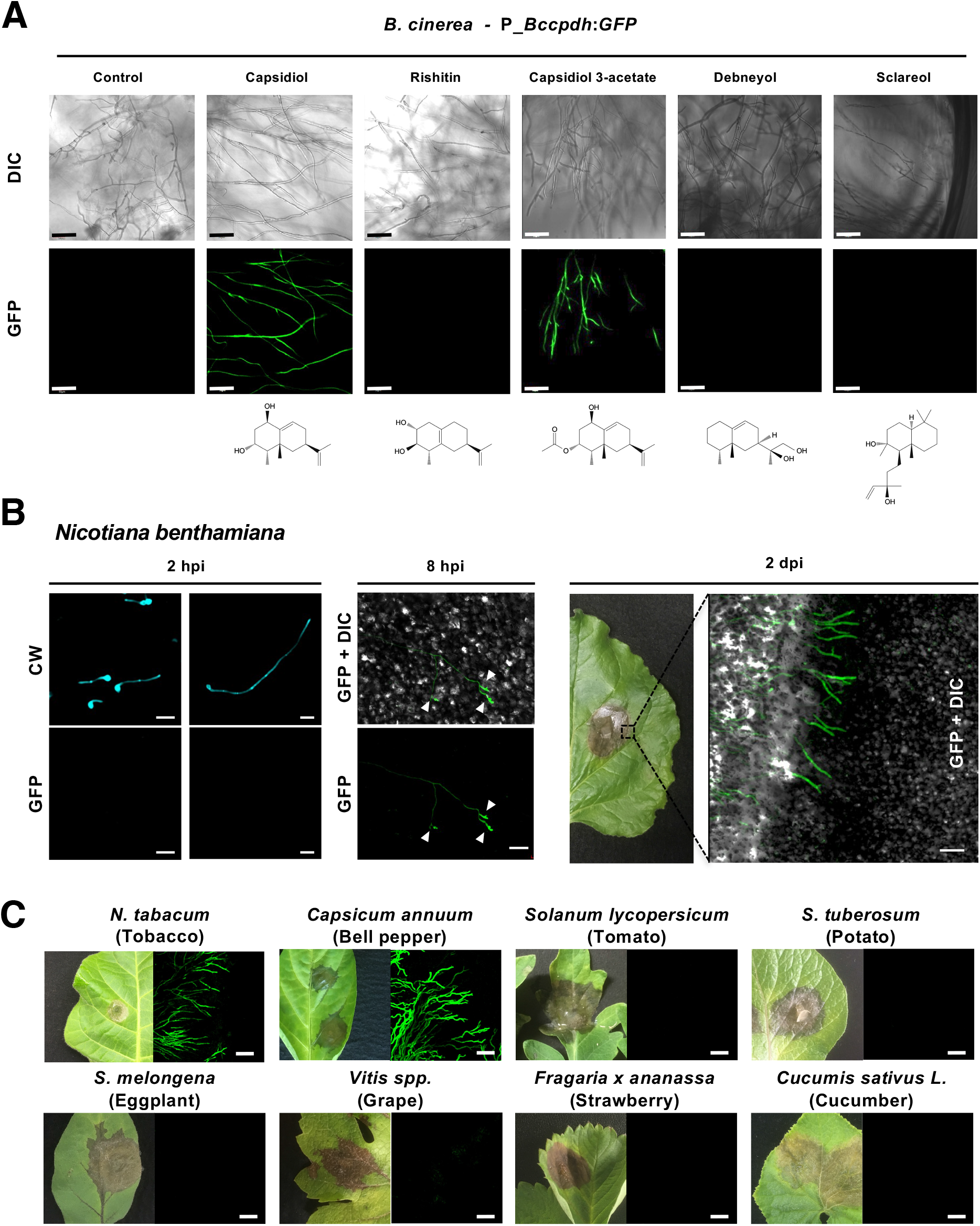
Specific activation of the *B. cinerea Bccpdh* promoter by capsidiol and its derivative. **(A)** Mycelia of *B. cinerea* transformant containing *GFP* gene under the control of 1 kb *Bccpdh* promoter (P_*Bccpdh:GFP*) was incubated in CM media containing 500 µM of anti-microbial terpenoids. Bars = 50 µm. **(B)** (left and middle) Leaves of *N. benthamiana* were inoculated with conidia of *B. cinerea* P_*Bccpdh:GFP* transformant. Expression of GFP in germinating conidia on the leaf surface was observed by confocal laser microscopy 2 or 8 h after inoculation (hpi). CW, stained with calcofluor white. Arrowheads indicate the appressoria of *B. cinerea*. Bars = 20 µm. (right) Leaves of *N. benthamiana* were inoculated with mycelia of *B. cinerea* P_*Bccpdh:GFP* transformant and the edge of the lesion was observed by confocal laser microscopy 2 d after the inoculation (2 dpi). Bar = 100 µm. **(C)** Leaves of indicated plant were inoculated with mycelia of *B. cinerea* P_*Bccpdh:GFP* transformant and the edge of the lesion was observed by confocal laser microscopy 2 d after the inoculation. Bars = 100 µm.

Analysis of different lengths of *Bccpdh* promoters indicated that 250 bp of promoter sequence are sufficient for specific activation of the promoter by capsidiol treatment, and the cis-element required for capsidiol-specific expression was shown to be within -250 and -200 bp upstream of the start codon (Supplemental Figs. S8 and 9). By further analysis using *B. cinerea* transformant P_*Bccpdh:Luc* using luciferase as a quantitative marker, it was shown that *Bccpdh* promoter was activated within 2 h of capsidiol treatment, and the expression activity decreased as capsidiol was metabolized (Supplemental Figs. S10, Supplementary Notes 3).

### Expression of *Bccpdh* is specifically induced during the infection of capsidiol producing plant species

Leaves of *Nicotiana benthamiana*, which produce capsidiol as the major phytoalexin (13), were inoculated with conidia of the *B. cinerea* transformant P_*Bccpdh:GFP*. No GFP signal was detected in germinating *B. cinerea* conidia grown on the surface of *N. benthamiana* leaves 2 h after the inoculation, but obvious GFP expression was detected in the appressoria at 8 h after the inoculation (Fig. 4B). Growing hyphae in the leaf tissue of *N. benthamiana* showed GFP fluorescence, and the intensity of GFP fluorescence in hyphae was stronger near the edge of the lesion compared with that of hyphae growing in the areas of dead tissue within the lesions (Fig. 4B), probably because capsidiol was eventually detoxified in these areas, which are heavily infected with *B. cinerea*.

To examine the possibility whether polyxenous *B. cinerea* activates *Bccpdh* for detoxification of other phytoalexin or anti-microbial compounds produced in other plant species, the *B. cinerea* P_*Bccpdh:GFP* transformant was inoculated on a wide variety of plants. Over 50 plant species were used for the inoculation test including 9 Solanaceae, 6 Brassicaceae, 6 Rosaceae, 5 Fabaceae and 6 Asteraceae plants, and development of disease symptoms was observed in all tested plant species. Among the tested plants, expression of GFP under the control of *Bccpdh* promoter was only detected during the infection in three *Nicotiana* and two *Capsicum* species (Fig. 4C, Supplemental Fig. S11 and Table S4), all of which are reported to produce capsidiol (8, 32–34). These results further indicated that *B. cinerea* specifically recognizes capsidiol for the induction of *Bccpdh*.

### BcCPDH metabolizes and detoxifies capsidiol to capsenone in *B. cinerea*

*B. cinerea Bccpdh* KO mutant (*Δbccpdh*) was produced to investigate the function of BcCPDH (Supplemental Fig. S12). Mycelia of *B. cinerea* wild type and *Δbccpdh* were incubated with capsidiol and the metabolites were detected by LC/MS. While capsidiol was metabolized to capsenone in the wild type strain, most of the capsidiol remained unmetabolized two days after incubation with the *Δbccpdh* strain (Fig. 5A). Instead, oxidized capsidiol, which was not detectable in the wild type strain, was detected in *Δbccpdh* incubations (Supplemental Fig. S13 and Supplementary Notes 4). While the growth of *B. cinerea* hyphae was not affected by the disruption of the *Bccpdh* gene in 100 µM capsidiol, growth of *Δbccpdh* was diminished compared with that of wild type *B. cinerea* at higher concentrations of capsidiol (Fig. 5B). These results confirmed that BcCPDH is the enzyme responsible for the detoxification of capsidiol in *B. cinerea*.

**Fig. 5.**
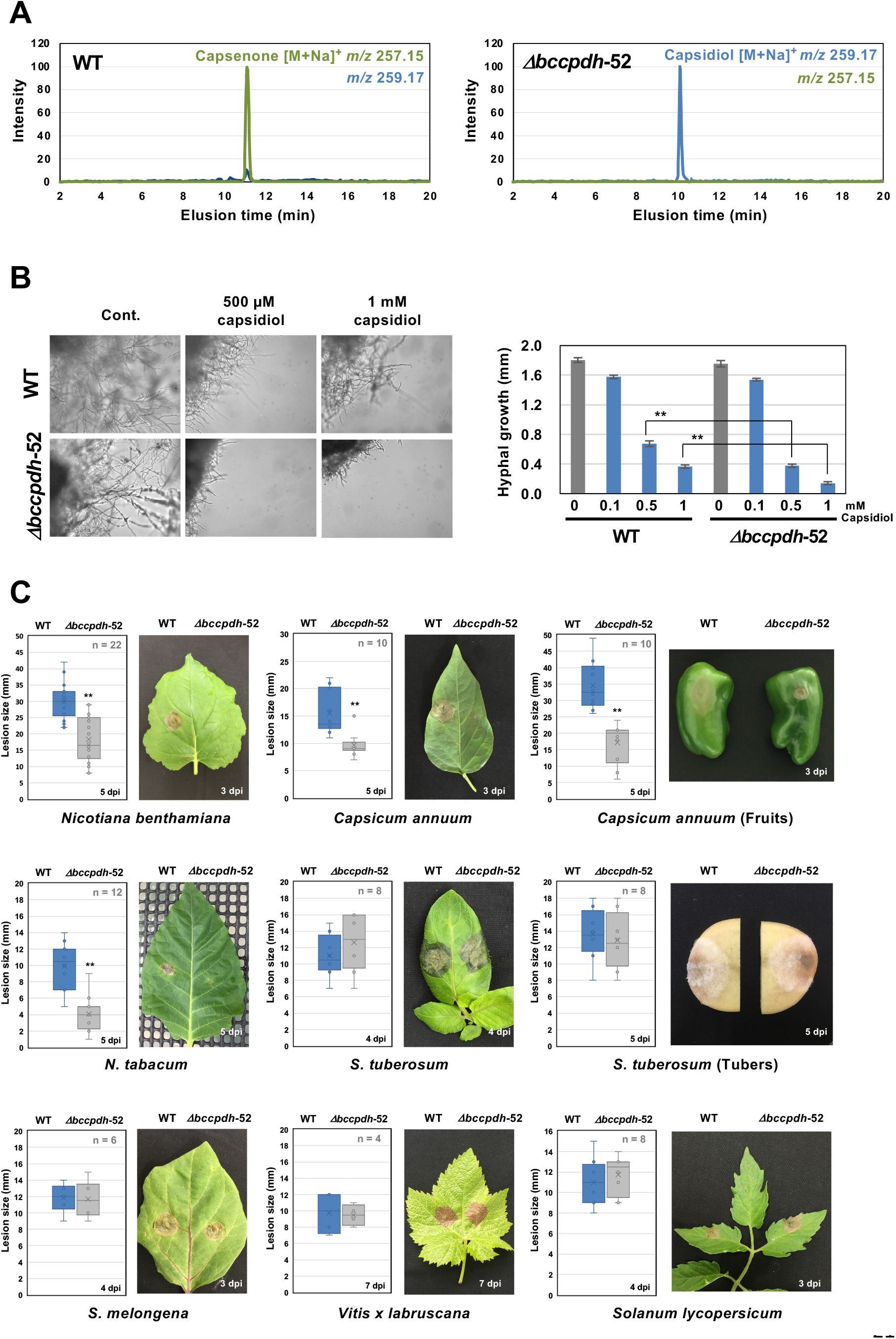
*B. cinerea* BcCPDH is essential for the detoxification of capsidiol and virulence in the plant species producing capsidiol. **(A)** Mycelial block (approx. 1 mm^3^) of *B. cinerea* wild type (WT) or *Bccpdh* KO mutant strain (*Δbccpdh*-52) was incubated in 100 µM capsidiol for 4 d. Capsenone and capsidiol was detected by LC/MS. **(B)** Mycelial blocks of *B. cinerea* WT or *Δbccpdh*-52 were incubated in capsidiol and outgrowth of hyphae was measured after 36 h of incubation. Data are mean ± SE (n = 10). Asterisks indicate a significant difference from WT as assessed by two-tailed Student’s *t*-test, **P < 0.01. **(C)** Leaves, tubers or fruits of indicated plant were inoculated with mycelial block (5 mm^3^) of *B. cinerea* WT or *Δbccpdh*-52 and lesion size was measured between 4 and 7 days after inoculation (dpi). Asterisks indicate a significant difference from WT as assessed by two-tailed Student’s *t*-test. **P < 0.01. Lines and crosses (x) in the columns indicate the median and mean values, respectively.

### Epoxidation of capsidiol and capsenone is mediated via Bcin16g01490

The incubation of the *Δbccpdh* knockout strain in capsidiol revealed the presence of a potential secondary capsidiol degradation pathway, which resulted in the accumulation of oxidized capsidiol. Since we were unable to find any indication for such a pathway in our RNAseq results for 100 µM capsidiol treated incubations, we extended our search to a preliminary RNAseq analysis which used 500 µM capsidiol treatments. We identified the gene Bcin16g01490 encoding a cytochrome P450, which was significantly upregulated under these elevated capsidiol concentrations (Supplemental Table S1 and Fig. S14A). Heterologous expression of Bcin16g01490 in *E. festucae*, resulted in the conversion of capsidiol to oxidized capsidiol as detected during *Δbccpdh* strain incubations (Supplemental Figs. S13 and 14, Supplementary Notes 4). Moreover, sequential incubation of capsidiol with the *Bccpdh* expressing *E. festucae* transformant followed by a Bcin16g01490 expressing transformant reproduced the reactions detected in *B. cinerea* (Supplemental Figs. S4 and S14B). Structural analysis of oxidized capsenone indicated that the end product of these reactions is capsenone 11,12-epoxide (Fig. 1 and Supplemental Fig. S15-17, Supplementary Notes 5).

### BcCPDH is required for the pathogenicity of *B. cinerea* on plant species producing capsidiol

Pathogenicity of *B. cinerea Δbccpdh* transformants was tested for several plant species. On plant species that produce capsidiol, such as *N. benthamiana, N. tabacum* and *C. annuum*, the development of disease symptoms by *Δbccpdh* was significantly reduced compared with those caused by wild type *B. cinerea*. In contrast, both wild type and *Δbccpdh* strains caused comparable symptoms in potato, tomato, grape and eggplant, which is consistent with the finding that expression of *Bccpdh* is not induced during the infection in these plant species (Fig. 5C, Supplemental Fig. S18). In the complementation strain, the ability to metabolize capsidiol to capsenone was restored, as was virulence against capsidiol producing plants (Supplemental Fig. S19). These results indicated that BcCPDH is a dedicated enzyme for the virulence of *B. cinerea* on capsidiol-producing plant species.

### Distribution of *Bccpdh* homologues in the fungal kingdom

The distribution of *Bccpdh* homologs in taxonomically related fungal species (Ascomycota, Leotiomycetes) was investigated. A search for *Bccpdh* homologs in the genome sequences of 6 host-specialized phytopathogenic *Botrytis* species, such as *B. tulipae* (pathogen of tulip), *B. hyacinthi* (hyacinth) and *B. paeoniae* (peony), indicated that among *Botrytis* species, *Bccpdh* is a unique gene only found in *B. cinerea* (Supplemental Fig. S20). Consistently, 4 *Botrytis* species, not including *B. cinerea*, were incubated with capsidiol, but no CPDH activity was detected for the tested strains (Supplemental Fig. S21). Search in 11 draft genomes of Leotiomycete fungi also indicated the absence of *Bccpdh* homologs in these species. Comparison of the corresponding genome regions between *Botrytis* species and *S. sclerotiorum* (another polyxenous pathogen belonging to Leotiomycetes) revealed that the approx. 4.9 kb sequence surrounding *Bccpdh* is unique to *B. cinerea* (Fig. 6). This unique sequence shows a lower GC content compared to the surrounding region (Supplemental Fig. S22), suggesting that *Bccpdh* might have been obtained via horizontal gene transfer (35). A blast search using BcCPDH as query sequence revealed that probable orthologs can be found only in some Pezizomycotina fungi belonging to Ascomycota (Supplemental Fig. S20). Orthologs were found from a taxonomically diverse range of fungal species, including animal and insect pathogens, and there was no correlation between their homology and taxonomic relationship, which might indicate multiple horizontal gene transfer events of the *cpdh* gene in the diversification of Ascomycota fungi (Supplemental Figs. 20, 23-25, Supplementary Notes 6). Among plant pathogenic fungi, *Bccpdh* homologs were found in *Fusarium* species, all of which are pathogens that infect plants that do not produce capsidiol. This result is consistent with a previous report showing that some *Fusarium* species can metabolize capsidiol to capsenone (36).

**Fig. 6.**
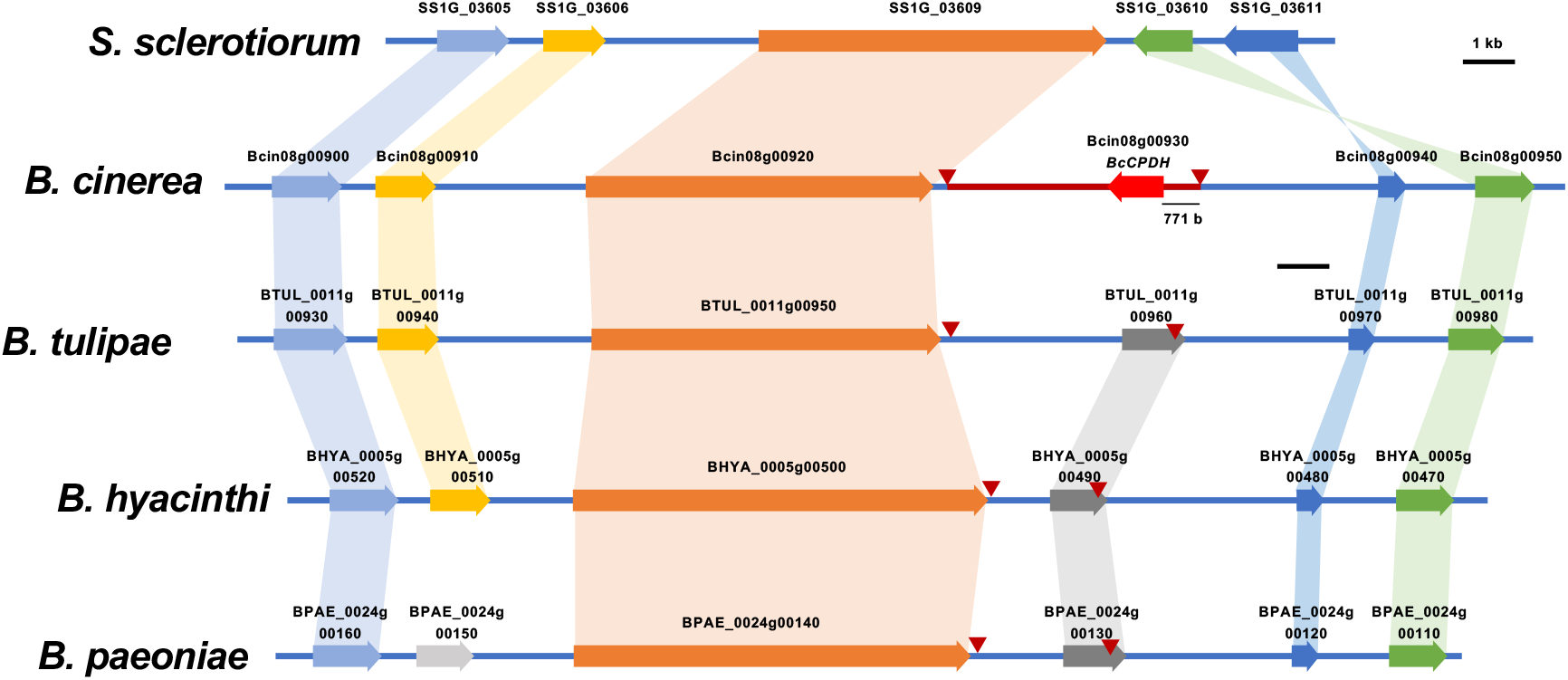
Comparison of the *B. cinerea Bccpdh* gene locus with corresponding genome region of other *Botrytis* species and *Sclerotinia sclerotiorum*. Edge of conserved region among *Botrytis species* and specific region for *B. cinerea* (red line) were indicated by red arrowheads.

### CPDH activity is conserved among *Botrytis cinerea* strains

Although *B. cinerea* is a polyxenous fungus, it has been reported that different strains of *B. cinerea* exhibit various degrees of virulence on different host plants (37, 38). Hence, we investigated whether BcCPDH is conserved among strains isolated from diverse plant species. Twenty-four *B. cinerea* strains isolated from 14 different plant species were incubated with capsidiol and resultant capsenone was detected by GC/MS. All tested *B. cinerea* strains can metabolize capsidiol to capsenone, indicating that BcCPDH is highly conserved among *B. cinerea* strains even though their (most recent) host was not a producer of capsidiol (Fig. 7A, Supplemental Table S5). Similarly, 17 strains of *F. oxysporum* isolated from 7 plant species were subjected to the analysis for CPDH activity. Eight out of 17 *F. oxysporum* strains showed CPDH activity, and the activity was not related to the natural host of the strains (Fig. 7B, Supplemental Table S6). This result is consistent with the finding that *Bccpdh* orthologues can be found in 5 out of 14 available genomes of *F. oxysporum* (Supplemental Fig. S25, data not shown). These results indicate that the cpdh gene is randomly distributed in *F. oxysporum* strains, whereas it is highly conserved among *B. cinerea* strains.

**Fig. 7.**
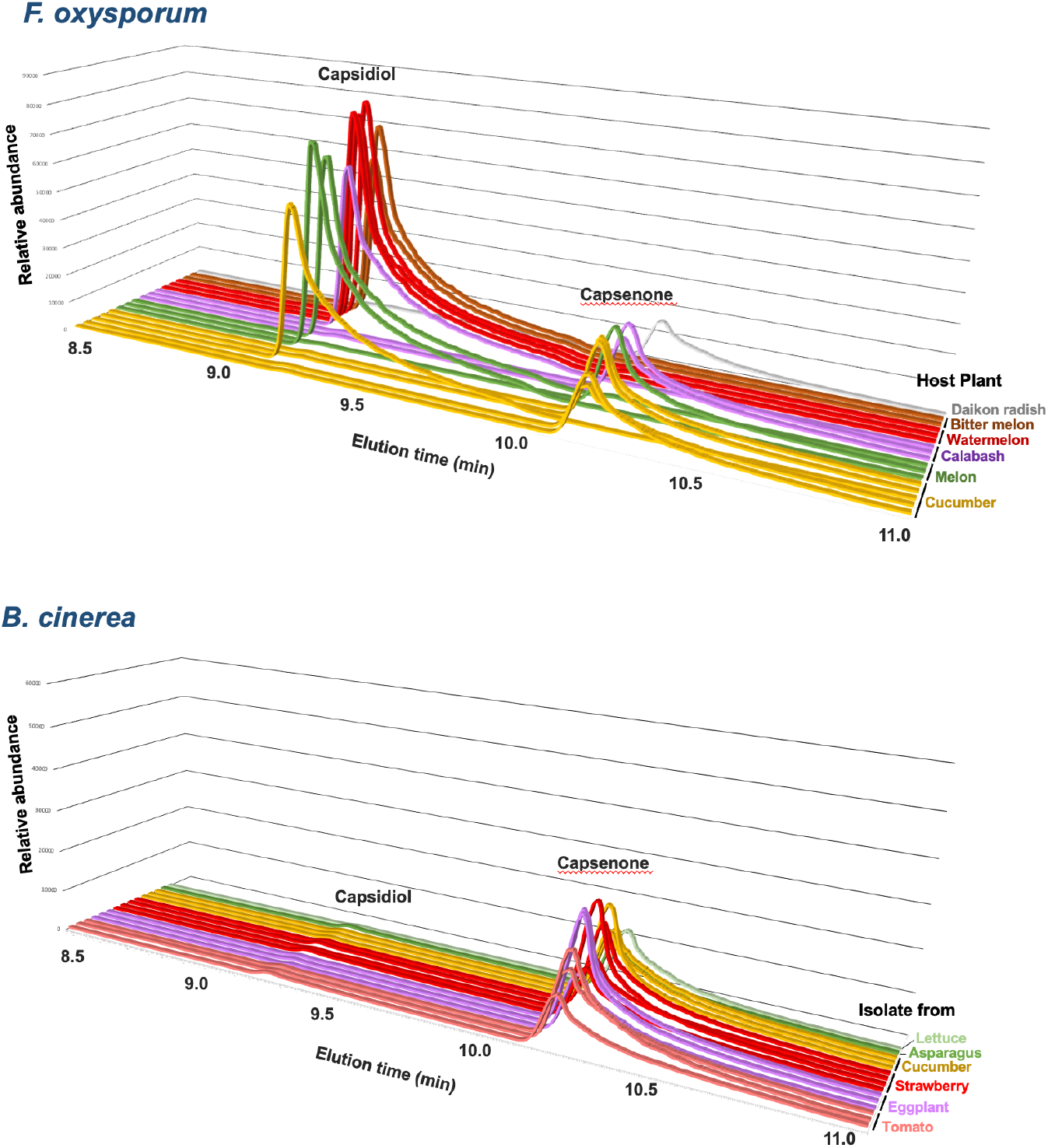
CPDH activity in *F. oxysporum* and *B. cinerea* strains isolated form a variety of plants. CPDH activity in *F. oxysporum* and *B. cinerea* strains isolated form a variety of plants. Mycelial blocks (approx. 1 mm^3^) were incubated in 500 µM capsidiol for 2 days and capsidiol or capsenone were detected by GC/MS. Each elution profile describes the capsidiol and capsenone content in the solution after incubation of the strain with capsidiol. Strains with a substantial peak for capsidiol indicate the absence of CPDH activity.

## Discussion

To survive the exposure to microorganisms in the environment, different plant species have developed diverse resistance mechanisms over the course of evolution. The structural variety of phytoalexins is a prime example of such diversity: different plant families produce phytoalexins with relatively similar basic structures, but their side-chain structures often differ from species to species (4). These differences in structure between species may have contributed to the determination of host specificity between plants and pathogens, as in some cases a pathogen may have acquired resistance to a particular phytoalexin, but an analogous substance is not overcome by the same resistance mechanism (39). However, plant pathogens with a broad host range such as *B. cinerea* must employ strategies to counter a multitude of diverse phytoalexins. The prompt killing of plant cells, or the presence of efflux pumps of extremely low specificity may present effective strategies for such pathogens. Indeed, *B. cinerea* produces host-nonspecific phytotoxins (such as botrydial and botcinic acid, 40, 41) upon infection, and activates transporters capable of effluxing diverse substances such as PDR-type ABC transporter BcatrB and MFS transporter Bcmfs1 (26, 27, 42). More recently, it has been shown that *B. cinerea* suppresses the immune response of different plant species via gene silencing by delivering various small RNAs into host cells (43, 44).

In addition to effective infection mechanisms which generally facilitate infection of a wide range of plants, this study revealed that *B. cinerea* responds to chemical cues from the host plant to adapt its infection strategy to overcome specific plant resistance mechanisms. *B. cinerea* strictly distinguishes structurally similar phytoalexins, such as capsidiol and rishitin, and activates appropriate detoxification responses. Interestingly, treatment of *B. cinerea* with phytoalexins activated not only genes involved in detoxification and efflux of toxic compounds, but also genes predicted to be involved in pathogenicity to plants. These results support the notion that *B. cinerea* utilizes phytoalexins as a means to identify a given host and adjust the method of infection.

Expression of the *Bccpdh* gene is activated specifically during infection of capsidiol-producing plants, and the pathogenicity of the *bccpdh* mutant strains is compromised on plants producing capsidiol, but does not suffer any disadvantages on plants that do not produce capsidiol. These results suggest that BcCPDH is a critical component that specifically enables *B. cinerea* to infect capsidiol producing plants. Despite capsidiol producing plants representing only a small fraction of the over 400 host plants of *B. cinerea*, CPDH activity was maintained in all investigated *B. cinerea* strains isolated from plants that do not produce capsidiol. This may hint at the presence of a selection pressure against the loss of CPDH, despite it only affecting a limited number of host plants. This poses the question of whether *B. cinerea* is able to maintain acquired resistances for a prolonged time, which may explain how it evolved and establish itself as the polyxenous pathogen that it is.

### How does *B. cinerea* recognize phytoalexins with similar structures?

Although *B. cinerea* is known to have the ability to metabolize a wide variety of phytoalexins (18), only a small portion of these detoxification mechanisms are required for the infection of any particular plant. Thus, *B. cinerea* needs to strictly control those various detoxification mechanisms, since maintaining sufficient levels of all these detoxification mechanisms represents a considerable waste of resources. Elucidating the mechanism by which *B. cinerea* recognizes phytoalexins and activates a specific set of genes is one of the important subjects for further research. Two major scenarios can be envisioned: first, *B. cinerea* may possess receptors that can identify the chemical structures of phytoalexins. In plants, various terpenes with diverse structures are employed as hormones recognized by specific receptors, such as gibberellins, cytokinins, and abscisic acid (39). However, the possibility that *B. cinerea* maintains receptors to perceive all of the myriad of phytoalexins may not be realistic. The second scenario would be that *B. cinerea* recognizes the damage caused by phytoalexins. However, as for capsidiol and rishitin, both are presumed to have cell membranes as their targets, and these phytoalexins are relatively unspecific toxicants that cause damage even to plant cells (5, 13, 17, 45), so it is not certain whether there exists a specific target by which these two compounds can be distinguished. Given that some genes, such as *BcatrB* encoding a multidrug resistance pump, are commonly induced by structurally unrelated phytoalexins and fungicides treatments (26, 27), *B. cinerea* may possess both induction mechanisms. Using the reporter system developed in this study, mutant strains defective in capsidiol response could be isolated to elucidate the mechanism by which *B. cinerea* distinctly identifies phytoalexins.

### Does *B. cinerea* possess other capsidiol resistance mechanism besides detoxification by BcCPDH?

Although the *bccpdh* knockout mutants showed reduced virulence on plant species producing capsidiol, the mutant can develop the disease symptom on these plants and showed tolerance to 100 µM capsidiol, same as the wild type. In contrast, *Phytophthora* spp., *Alternaria* spp. and *E. festucae* are sensitive to the same concentration of capsidiol. Although we also isolated Bcin16g01490, which we found to oxidize capsidiol, its activity is fairly limited, indicating that it is unlikely to confer sufficient protection against capsidiol. We therefore presumed that *B. cinerea* has another mechanism for capsidiol tolerance other than detoxification. RNAseq analysis of *B. cinerea* genes upregulated by capsidiol treatment identified 2 genes (Bcin15g00040.1 and Bcin14g02870.1/Bcmfs1) encoding MFS transporters and a gene (Bcin01g05890.1/Bcbmr1) encoding an ABC transporter (Supplemental Table S1). Bcmfs1 has been reported to be involved in the resistance of *B. cinerea* to structurally different natural toxicants (camptothecin produced by the plant *Camptotheca acuminata* and cercosporin produced by the plant pathogenic fungus *Cercospora kikuchii*) and fungicides (sterol demethylation inhibitors, DMIs) (42). The *bcbmr1* mutants showed an increased sensitivity to fungicides, polyoxin (a chitin synthetase inhibitor) and iprobenfos (a choline biosynthesis inhibitor) (46). The expression of one or more of these transporters may be involved in capsidiol efflux from *B. cinerea* cells, and regulated by signaling that is common to *Bccpdh*.

### Why and how can *B. cinerea* strains stably maintain BcCPDH?

CPDH activity was detected in all tested *B. cinerea* strains isolated from plants that do not produce capsidiol, and the *Bccpdh* gene is conserved in the genomes of published *B. cinerea* strains. In *F. oxysporum*, in contrast, there were strains with and without CPDH activity regardless of the host plant, and consistently, some publicly available *F. oxysporum* genomes contain *cpdh* homologs and others do not. As shown in this study, metabolic capacity (and tolerance) to capsidiol and rishitin can be detected in some fungal strains regardless of the natural host of these pathogens, and perhaps unused detoxification activities are maintained in the population of phytopathogenic filamentous fungi for future use.

It is theoretically implausible that all *B. cinerea* strains maintain an enzyme required only upon infection of capsidiol-producing plants, even if said gene is completely repressed in its expression under normal circumstances. Given that *B. cinerea* has an extremely wide host range, some strains will go for long periods where capsidiol detoxification ability is not a selection pressure. One possibility is that BcCPDH has functions other than capsidiol degradation. For example, BcCPDH might be involved in the detoxification of antimicrobial substances produced by insects, because it has been reported that spores of *B. cinerea* are transmitted between plants via insects such as thrips (47). Since homologous genes for *Bccpdh* are found in several insect-infecting fungi, it is expected that CPDH-like enzymes in these species can metabolize insect-derived substances. We, therefore, performed preliminary experiments to determine whether *Bccpdh* expression can be induced by inoculation of *B. cinerea* P_*Bccpdh:GFP* transformant with several insects, but to date, no induction of GFP expression has been detected (data not shown). Alternatively, *B. cinerea* may have acquired the trait of having many hosts because of its ability to maintain rarely used virulence factors. Future clarification of the phytoalexin recognition mechanisms and comparative analysis of the genome with that of closely related *Botrytis* species are anticipated to elucidate unknown features of *B. cinerea* that have led to its evolution as a polyxenous fungi.

## Materials and Methods

Comprehensive descriptions of the materials and methods used in this study, including, biological materials, vector construction, transformation of *E. festucae* and *B. cinerea*, confocal microscopy, pathogenicity tests and detection and structural analysis of phytoalexins and their derivatives, are available in Supplemental data.

## Data Availability

RNA-seq data reported in this work are available in GenBank under the accession number DRA013980.

## Acknowledgments

We thank Prof. Barry Scott (Massey University, New Zealand) for providing *E. festucae* strain Fl1 and critical reading of the manuscript. We also thank Dr. David Jones (The Australian National University, Australia) for *N. benthamiana* seeds, Ms. Kayo Shirai (Hokkaido Central Agricultural Experiment Station, Japan), and Dr. Seishi Akino (Hokkaido University, Japan) for providing *P. infestans* isolate 08YD1, Prof. Takashi Tsuge (Chubu University, Japan) for providing *Fusarium* strains, Dr. Haruhisa Suga for providing *Fusarium graminearum* strain, Mr. Masashi Matsusaki (Aichi Prefectural Agricultural Research Center, Japan) for providing *Fusarium oxysporum* f. sp. *lycopersici* strain, and Mr. Taku Kawakami (Mie Prefecture Agricultural Research Institute, Japan) for providing *B. cinerea* strains. We are also grateful to Prof. Kazuhito Kawakita for valuable suggestions, and Dr. Kenji Asano and Mr. Seiji Tamiya (National Agricultural Research Center for Hokkaido Region, Japan) and Mr. Yasuki Tahara (Nagoya University, Japan) for providing tubers of potato cultivars.

