## Supplementary Figs and Tables for "*Botrytis cinerea* identifies host plants via the recognition of antifungal capsidiol to induce expression of a specific detoxification gene"


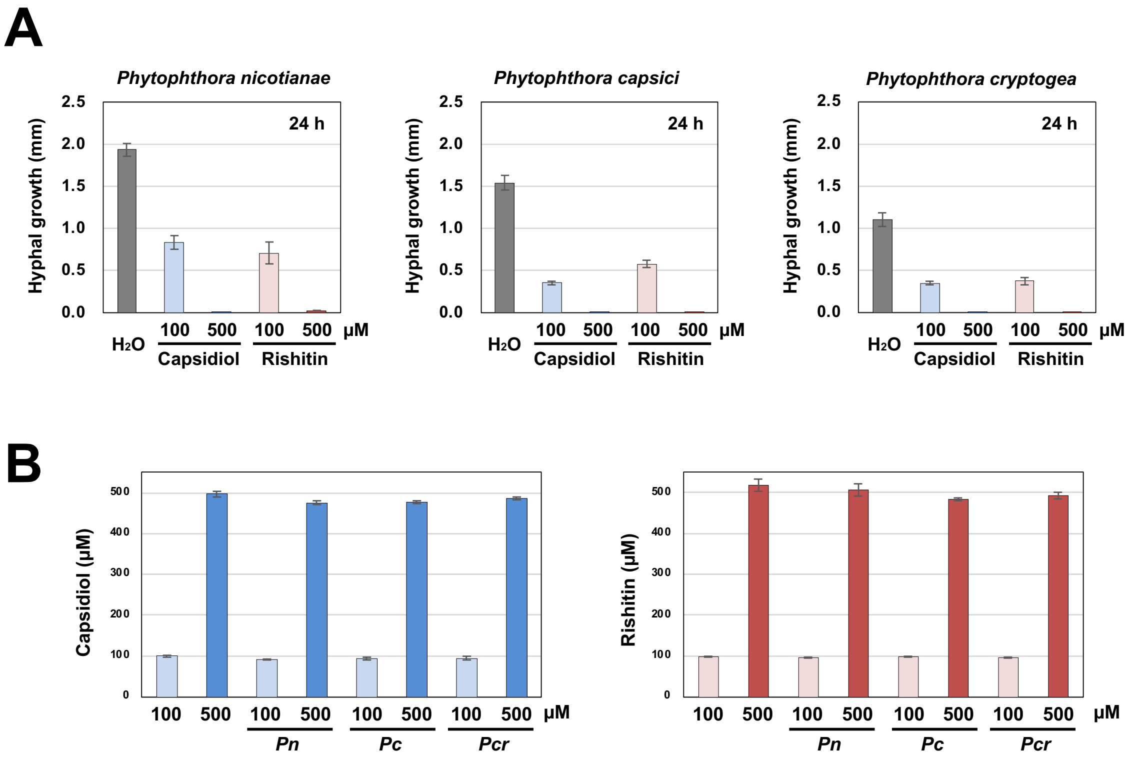


**Fig. S1.** Sensitivity and metabolic capacity of sesquiterpenoid phytoalexins in oomycete pathogens isolated from Solanaceae plants. **(A)** Mycelial blocks (approx. 1 $mm^3$) of the indicated pathogen were incubated in 50 μl water, 100 or 500 μM capsidiol or rishitin. Outgrowth of hyphae from the mycelial block was measured after 24 h of incubation (n = 6). **(B)** Residual capsidiol and rishitin was quantified after 48 h of incubation (n = 3). *Pn*, *Phytophthora nicatianae* (strain Pn96 isolated from tobacco); *Pc*, *P. capsici* (strain CH01CMP1 isolated from green pepper); *Pcr*, *P. cryptogea* (strain CH88-18 isolated from nipplefruit).

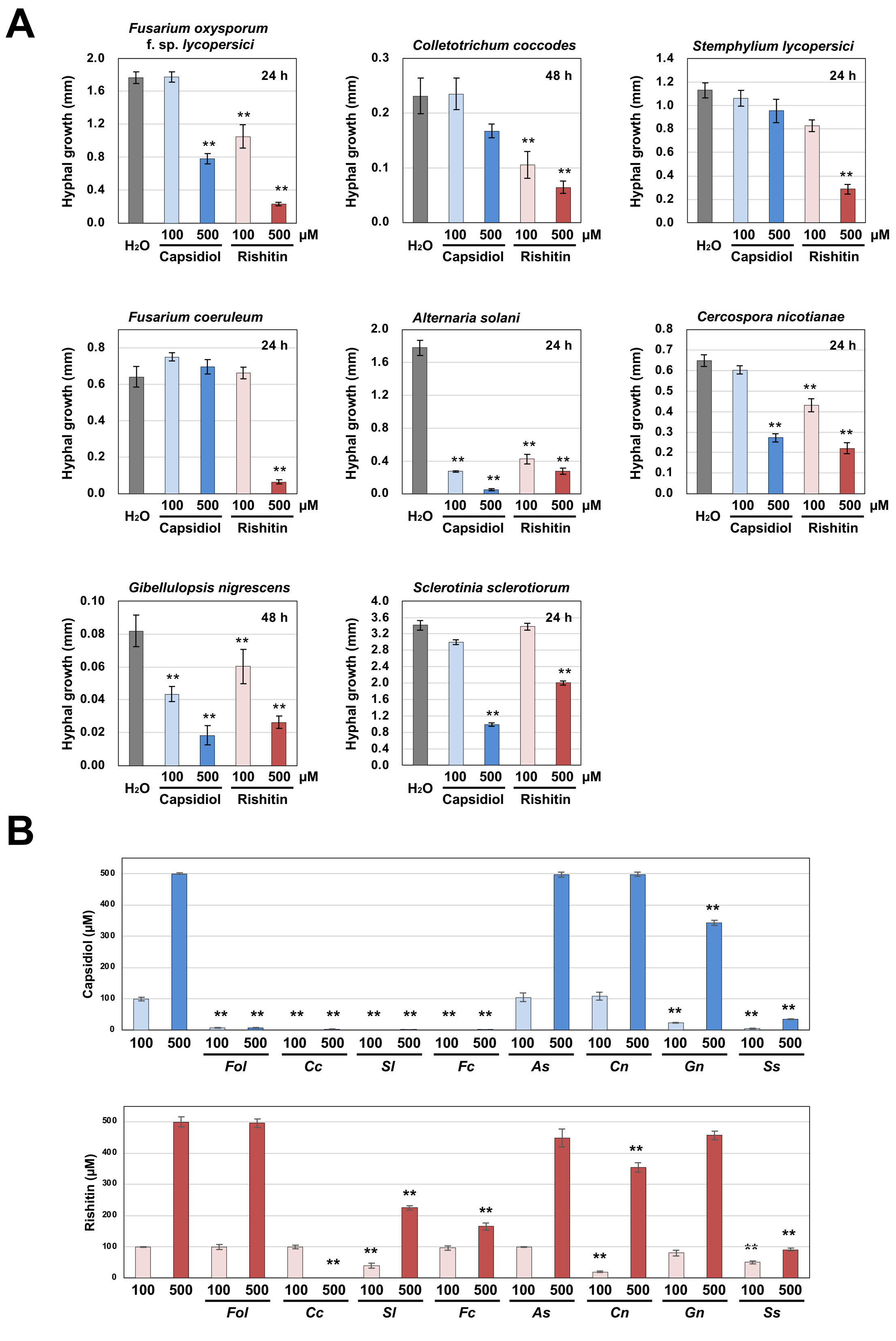


**Fig. S2.** Sensitivity and metabolic capacity of sesquiterpenoid phytoalexins in fungal pathogens isolated from Solanaceae plants. **(A)** Mycelial blocks (approx. 1 $mm^3$) of the indicated pathogen were incubated in 50 µl water, 100 µM or 500 µM capsidiol or rishitin. Outgrowth of hyphae from the mycelial block was measured after 24 h or 48 h of incubation (n = 6). **(B)** Residual capsidiol and rishitin was quantified by GC/MS after 48 h of incubation. Data marked with asterisks are significantly different from control as assessed by the two-tailed Student's *t*-test: $**P < 0.01$. *Fol*, *Fusarium oxysporum* f. sp. *lycopersici* (strain 9855-1 isolated from tomato); *Cc*, *Colletotrichum coccodes* (strain 9855-1 isolated from potato); *Sl*, *Stemphylium lycopersici* (strain KuNBY1 isolated from tobacco); Fc, *F. coeruleum* (strain K. Kita 37 isolated from potato); As, *Alternaria solani* (KL1 isolated from potato); *Cn*, *Cercospora nicotianae* (strain CTC5 isolated from tobacco); *Gn*, *Gibellulopsis nigrescens* (strain Kita44 isolated from potato); *Ss*, *Sclerotinia sclerotiorum* (isolate SU-1 isolated from eggplant).

### Supplementary Note 1

#### Several Fungal Pathogens Isolated from Solanaceae plants can metabolize capsidiol or rishitin.

*Fusarium oxysporum* f. sp. *lycopersici* (*Fol*), a soilborne plant pathogen causes Fusarium wilt on tomato. *Fol* strain 9855-1 can metabolize capsidiol and showed tolerance to 100 μM capsidiol, while it cannot metabolize rishitin, which is produced by its host plant tomato.

*Colletotrichum coccodes* (*Cc*) is known to have a wide host range that causes anthracnose on tomato and onion, and black dot disease on potato. *Cc* strain PTK1 (isolated from potato) can metabolize both capsidiol and rishitin, and shows tolerance to 100 μM capsidiol. Metabolism of rishitin was not observed when treated with 100 μM but was induced when 500 μM were used. Thus, rishitin metabolism in *Cc* may be activated when *Cc* is exposed to high concentrations of rishitin.

*Stemphylium lycopersici* (*Sl*) has been isolated from a broad range of host plants, including tobacco and tomato. *Sl* strain KuNBY1 isolated from tobacco metabolizes both capsidiol and rishitin, and showed tolerance to 100 and 500 μM capsidiol and 100 μM rishitin.

*Fusarium coeruleum* (*Fc*) is the causal agent of potato dry rot. *Fc* strain K. Kita 37 can metabolize both capsidiol and rishitin, and showed tolerance to 500 μM capsidiol and 100 μM rishitin. Metabolism of rishitin was not observed when treated with 100 μM but induced when 500 μM were used. Similar to *Cc*, the metabolism of rishitin was induced when *Fc* was incubated in 500 μM rishitin.

*Alternaria solani* (*As*) is the causal pathogen of tomato and potato early blight. *As* strain KL1 isolated from potato can metabolize neither capsidiol nor rishitin, and is sensitive to both phytoalexins.

*Cercospora nicotianae* (*Cn*) is the pathogen causing tobacco frog-eye leaf spot. Although *Cn* strain CTC5 cannot metabolize capsidiol, it is tolerant to 100 μM capsidiol. *Cn* can partially metabolize rishitin.

*Gibellulopsis nigrescens* (*Gn*, former *Verticillium nigrescens*) is the causal agent of Verticillium wilt of potato. *Gn* strain kita44 can partially metabolize capsidiol, but didn't show tolerance to capsidiol.

*Ss*, *Sclerotinia sclerotiorum* (*Ss*) is a polyxenous pathogen causing white mold on a wide range of plant species. *Ss* isolate SU-1 (isolated from eggplant) can metabolize both capsidiol and rishitin, and showed tolerance to 100 μM capsidiol and 100 μM rishitin.

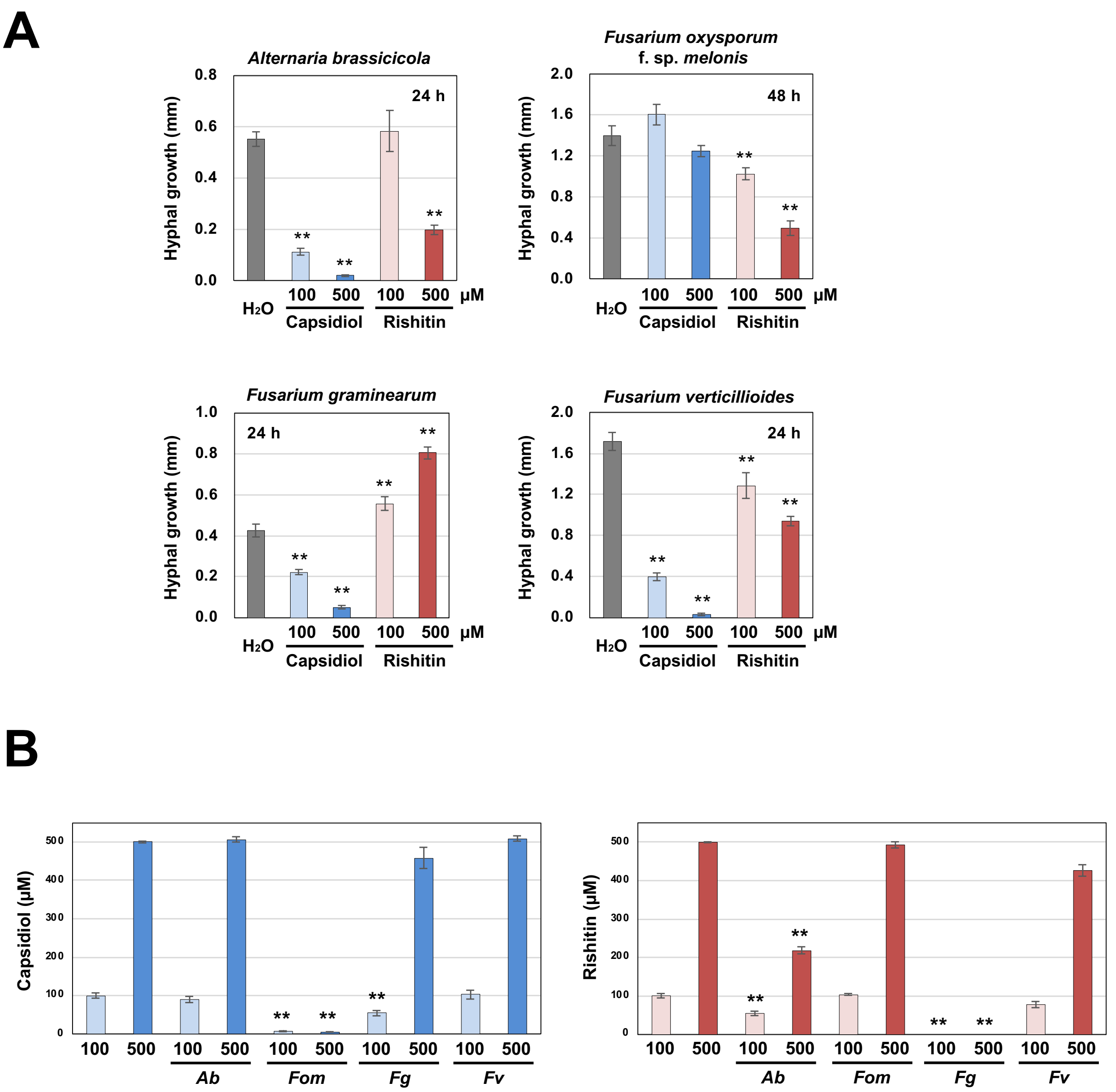


**Fig. S3.** Sensitivity and metabolic capacity of sesquiterpenoid phytoalexins in fungal pathogens. **(A)** Mycelial blocks (approx. 1 mm$^3$) of the indicated pathogen were incubated in 50 µl water, 100 µM or 500 µM capsidiol or rishitin. Outgrowth of hyphae from the mycelial block was measured after 24 h incubation (n = 6). **(B)** Residual capsidiol and rishitin was quantified by GC/MS after 48 h of incubation. Data marked with asterisks are significantly different from control as assessed by the two-tailed Student's *t*-test: $**P < 0.01$. *Ab*, *Alternaria brassicicola* (strain BA31 isolated from Broccoli); *Fom*, *Fusarium oxysporum* f. sp. *melonis* (strain Mel02010 isolated from melon); *Fg*, *F. graminearum* sensu stricto (strain 407011 isolated from wheat); Fv, *F. verticillioides* (strain Maize L-2 isolated from maize).

**Supplementary Note 2**

**Several Fungal Pathogens Isolated from Non-Solanaceae Plants Can also Metabolize Capsidiol or Rishitin.**

*Alternaria brassicicola* (*Ab*) is a necrotrophic pathogen that causes black spot disease, particularly on *Brassica* species. *Ab* strain BA31 (isolated from broccoli) can metabolize rishitin and showed tolerance to 100 µM rishitin.

*Fusarium oxysporum* f. sp. *melonis* (*Fom*) is pathogenic on melon, causing Fusarium wilt. *Fom* strain Mel02010 (isolated from melon, Namiki *et al.* 1994) can metabolize capsidiol and showed tolerance to 100 and 500 µM capsidiol.

*Fusarium graminearum* sensu stricto (*Fg*) is the causal agent of Fusarium head blight of cereals including barley and wheat. *Fg* strain 407011 (isolated from wheat, Suga *et al.* 2016) cannot metabolize capsidiol, and its growth was inhibited by capsidiol. Notably, in contrast, the growth of *Fg* is significantly enhanced in 100 and 500 µM rishitin and *Fg* strain 407011 can metabolize rishitin, indicating that *Fg* strain 407011 can metabolize and assimilate rishitin.

*F. verticillioides* (*Fv*) is a major fungal pathogen of cereals, such as wheat, sorghum and maize. Asymptomatic endophytic infection of this fungus in maize is also reported. *Fv* strain Maize L-2 (isolated from maize) cannot metabolize capsidiol and rishitin, and is sensitive to both phytoalexins.

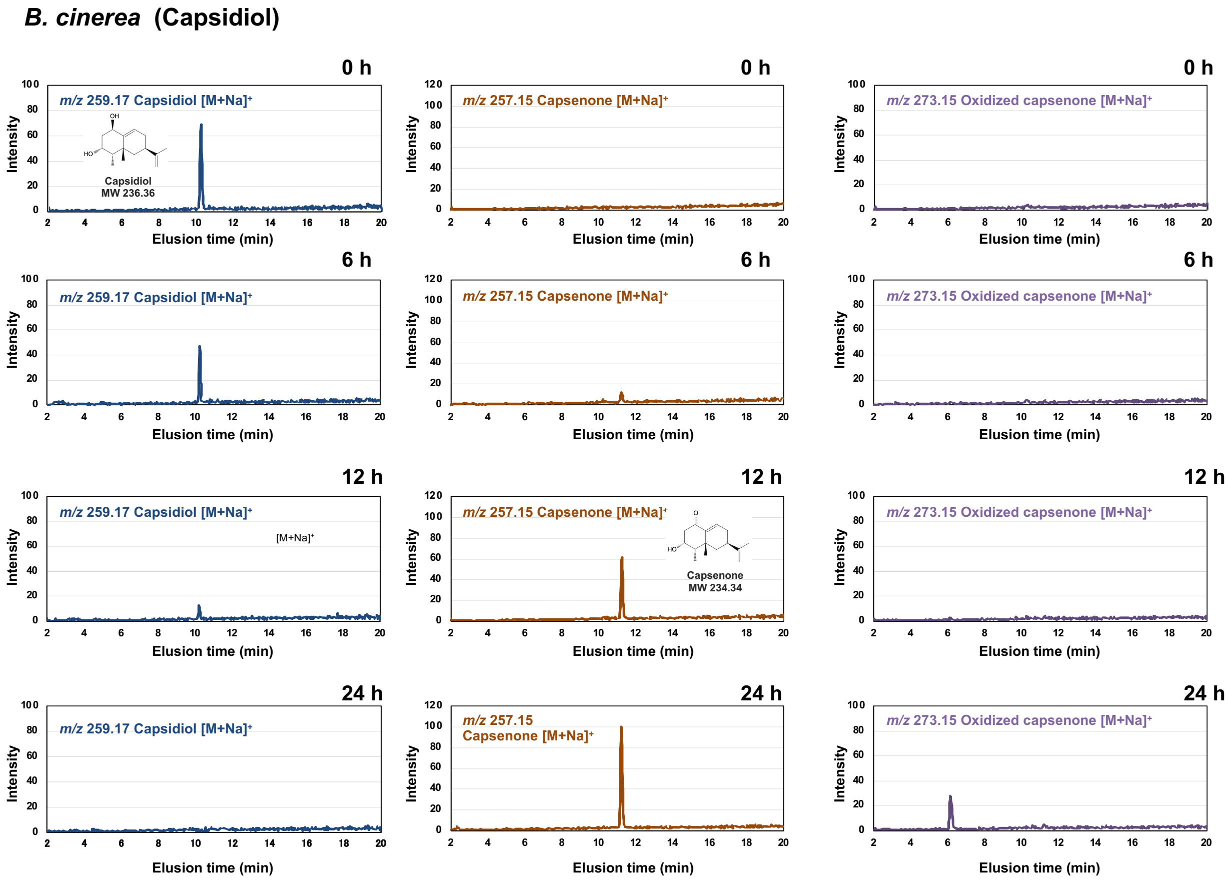


**Fig. S4.** Metabolism of capsidiol by *Botrytis cinerea*.

Mycelial blocks (approx. 1 $mm^3$) of *B. cinerea* were incubated in 50 µl of 100 µM capsidiol, and the residual capsidiol and its metabolites were detected by LC/MS.

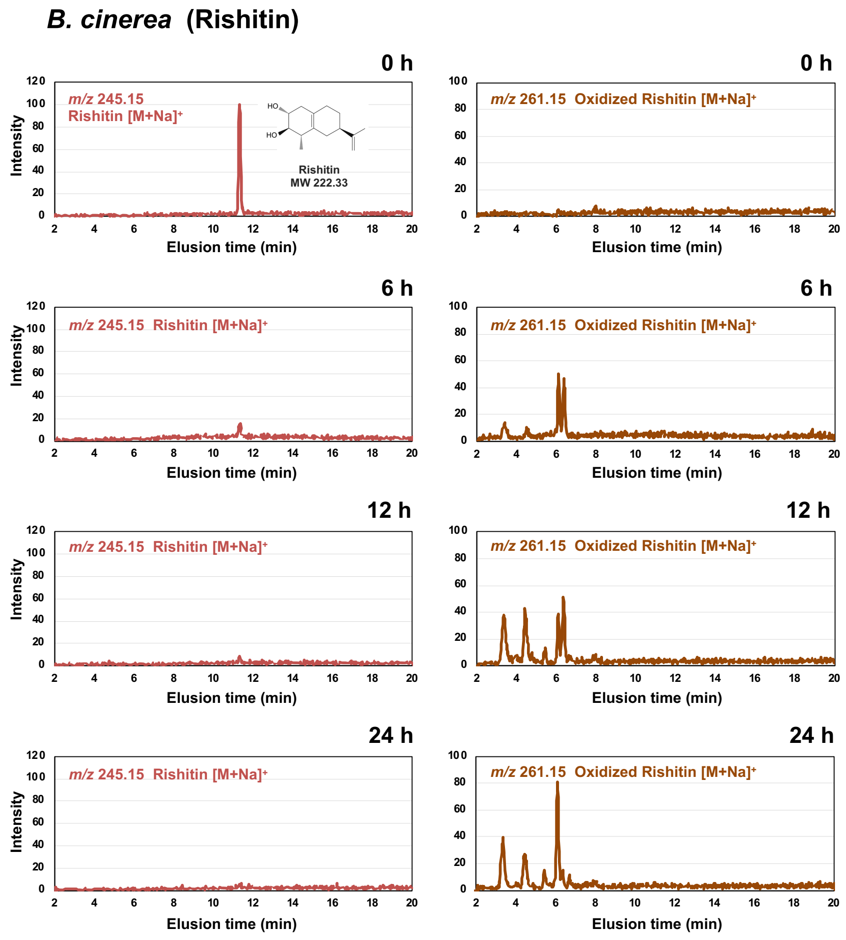


**Fig. S5.** Metabolism of rishitin by *Botrytis cinerea*.

Mycelial blocks (approx. 1 mm$^3$) of *B. cinerea* were incubated in 50 µl of 100 µM rishitin, and residual rishitin and its metabolites were detected by LC/MS.

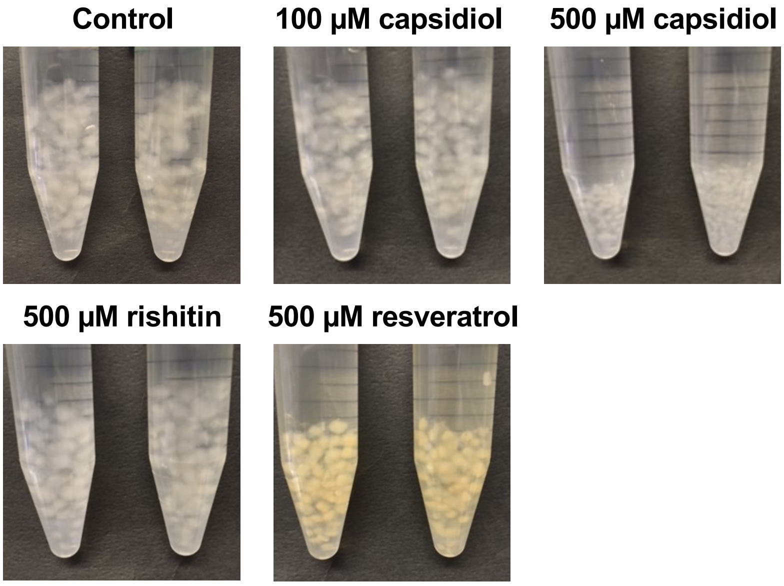


**Fig. S6.** Mycelial blocks of *B. cinerea* were incubated in CM medium or CM medium containing 100 µM or 500 µM capsidiol, 500 µM rishitin, or 500 µM resveratrol. Images were taken after 24 h of incubation.

**Table S1.** *Botrytis cinerea* genes significantly upregulated by treatment with capsidiol.

| Gene ID | FPKM value (Average) | | | | Log2 fold change (relative to control) | | | | | | Annotation | Motif |
|---|---|---|---|---|---|---|---|---|---|---|---|---|
| | Control | 100 µM Capsidiol | 500 µM Rishitin | 500 µM Resveratrol | 100 µM Capsidiol | P value | 500 µM Rishitin | P value | 500 µM Resveratrol | P value | | |
| Bcin08g00930.1 | 0.64 | 826.91 | 1.05 | 0.73 | 10.34 | 0.00 | 0.71 | 0.29 | 0.18 | 0.82 | Hypothetical protein | Short chain dehydrogenase; cl27753 |
| Bcin15g00050.1 | 1.12 | 398.16 | 0.87 | 0.89 | 8.47 | 0.00 | -0.37 | 0.13 | -0.33 | 0.39 | Bcayt1 | Transferase family; cl23789 |
| Bcin15g00040.1 | 0.47 | 147.83 | 0.31 | 0.38 | 8.30 | 0.00 | -0.61 | 0.44 | -0.31 | 0.63 | Hypothetical protein | Major Facilitator Superfamily; cd06174 |
| Bcin15g00030.1 | 0.12 | 25.49 | 0.28 | 0.08 | 7.72 | 0.01 | 1.22 | 0.27 | -0.58 | 0.72 | Hypothetical protein | Protein of unknown function (DUF3237); cl07905 |
| Bcin15g00020.1 | 0.13 | 17.32 | 0.35 | 0.70 | 7.04 | 0.00 | 1.42 | 0.42 | 2.42 | 0.01 | Hypothetical protein | Domain of unknown function (DUF1330); cl22966 |
| Bcin10g03040.1 | 2.64 | 225.97 | 3.19 | 2.76 | 6.42 | 0.00 | 0.27 | 0.51 | 0.06 | 0.92 | Hypothetical protein | Capsular polysaccharide synthesis protein; cl26275 |
| Bcin12g01750.1 | 1.48 | 73.95 | 2.42 | 1.06 | 5.64 | 0.00 | 0.71 | 0.18 | -0.48 | 0.53 | Hypothetical protein | Classical short-chain dehydrogenases/reductases (SDR); cd05233 |
| Bcin12g01130.1 | 3.43 | 76.66 | 2.80 | 3.16 | 4.48 | 0.00 | -0.29 | 0.41 | -0.12 | 0.78 | Hypothetical protein | Cytochrome P450; cl12078 |
| Bcin12g01740.1 | 0.59 | 11.39 | 0.59 | 0.69 | 4.27 | 0.00 | 0.00 | 1.00 | 0.23 | 0.88 | Hypothetical protein | ND |
| Bcin06g00510.1 | 0.59 | 8.89 | 0.74 | 2.27 | 3.91 | 0.04 | 0.32 | 0.29 | 1.94 | 0.19 | Bhp3 | Fungal hydrophobin; pfam06766 |
| Bcin01g01350.1 | 15.23 | 208.34 | 17.06 | 15.61 | 3.77 | 0.00 | 0.16 | 0.77 | 0.04 | 0.95 | Hypothetical protein | Short-chain dehydrogenases/reductases (SDR); cl25409 |
| Bcin13g05160.1 | 8.28 | 110.88 | 4.06 | 8.00 | 3.74 | 0.00 | -1.03 | 0.10 | -0.05 | 0.90 | Hypothetical protein | ND |
| Bcin14g02870.1 | 2.98 | 34.21 | 3.12 | 2.34 | 3.52 | 0.00 | 0.07 | 0.85 | -0.35 | 0.41 | Bcmfs1 | Fungal trichothecene efflux pump (TRI12); cl27908 |
| Bcin01g05890.1 | 13.67 | 155.81 | 51.04 | 16.72 | 3.51 | 0.00 | 1.90 | 0.00 | 0.29 | 0.57 | Bmr1 | Macrolide transporter ATP-binding /permease protein; cl28180 |
| Bcin02g07070.1 | 3.71 | 30.45 | 11.52 | 4.45 | 3.04 | 0.01 | 1.64 | 0.38 | 0.26 | 0.61 | Hypothetical protein | ND |
| Bcin01g03450.1 | 0.21 | 1.62 | 2.63 | 1.83 | 2.95 | 0.00 | 3.65 | 0.14 | 3.13 | 0.15 | Hypothetical protein | Domain of unknown function (DUF4185); cl16414 |
| Bcin13g05150.1 | 1013.70 | 7444.87 | 808.85 | 590.14 | 2.88 | 0.00 | -0.33 | 0.30 | -0.78 | 0.04 | Hypothetical protein | Basic leucine zipper (bZIP), DNA-binding and dimerization domain; cd12193 |
| Bcin12g06760.1 | 20.11 | 134.82 | 3.86 | 8.10 | 2.75 | 0.04 | -2.38 | 0.11 | -1.31 | 0.23 | Hypothetical protein | ND |
| Bcin04g05650.1 | 3.77 | 21.76 | 17.45 | 5.47 | 2.53 | 0.02 | 2.21 | 0.32 | 0.54 | 0.12 | Hypothetical protein | Bicupin, oxalate decarboxylase family; TIGR03404 |
| Bcin08g02330.1 | 4.13 | 21.83 | 5.38 | 4.27 | 2.40 | 0.00 | 0.38 | 0.31 | 0.05 | 0.92 | Bcria1 | Elongation factor Tu GTP binding domain; cl27769 |
| Bcin10g01350.1 | 1.56 | 8.11 | 1.44 | 1.07 | 2.38 | 0.00 | -0.12 | 0.81 | -0.54 | 0.41 | Hypothetical protein | Short-chain dehydrogenases/reductases (SDR); cl25409 |
| Bcin16g00810.1 | 0.98 | 4.29 | 2.59 | 0.92 | 2.13 | 0.02 | 1.40 | 0.39 | -0.09 | 0.94 | Hypothetical protein | ND |
| Bcin16g01490.1 | 0.42 | 1.83 | 23.66 | 3.09 | 2.12 | 0.02 | 5.81 | 0.03 | 2.87 | 0.38 | Hypothetical protein | Cytochrome P450; cl12078 |
| Bcin15g00060.1 | 1.02 | 4.43 | 1.80 | 1.38 | 2.11 | 0.01 | 0.81 | 0.00 | 0.43 | 0.19 | Hypothetical protein | Glutathione S-transferase; cl25459 |
| Bcin10g05150.1 | 61.52 | 260.33 | 90.00 | 85.38 | 2.08 | 0.00 | 0.55 | 0.02 | 0.47 | 0.07 | Bctif6 | Eukaryotic translation initiation factor 6-like protein; PTZ00136 |

ND, not detected.

**Table S2.** *Botrytis cinerea* genes significantly upregulated by treatment with rishitin

| Gene ID | FPKM value (Average) | | | | Log2 fold change (relative to control) | | | | | | Annotation | Motif |
|---|---|---|---|---|---|---|---|---|---|---|---|---|
| | Control | 100 µM Capsidiol | 500 µM Rishitin | 500 µM Resveratrol | 100 µM Capsidiol | P value | 500 µM Rishitin | P value | 500 µM Resveratrol | P value | | |
| Bcin07g05430.1 | 0.21 | 0.04 | 24.89 | 2.48 | -2.32 | 0.41 | 6.86 | 0.01 | 3.54 | 0.02 | Hypothetical protein | Cytochrome P450; cl12078 |
| Bcin13g00710.1 | 0.80 | 0.51 | 69.45 | 61.88 | -0.64 | 0.10 | 6.44 | 0.01 | 6.28 | 0.03 | BcatrB | Macrolide transporter ATP-binding /permease protein; cl28180 |
| Bcin08g04910.1 | 0.50 | 0.18 | 29.52 | 1.50 | -1.50 | 0.05 | 5.89 | 0.02 | 1.58 | 0.00 | Hypothetical protein | Phenylcoumaran benzylic ether reductase like, atypical SDRs; cd05259 |
| Bcin16g01490.1 | 0.42 | 1.83 | 23.66 | 3.09 | 2.12 | 0.02 | 5.81 | 0.03 | 2.87 | 0.38 | Hypothetical protein | Cytochrome P450; cl12078 |
| Bcin06g00650.1 | 0.09 | 0.11 | 3.65 | 3.52 | 0.26 | 0.84 | 5.28 | 0.00 | 5.22 | 0.01 | Hypothetical protein | Cytochrome P450; cl12078 |
| Bcin07g02220.1 | 1.07 | 2.90 | 17.52 | 1.35 | 1.43 | 0.02 | 4.03 | 0.01 | 0.34 | 0.53 | Bmr3 | Macrolide transporter ATP-binding /permease protein; cl28180 |
| Bcin03g04480.1 | 2.45 | 2.77 | 32.63 | 7.67 | 0.17 | 0.67 | 3.73 | 0.02 | 1.65 | 0.04 | Hypothetical protein | Classical short-chain dehydrogenases/reductases; cd05233 |
| Bcin08g04920.1 | 0.62 | 0.59 | 6.84 | 1.10 | -0.08 | 0.47 | 3.45 | 0.03 | 0.82 | 0.10 | Hypothetical protein | GAL4-like Zn2Cys6 binuclear cluster DNA-binding domain; cd00067 |
| Bcin04g03050.1 | 0.55 | 0.71 | 5.74 | 1.26 | 0.37 | 0.61 | 3.39 | 0.00 | 1.20 | 0.11 | Bcgpr5 | Seven-transmembrane G protein-coupled receptor superfamily; cl28897 |
| Bcin13g02720.1 | 11.26 | 26.69 | 105.31 | 14.45 | 1.25 | 0.01 | 3.23 | 0.00 | 0.36 | 0.30 | BcatrD | Pleiotropic Drug Resistance (PDR) Family protein; TIGR00956 |
| Bcin06g07120.1 | 5.21 | 5.26 | 43.77 | 8.22 | 0.02 | 0.94 | 3.07 | 0.01 | 0.66 | 0.32 | Hypothetical protein | Hypothetical protein; Provisional; cl27550 |
| Bcin14g01070.1 | 48.96 | 53.91 | 386.41 | 123.65 | 0.14 | 0.51 | 2.98 | 0.01 | 1.34 | 0.00 | Hypothetical protein | Medium chain reductase/dehydrogenase/zinc-dependent alcohol dehydrogenase-like family; cl16912 |
| Bcin05g05130.1 | 1.49 | 1.90 | 11.27 | 1.39 | 0.35 | 0.28 | 2.92 | 0.02 | -0.10 | 0.72 | Hypothetical protein | ND |
| Bcin05g05120.1 | 1.25 | 1.82 | 9.36 | 1.65 | 0.54 | 0.16 | 2.90 | 0.01 | 0.39 | 0.39 | Hypothetical protein | Short chain dehydrogenase; cl27753 |
| Bcin16g02850.1 | 2.59 | 3.08 | 18.78 | 6.32 | 0.25 | 0.16 | 2.86 | 0.00 | 1.29 | 0.01 | Hypothetical protein | FAD/FMN-containing dehydrogenase; COG0277 |
| Bcin03g07770.1 | 4.39 | 9.82 | 28.87 | 106.82 | 1.16 | 0.03 | 2.72 | 0.04 | 4.60 | 0.00 | Bcalo4 | NAD(P)H-nitrite reductase, large subunit; cl26176 |
| Bcin07g04570.1 | 8.17 | 7.47 | 44.67 | 11.76 | -0.13 | 0.45 | 2.45 | 0.01 | 0.53 | 0.14 | Hypothetical protein | UDP-glucoronosyl and UDP-glucosyl transferase; cl26154 |
| Bcin08g04720.1 | 0.89 | 2.16 | 4.33 | 1.72 | 1.27 | 0.08 | 2.27 | 0.02 | 0.94 | 0.20 | Hypothetical protein | ND |
| Bcin15g01970.1 | 2.25 | 4.02 | 10.77 | 0.48 | 0.84 | 0.17 | 2.26 | 0.00 | -2.23 | 0.04 | Hypothetical protein | ND |
| Bcin01g00080.1 | 2.72 | 3.52 | 12.75 | 5.38 | 0.37 | 0.21 | 2.23 | 0.00 | 0.98 | 0.04 | Bcboa8 | 2-polyprenyl-6-methoxyphenol hydroxylase and related FAD-dependent oxidoreductases; COG0654 |
| Bcin11g01310.1 | 2.34 | 1.12 | 10.85 | 3.17 | -1.06 | 0.01 | 2.21 | 0.00 | 0.44 | 0.26 | Hypothetical protein | Alpha/beta hydrolases; cl21494 |
| Bcin09g00450.1 | 1.75 | 3.54 | 7.65 | 3.66 | 1.02 | 0.00 | 2.13 | 0.02 | 1.07 | 0.00 | Hypothetical protein | Alpha/beta hydrolases; cl21494 |
| Bcin08g04610.1 | 18.99 | 24.39 | 81.98 | 21.31 | 0.36 | 0.26 | 2.11 | 0.00 | 0.17 | 0.70 | Hypothetical protein | Short chain dehydrogenase; cl27753 |
| Bcin15g04480.1 | 9.56 | 17.27 | 40.58 | 16.64 | 0.85 | 0.01 | 2.08 | 0.00 | 0.80 | 0.16 | Hypothetical protein | Short chain dehydrogenase; cl27753 |
| Bcin11g02050.1 | 5.51 | 8.15 | 23.26 | 5.98 | 0.56 | 0.04 | 2.08 | 0.01 | 0.12 | 0.74 | Hypothetical protein | 17-beta-hydroxysteroid dehydrogenase XI-like, short-chain dehydrogenases/reductases; cd05339 |
| Bcin06g01060.1 | 1.23 | 1.20 | 5.12 | 3.73 | -0.04 | 0.95 | 2.06 | 0.00 | 1.60 | 0.10 | Hypothetical protein | ND |
| Bcin04g06990.1 | 22.62 | 26.67 | 93.16 | 7.50 | 0.24 | 0.24 | 2.04 | 0.00 | -1.59 | 0.00 | Hypothetical protein | Glycosyl hydrolase family 43; cd08998 |

ND, not detected.

**Table S3.** *Botrytis cinerea* genes significantly upregulated by treatment with resveratrol

| Gene ID | FPKM value (Average) | | | | Log2 fold change (relative to control) | | | | | | Annotation | Motif |
|---|---|---|---|---|---|---|---|---|---|---|---|---|
| | Control | 100 µM Capsidiol | 500 µM Rishitin | 500 µM Resveratrol | 100 µM Capsidiol | P value | 500 µM Rishitin | P value | 500 µM Resveratrol | P value | | |
| Bcin14g05330.1 | 0.60 | 0.56 | 1.37 | 102.69 | -0.12 | 0.89 | 1.18 | 0.53 | 7.41 | 0.00 | Hypothetical protein | Intradiol_dioxygenase_like; cd03457 |
| Bcin13g00710.1 | 0.80 | 0.51 | 69.45 | 61.88 | -0.64 | 0.10 | 6.44 | 0.01 | 6.28 | 0.03 | BcatrB | Macrolide transporter ATP-binding /permease protein; cl28180 |
| Bcin06g00650.1 | 0.09 | 0.11 | 3.65 | 3.52 | 0.26 | 0.84 | 5.28 | 0.00 | 5.22 | 0.01 | Hypothetical protein | Cytochrome P450; cl12078 |
| Bcin03g07770.1 | 4.39 | 9.82 | 28.87 | 106.82 | 1.16 | 0.03 | 2.72 | 0.04 | 4.60 | 0.00 | Bcalo4 | NAD(P)H-nitrite reductase, large subunit; cl26176 |
| Bcin16g04210.1 | 26.92 | 16.27 | 18.72 | 556.50 | -0.73 | 0.04 | -0.52 | 0.33 | 4.37 | 0.00 | Bcidl1 | Intradiol_dioxygenase_like; cd03457 |
| Bcin14g02510.1 | 44.07 | 12.30 | 211.14 | 816.50 | -1.84 | 0.00 | 2.26 | 0.09 | 4.21 | 0.00 | Bclcc2 | Multicopper oxidase with three cupredoxin domains; cl28276 |
| Bcin07g04100.1 | 2.47 | 3.02 | 2.12 | 45.11 | 0.29 | 0.56 | -0.22 | 0.78 | 4.19 | 0.01 | Hypothetical protein | S-adenosylmethionine-dependent methyltransferases class I; cl17173 |
| Bcin04g02650.1 | 2.54 | 4.34 | 8.37 | 31.25 | 0.77 | 0.04 | 1.72 | 0.04 | 3.62 | 0.00 | Hypothetical protein | FAD binding domain; cl27552 |
| Bcin07g05430.1 | 0.21 | 0.04 | 24.89 | 2.48 | -2.32 | 0.41 | 6.86 | 0.01 | 3.54 | 0.02 | Hypothetical protein | Cytochrome P450; cl12078 |
| Bcin06g06350.1 | 38.12 | 41.34 | 51.62 | 413.20 | 0.12 | 0.69 | 0.44 | 0.49 | 3.44 | 0.00 | Bcnqo1 | NADPH-dependent FMN reductase; cl00438 |
| Bcin05g07100.1 | 67.10 | 93.43 | 151.28 | 711.57 | 0.48 | 0.39 | 1.17 | 0.26 | 3.41 | 0.04 | Hypothetical protein | Redox-sensitive bicupin YhaK, pirin superfamily; COG1741 |
| Bcin15g05080.1 | 4.05 | 1.47 | 2.17 | 27.75 | -1.46 | 0.15 | -0.90 | 0.25 | 2.78 | 0.02 | Bcgas5 | Glycosyl hydrolase family 1; cl23725 |
| Bcin06g05110.1 | 22.06 | 33.15 | 63.56 | 148.92 | 0.59 | 0.00 | 1.53 | 0.05 | 2.76 | 0.02 | Hypothetical protein | Redox-sensitive bicupin YhaK, pirin superfamily; COG1741 |
| Bcin04g06040.1 | 4.29 | 1.80 | 8.56 | 28.76 | -1.25 | 0.25 | 1.00 | 0.14 | 2.75 | 0.01 | Hypothetical protein | CESA_CelA_like; cd06421 |
| Bcin07g03750.1 | 14.49 | 15.26 | 16.26 | 89.34 | 0.07 | 0.79 | 0.17 | 0.38 | 2.62 | 0.00 | Hypothetical protein | ND |
| Bcin15g00020.1 | 0.13 | 17.32 | 0.35 | 0.70 | 7.04 | 0.00 | 1.42 | 0.42 | 2.42 | 0.01 | Hypothetical protein | Domain of unknown function (DUF1330); cl22966 |
| Bcin02g04490.1 | 41.96 | 44.10 | 101.46 | 221.26 | 0.07 | 0.40 | 1.27 | 0.00 | 2.40 | 0.00 | Hypothetical protein | The Class III extradiol dioxygenase, 4,5-DOPA dioxygenase, cd07363 |
| Bcin03g05130.1 | 1.06 | 0.87 | 0.95 | 5.44 | -0.27 | 0.43 | -0.15 | 0.67 | 2.37 | 0.00 | Bcalo3 | NAD(P)H-nitrite reductase, large subunit; cl26176 |
| Bcin03g06320.1 | 96.23 | 81.03 | 77.15 | 433.90 | -0.25 | 0.41 | -0.32 | 0.19 | 2.17 | 0.00 | Hypothetical protein | Methyltransferase domain; pfam13649 |
| Bcin04g06920.1 | 3.39 | 2.82 | 4.41 | 15.01 | -0.26 | 0.57 | 0.38 | 0.47 | 2.15 | 0.01 | Hypothetical protein | alpha/beta hydrolases; cl21494 |
| Bcin11g06310.2 | 0.22 | 0.34 | 2.74 | 0.98 | 0.58 | 0.12 | 3.61 | 0.13 | 2.13 | 0.02 | Hypothetical protein | Phenylcoumaran benzylic ether reductase like, atypical SDRs; cd05259 |
| Bcin11g02620.1 | 4.91 | 6.13 | 12.84 | 20.03 | 0.32 | 0.37 | 1.39 | 0.10 | 2.03 | 0.00 | Bchol1 | Major Facilitator Superfamily; cd06174 |
| Bcin02g03510.1 | 30.05 | 22.31 | 44.17 | 120.32 | -0.43 | 0.22 | 0.56 | 0.06 | 2.00 | 0.00 | Hypothetical protein | ND |

ND, not detected.

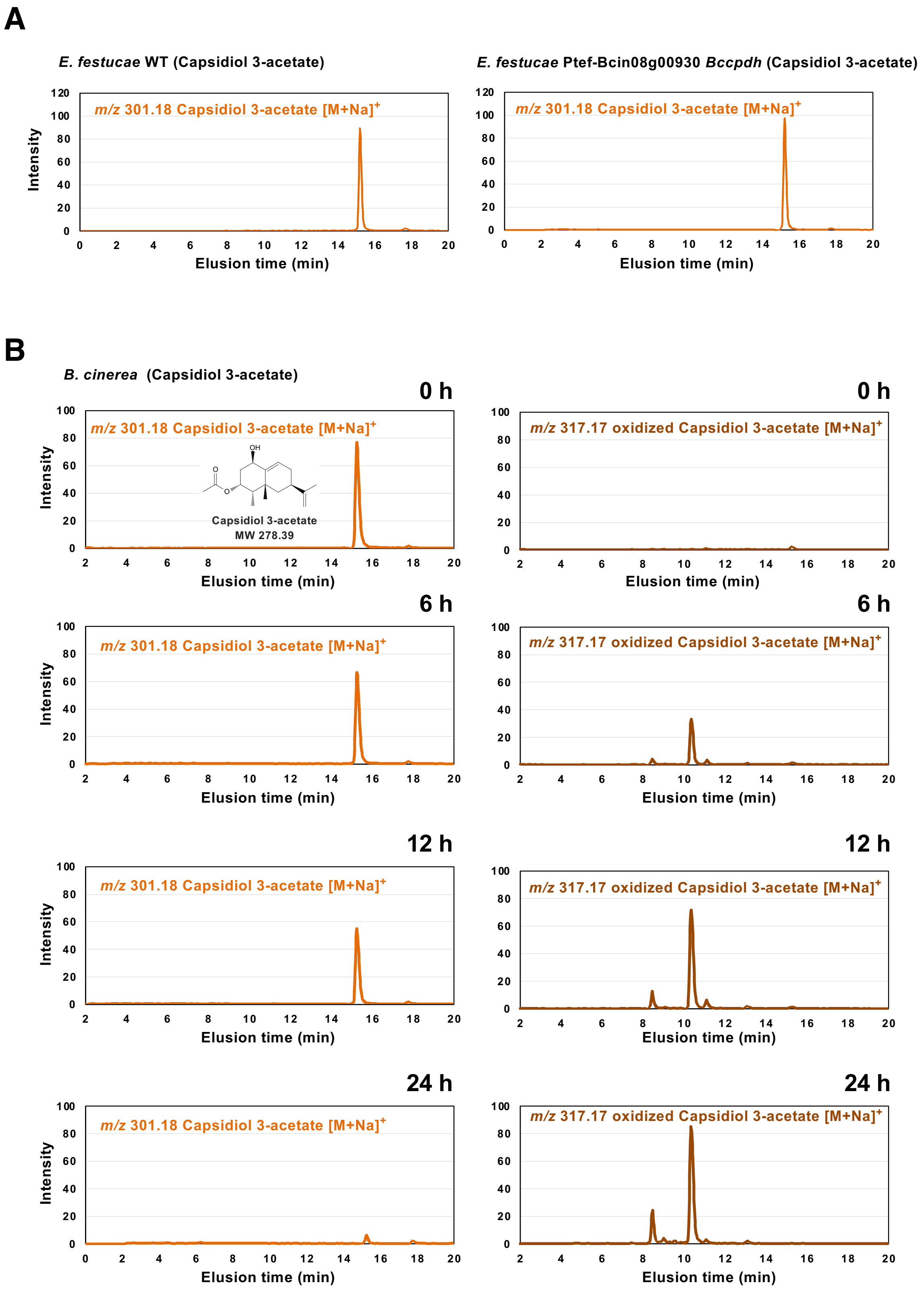


**Fig. S7. (A)** Mycelia of *E. festucae* wild type (WT) or transformants expressing Bcin08g00930 (*BcCPDH*) were incubated in 100 µM capsidiol acetate for 48 h. Capsidiol acetate was detected by LC/MS. **(B)** Mycelia of *B. cinerea* was incubated in 100 µM capsidiol acetate for indicated time and capsidiol acetate and their oxidized metabolites were detected by LC/MS.

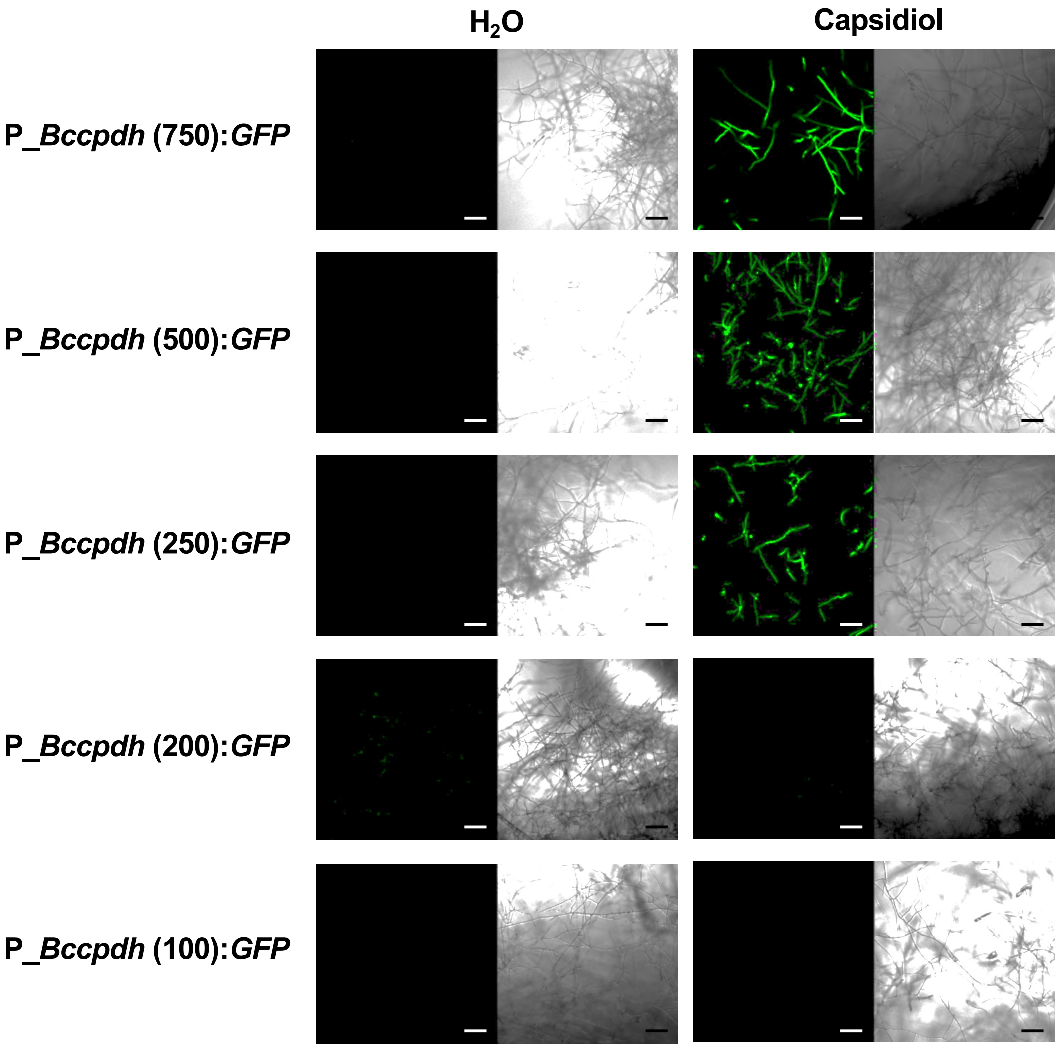


**Fig. S8.** *B. cinerea* transformants expressing GFP under the control of different lengths (upstream from the start codon of the gene) of *Bccpdh* promoter were incubated in water or 500 μM capsidiol. Expression of GFP was monitored by confocal laser microscopy 1 day after the treatment. Bars = 150 μm.

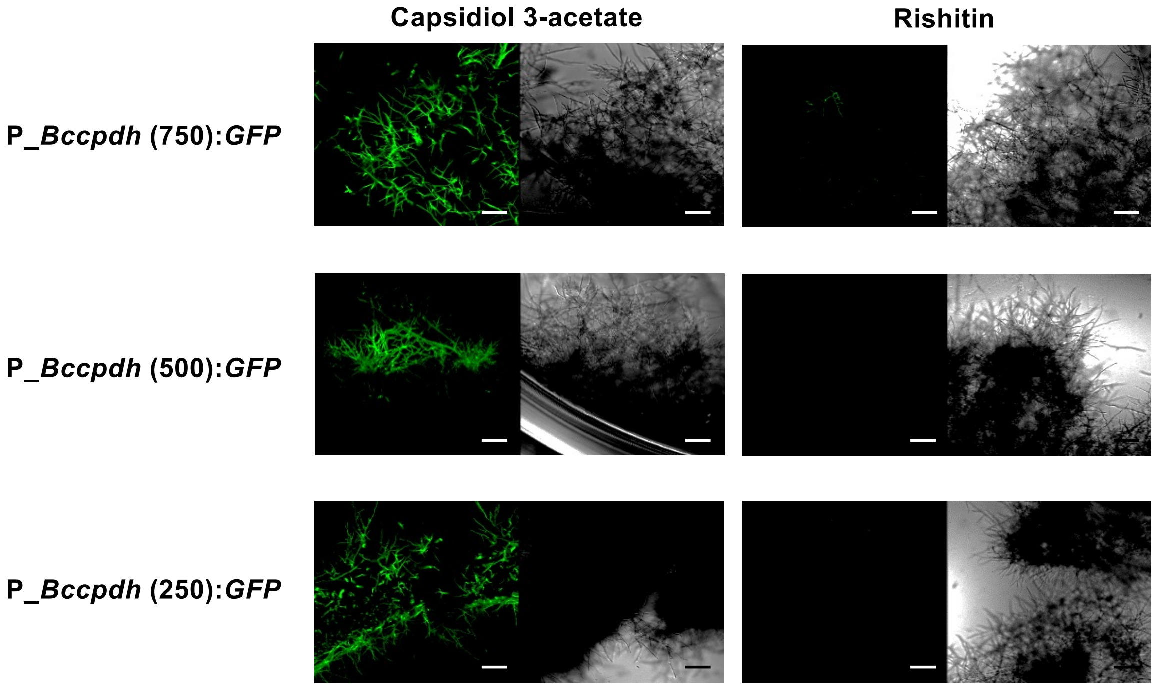


**Fig. S9.** *B. cinerea* transformants expressing GFP under the control of different lengths (upstream from the start codon of the gene) of *Bccpdh* promoter were incubated in 500 µM capsidiol 3-acetate or rishitin. Expression of GFP was monitored by confocal laser microscopy 1 day after the treatment. Bars = 150 µm.

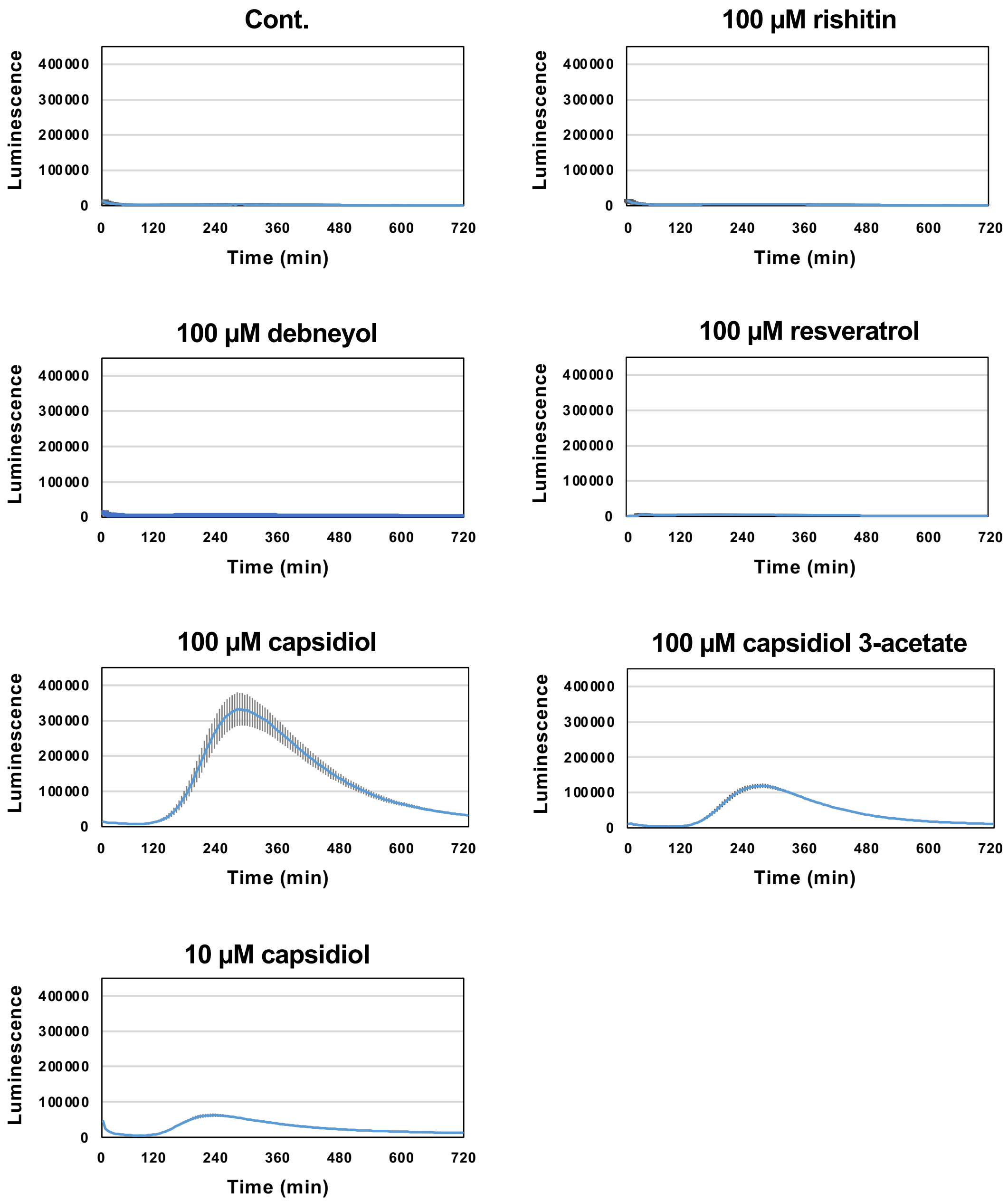


**Fig. S10.** Luminescence intensity of *B. cinerea* transformant P_*Bccpdh:Luc* containing the *Luciferase* gene under the control of 250 bp *Bccpdh* promoter. The transformant was incubated in water, 100 µM capsidiol, capsidiol 3-acetate, rishitin, resveratrol or debneyol or 10 µM capsidiol. 50 µM D-luciferin was used as the substrate of luciferase. Data are means ±SE (n = 4).

**Supplementary Note 3**

***Bccpdh* promoter is activated by capsidiol in a concentration-dependent manner.**

*B. cinerea* transformant P_*Bccpdh:Luc* was produced for the expression of *Luciferase* (*Luc*) under the control of 250 bp *Bccpdh* promoter (250 bp upstream from the start codon of the gene). The *Bccpdh* promoter was activated within the first 2 h after incubation with either 10 or 100 μM capsidiol. The peak of promoter activation was approx. at 5 h for 100 μM capsidiol and within 4 h for 10 μM capsidiol, and the degree and duration of promoter activation was concentration dependent. This result indicates that the activity of the *Bccpdh* promoter immediately decreases once capsidiol is metabolized.

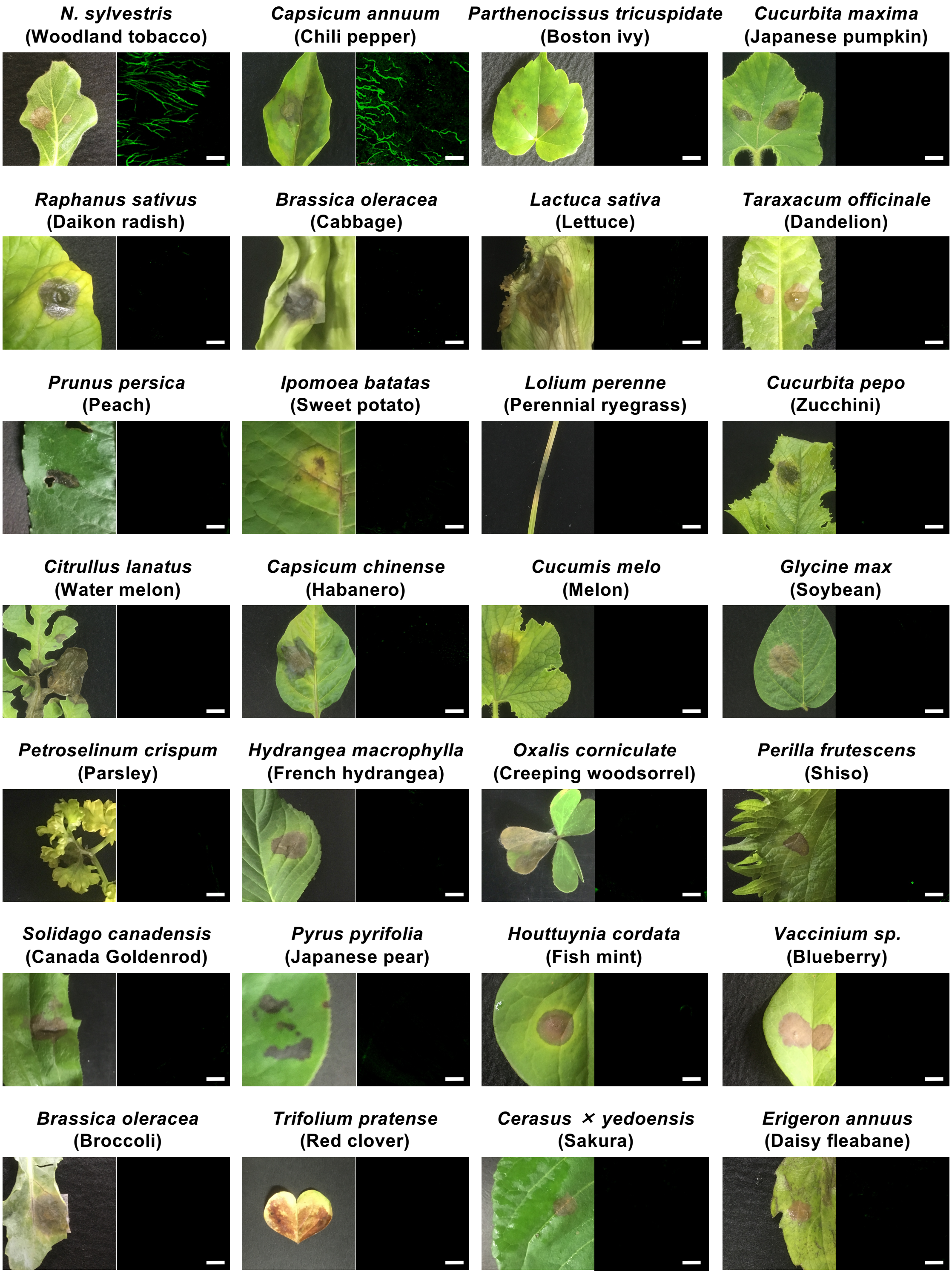


**Fig. S11.** *B. cinerea Bccpdh* promoter is activated during the infection in plants producing capsidiol. Leaves of indicated plants were inoculated with the mycelia of *B. cinerea* P_*Bccpdh*:*GFP* transformant and hyphae at the edge of the lesion was observed by confocal laser microscopy 2 or 3 d after the inoculation. Bars = 100 µm.

**Table S4.** Expression of GFP in *Botrytis cinerea* P_*Bccpdh* :*GFP* transformant on different plant species.

| Family | Host plant | Common name | Disease symptom | Expression of GFP in *B. cinerea* P_*Bccpdh* :*GFP* | Production of capsidiol (Reference) |
|---|---|---|---|---|---|
| Solanaceae | *Nicotiana benthamiana* | Benth | + | + | Matsukawa *et al.* (2013) |
| Solanaceae | *Nicotiana tabacum* | Tobacco | + | + | Bailey *et al.* (1975) |
| Solanaceae | *Nicotiana sylvestris* | Woodland tobacco | + | + | Bohlmann *et al.* (2002) |
| Solanaceae | *Capsicum annuum* | Bell pepper | + | + | Molot *et al.* (1981) |
| Solanaceae | *Capsicum annuum* | Chilli pepper | + | + | Molot *et al.* (1981) |
| Solanaceae | *Solunum lycopercicum* | Tomato | + | - | n.r |
| Solanaceae | *Solunum tuberosum* | Potato | + | - | n.r |
| Solanaceae | *Solunum melongena* | Eggplant | + | - | n.r |
| Rosaceae | *Fragaria* × *ananassa* | Strawberry | + | - | n.r |
| Rosaceae | *Pyrus pyrifolia* | Japanese pear | + | - | n.r |
| Rosaceae | *Prunus persica* | Peach | + | - | n.r |
| Rosaceae | *Cerasus* × *yedoensis* | Sakura | + | - | n.r |
| Rosaceae | *Malus domestica* | Apple | + | - | n.r |
| Rosaceae | *Rosa hybrida* | Miniature rose | + | - | n.r |
| Brassicaceae | *Brassica oleracea* var. *capitata* | Cabbage | + | - | n.r |
| Brassicaceae | *Arabidopsis thaliana* | Thale cress | + | - | n.r |
| Brassicaceae | *Brassica oleracea* var. *italica* | Broccoli | + | - | n.r |
| Brassicaceae | *Raphanus sativus* var. *longipinnatus* | Daikon radish | + | - | n.r |
| Brassicaceae | *Brassica rapa* var. *pekinensis* | Chinese cabbage | + | - | n.r |
| Brassicaceae | *Brassica rapa* var. *rapa* | Turnip | + | - | n.r |
| Fabaceae | *Phaseolus vulgaris* | Common bean | + | - | n.r |
| Fabaceae | *Pisum sativum* | Pea | + | - | n.r |
| Fabaceae | Glycine max | Soybean | + | - | n.r |
| Fabaceae | *Trifolium repens* | White clover | + | - | n.r |
| Fabaceae | *Trifolium pratense* | Red clover | + | - | n.r |
| Asteraceae | *Taraxacum officinale* | Dandelion | + | - | n.r |
| Asteraceae | *Erigeron annuus* | Annual fleabane | + | - | n.r |
| Asteraceae | *Lactuca sativa* | Lettuce | + | - | n.r |
| Asteraceae | *Solidago altissima* | Tall goldenrod | + | - | n.r |
| Asteraceae | *Glebionis coronaria* | Crown daisy | + | - | n.r |
| Asteraceae | *Chrysanthemum* × *morifolium* | Florist's daisy | + | - | n.r |
| Cucurbitaceae | *Cucurbita pepo* | Zucchini | + | - | n.r |
| Cucurbitaceae | *Cucurbita maxima* | Winter squash | + | - | n.r |
| Cucurbitaceae | *Cucumis sativus L.* | Cucumber | + | - | n.r |
| Cucurbitaceae | *Citrullus lanatus* | Watermelon | + | - | n.r |
| Amaryllidaceae | *Allium fistulosum* | Welsh onion | + | - | n.r |
| Amaryllidaceae | *Allium cepa* | Onion | + | - | n.r |
| Amaryllidaceae | *Allium tuberosum* | Chinese chives | + | - | n.r |
| Vitaceae | *Vitis* × *labruscana* | Grape | + | - | n.r |
| Vitaceae | *Parthenocissus tricuspidata* | Boston Ivy | + | - | n.r |
| Lamiaceae | *Perilla frutescens* var. *crispa* | Shiso | + | - | n.r |
| Lamiaceae | *Ocimum basilicum* | Basil | + | - | n.r |
| Poaceae | *Lolium perenne* | Perennial ryegrass | + | - | n.r |
| Sauruaceae | *Houttuynia cordata* | Fish mint | + | - | n.r |
| Moraceae | *Morus australis* | Mulberry | + | - | n.r |
| Malvaceae | Abelmoschus esculentus | Okra | + | - | n.r |
| Ebenaceae | *Diospyros kaki* | Persimmon | + | - | n.r |
| Asparagaceae | *Asparagus officinalis* | Asparagus | + | - | n.r |
| Ericaceae | *Vaccinium* ssp. | Blueberry | + | - | n.r |
| Hydrangeaceae | Hydrangea macrophylla | Hydrangea | + | - | n.r |
| Convolvulaceae | *Ipomoea batatas* | Sweet potato | + | - | n.r |
| Caryophyllaceae | *Dianthus caryophyllus* | Carnation | + | - | n.r |
| Oxalidaceae | *Oxalis corniculata* | Creeping woodsorrel | + | - | n.r |

n.r. not reported.

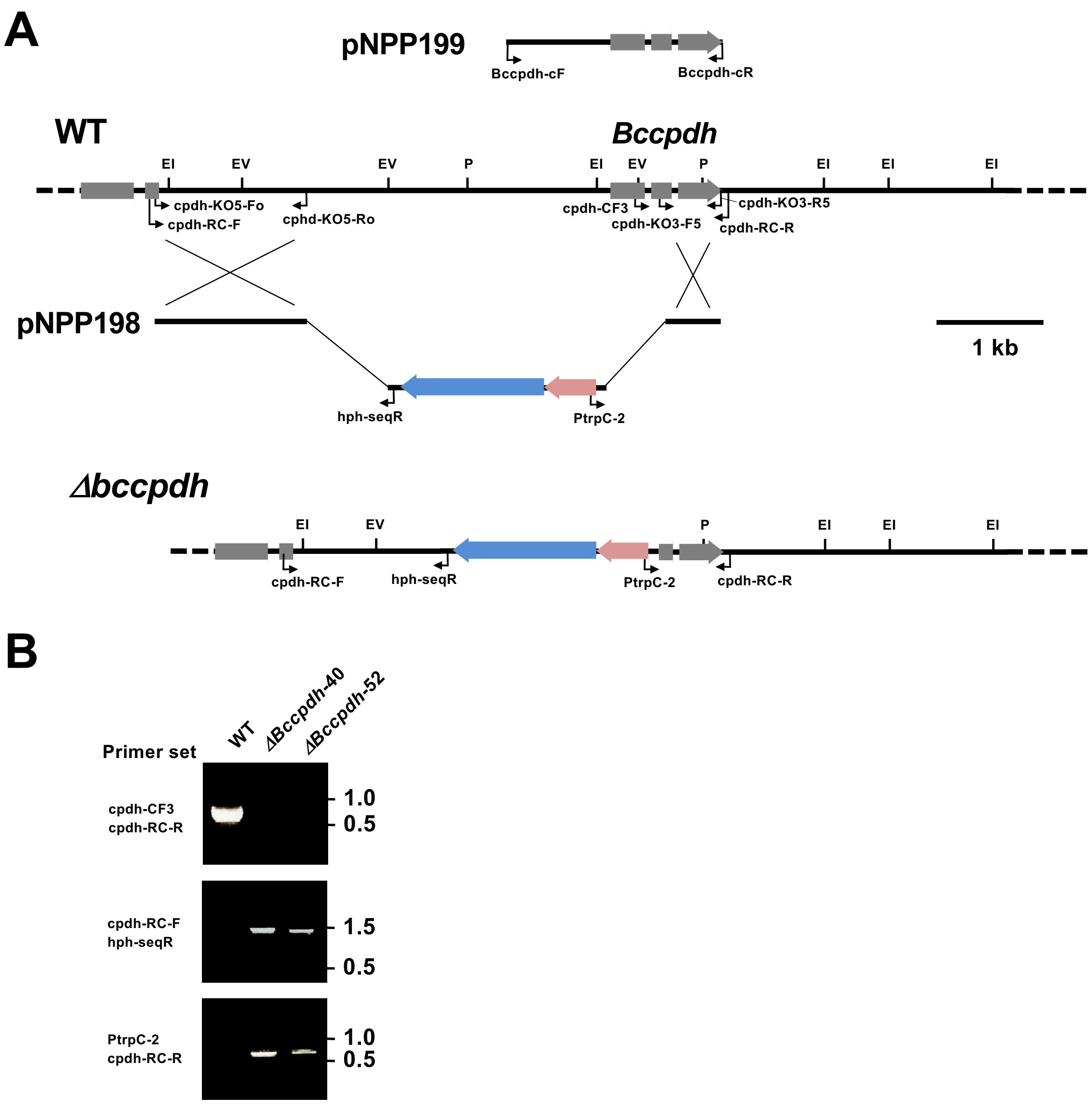


**Fig. S12.** Targeted gene replacement of the *B. cinerea Bccpdh* locus.

**(A)** Physical map of the *Bccpdh* wild-type (WT) genomic region, linear insert of *Bccpdh* replacement construct pNPP198 and complementation construct pNPP199, showing restriction enzyme sites for *Eco*RV (EV), *Eco*RI (EI) and *Pst*I (P). The mutated genomic locus of *Bccpdh* deletion mutant (*Δbccpdh*) is depicted to show homologous recombination of the *hph* cassette. Primers used for the construction of deletion vector and screening for the replacement event are indicated by arrowheads. **(B)** Confirmation of gene disruption in isolated *Δbccpdh* strains by PCR. Genomic DNA from *B. cinerea* wild type and *Δbccpdh* strains were used for PCR with indicated primers.

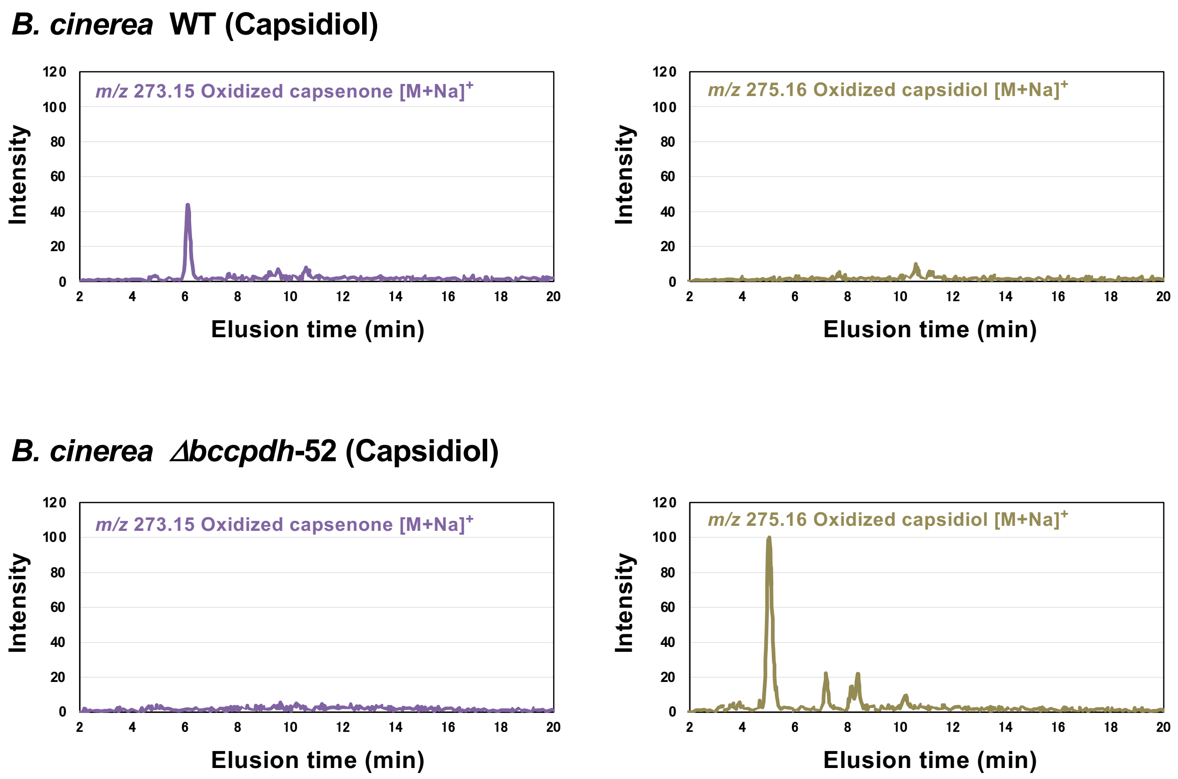


**Fig. S13.** Mycelial blocks (approx. 1 $mm^3$) of *B. cinerea* wild type (WT) or *Bccpdh* KO mutant strain (*Δbccpdh*-52) were incubated in 50 µl of 100 µM capsidiol for 4 days. Oxidized capsenone and oxidized capsidiol were detected by LC/MS.

### Supplementary Note 4

#### Capsidiol is oxidized in *Δbccpdh* by a cytochrome P450 encoded by Bcin16g01490.

After the incubation of capsidiol with *B. cinerea Δbccpdh*, oxidized capsidiol (one major and at least two minor peaks) were detected, while oxidized capsidiol was not detected in the metabolites after the incubation with wild type *B. cinerea* (Fig. S13), probably because capsidiol is quickly metabolized to capsenone (Fig. S4). The major oxidized capsidiol was detected after the incubation of capsidiol with *E. festucae* expressing Bcin16g01490 encoding a cytochrome P450 (Fig. S14B), indicating that the lack of BcCPDH alters the pathway in *Δbccpdh* towards a direct oxidation of capsidiol by Bcin16g01490 (Fig. S14C). It should be noted, however, that a substantial amount of capsidiol is remaining after the incubation with *B. cinerea Δbccpdh* or *E. festucae* expressing Bcin16g01490, indicating that this cytochrome P450 does not play a major role in capsidiol detoxification.

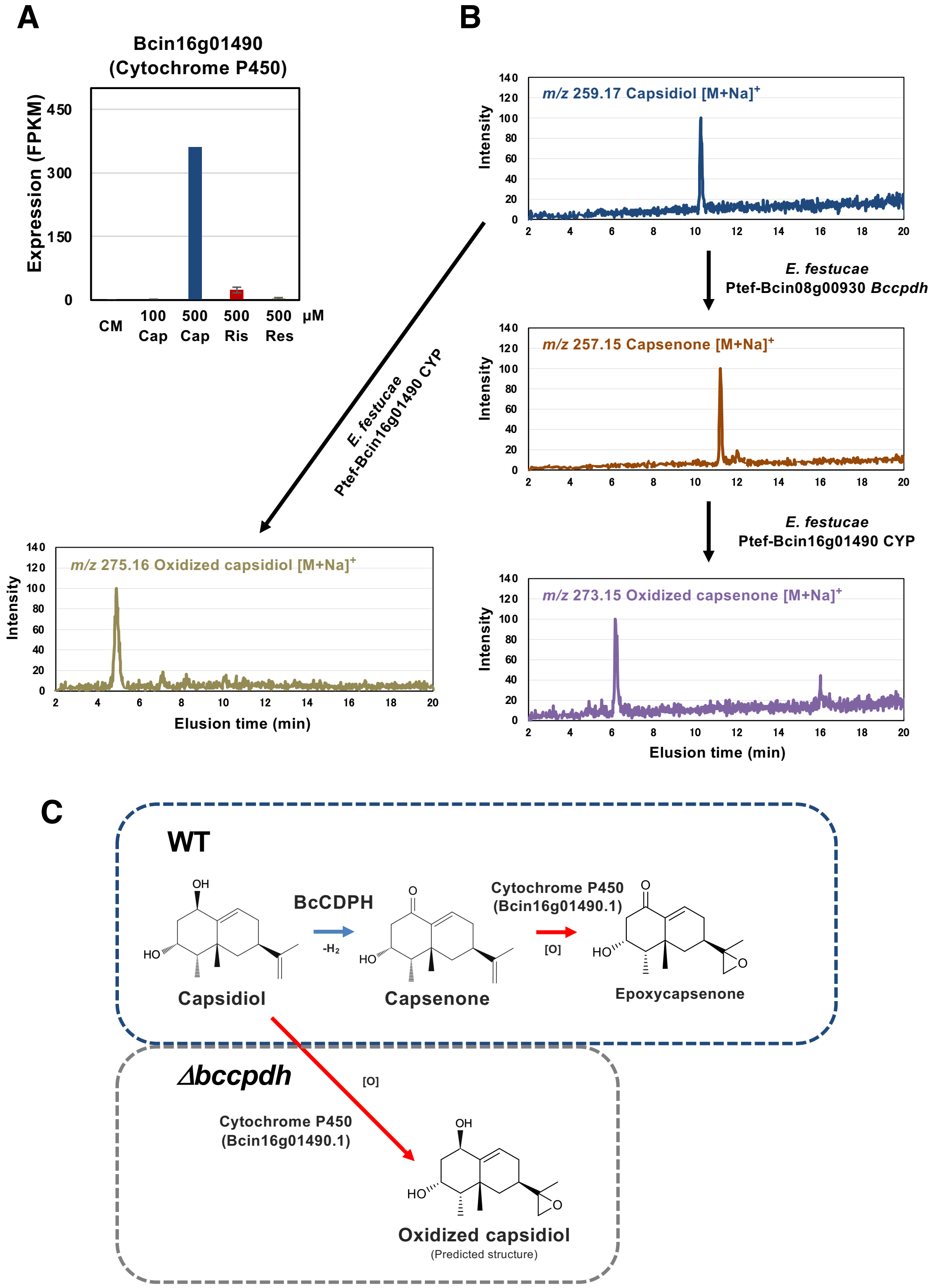


**Fig. S14. (A)** Expression profiles of Bcin16g01490. The gene expression (FPKM value) was determined by RNA-seq analysis of *B. cinerea* cultured in CM media containing 100 µM or 500 µM capsidiol, 500 µM rishitin or 500 µM resveratrol (n = 3) or 500 µM capsidiol (n = 1) for 24 h. Data are mean ± SE. **(B)** Mycelia of *E. festucae* transformant expressing *Bccpdh* was incubated in CM media containing 500 µM capsidiol for 70 h and collected culture filtrate was then incubated with *E. festucae* expressing Bcin16g01490 gene for 5 days. The resultant metabolite was subjected to the structural analysis as shown in Figs. S15-S17. Alternatively, the mycelia of *E. festucae* expressing Bcin16g01490 was incubated in 100 µM capsidiol for 4 days. The metabolite was detected by LC/MS. **(C)** Predicted metabolism of capsidiol in *B. cinerea* wild type (WT) and *Δbccpdh*.

### Supplementary Note 5

#### Chemical analysis of oxidized capsenone.

*E. festucae* transformant expressing Bcin08g00930 (*Bccpdh*) under the control of constitutive TEF prompter (Vanden Wymelenberg *et al.* 1997) was cultured in 10 ml CM media containing 100 μM capsidiol for 70 h and the culture filtrate was collected. The filtrate was sterilized using a syringe filter (pore size 0.45 μm, Millipore) and further incubated with *E. festucae* transformant expressing Bcin16g01490 (encoding a cytochrome P450) for 5 days.

The resultant supernatant was extracted with EtOAc and the extract was analyzed by LC/MS. The major peak appearing at 8.1 min showed the ion peaks of *m/z* 251.1638 (calcd for $C_{15}H_{23}O_3$ $[M+H]^+$: 251.1642) and 273.1455 (calcd for $C_{15}H_{22}O_3Na$ $[M+Na]^+$: 273.1461) (Fig. S15), suggesting the molecular formula to be $C_{15}H_{22}O_3$.

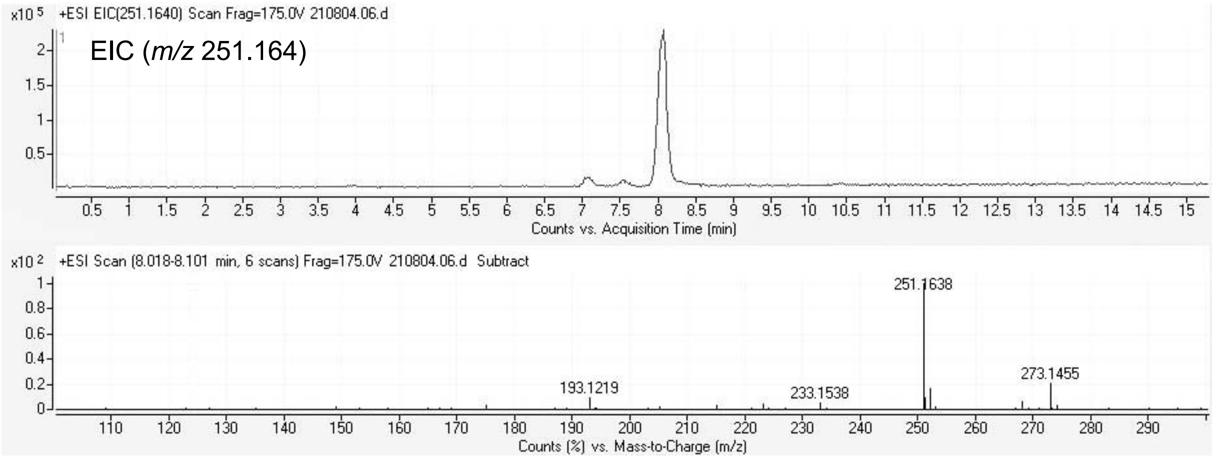


**Fig. S15.** LC/MS analysis of supernatant of *Epichloë* transformant cultured with capsidiol

The extract was further purified by HPLC (Fig. S16) to give the product that possesses the molecular formula of $C_{15}H_{22}O_3$ mentioned above. The molecular formula suggested that this product was formed from capsidiol ($C_{15}H_{24}O_2$) by dehydration (-2H) and oxygen insertion (+O).

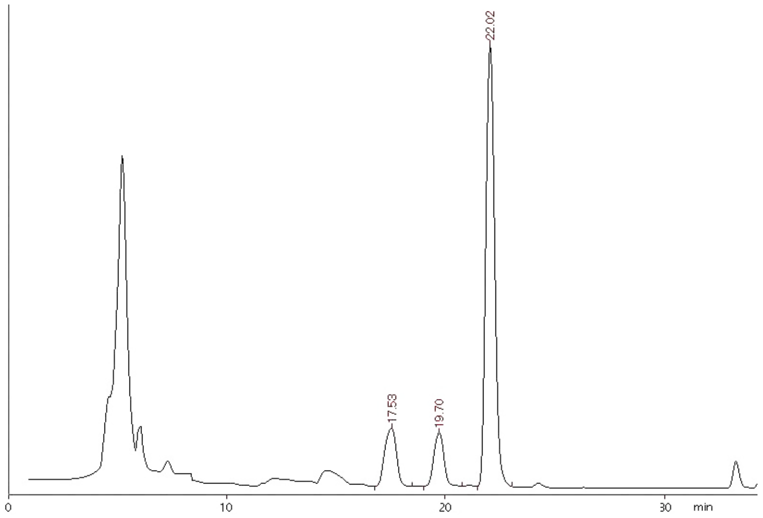


**Fig. S16.** Preparative HPLC of capsidiol metabolites.

The peak at 22 min was found to be capsenone 11,12-epoxide

The structure of the metabolite was determined by two-dimensional NMR analyses. COSY and TOCSY experiments revealed two partial frameworks corresponding to C2-C15 and C6-C9 of capsidiol (Fig. S17). Other components are two singlet methyls (C13 and C14) and a methylene group (C12) as suggested by $^{1}H$ NMR. The lack of the oxy-methine proton (H1) of capsidiol suggests that this position is oxidized to ketone like capsenone, which was supported by the absorption maximum at 250 nm (photodiode array detection in HPLC). The singlet methyl at C14 corresponds to the C14 position of capsidiol due to similar chemical shifts (1.29 and 1.36, respectively). The singlet methyl at C13 (d 1.73) of capsidiol was shifted to the high-field area at d 1.13, and olefinic protons at C12 (d 4.68 and 4.82) of capsidiol largely shifted to the high field area (d 2.58 and 2.68). These facts strongly suggested that the 1,1-disubstituted olefin at C11-C12 in capsidiol is oxidized to epoxide. Therefore, in the light of the molecular formula, we concluded that the metabolite is capsenone 11,12-epoxide as shown in Fig. S17.

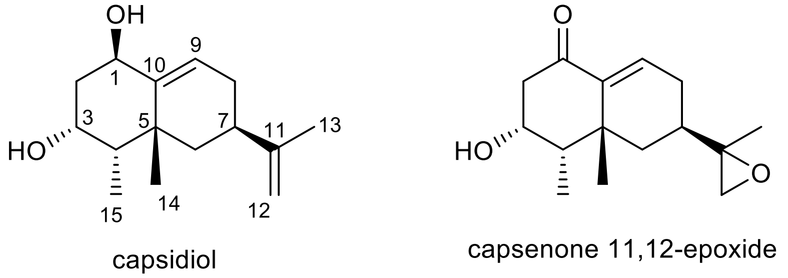


**Fig. S17.** Structures of capsidiol and capsenone 11,12-epoxide (Epoxycapsenone)

### Methods for the structural analysis of oxidized capsenone.

#### General procedure

NMR spectra were investigated on an Avance ARX400 spectrometer (Bruker Bio Spin, Yokohama, Japan). The chemical shifts (ppm) were referenced to the solvent residual peak at $d_H$ 7.26 ppm ($CDCl_3$). LC/MS was measured by a 1100 High-Performance Liquid Chromatography (HPLC) system (Agilent Technologies, Santa Clara, CA) connected to an Agilent 6520 Accurate-Mass Q-TOF spectrometer.

#### Extraction and LC/MS analysis

The supernatant of the culture broth (10 mL) with capsidiol (100 μM) was extracted with EtOAc (10 mL, twice). The organic layers were concentrated and the residual oil was dissolved in MeCN (0.5 mL) to give a stock solution (2 mM equivalent to capsidiol). A portion (2 μl) of the solution was diluted to 1 mL with 50% MeCN and 5 μl was used for LC/MS analysis.

#### Purification of capsenone 11,12-epoxide

The stock solution of the EtOAc extract was concentrated and re-dissolved in 30% MeCN (0.5 mL) and subjected to preparative HPLC [Develosil ODS-UG-5 (10 x 250 mm), 20-50% MeCN (45 min), 3 mL/min, detected at 230 nm] to give capsenone 11,12-epoxide (0.13 mg).

$^{1}H$ NMR ($CDCl_3$, 400 MHz) δ 6.68 (d, J=6.0 Hz, 1H, H-9), 4.47 (m, 1H, H-3), 2.72 (dd, J=16.4, 5.6 Hz, H-2), 2.68 (d, J=4.6 Hz, 1H, H-12), 2.58 (d, J=4.6 Hz, 1H, H-12), 2.35 (dd, J=16.4, 11.6 Hz, 1H, H-2), 2.33 (m, 1H, H-8), 1.99 (brd, J=14.4 Hz, 1H, H-6), 1.92 (m, 1H, H-4), 1.85 (ddd, J=17.0, 11.8, 2.2 Hz, 1H, H-8), 1.57 (m, 1H, H-7), 1.31 (t, J=14.4 Hz, 1H, H-6), 1.29 (s, 3H, H-13), 1.13 (s, 3H, H-14), 1.00 (d, J=7.2 Hz, H-15). ESI-TOF-MS(+) *m/z* 251.1638 (calcd for $C_{15}H_{23}O_3$ $[M+H]^+$: 251.1642), 273.1455 (calcd for $C_{15}H_{22}O_3Na$ $[M+Na]^+$: 273.1461).

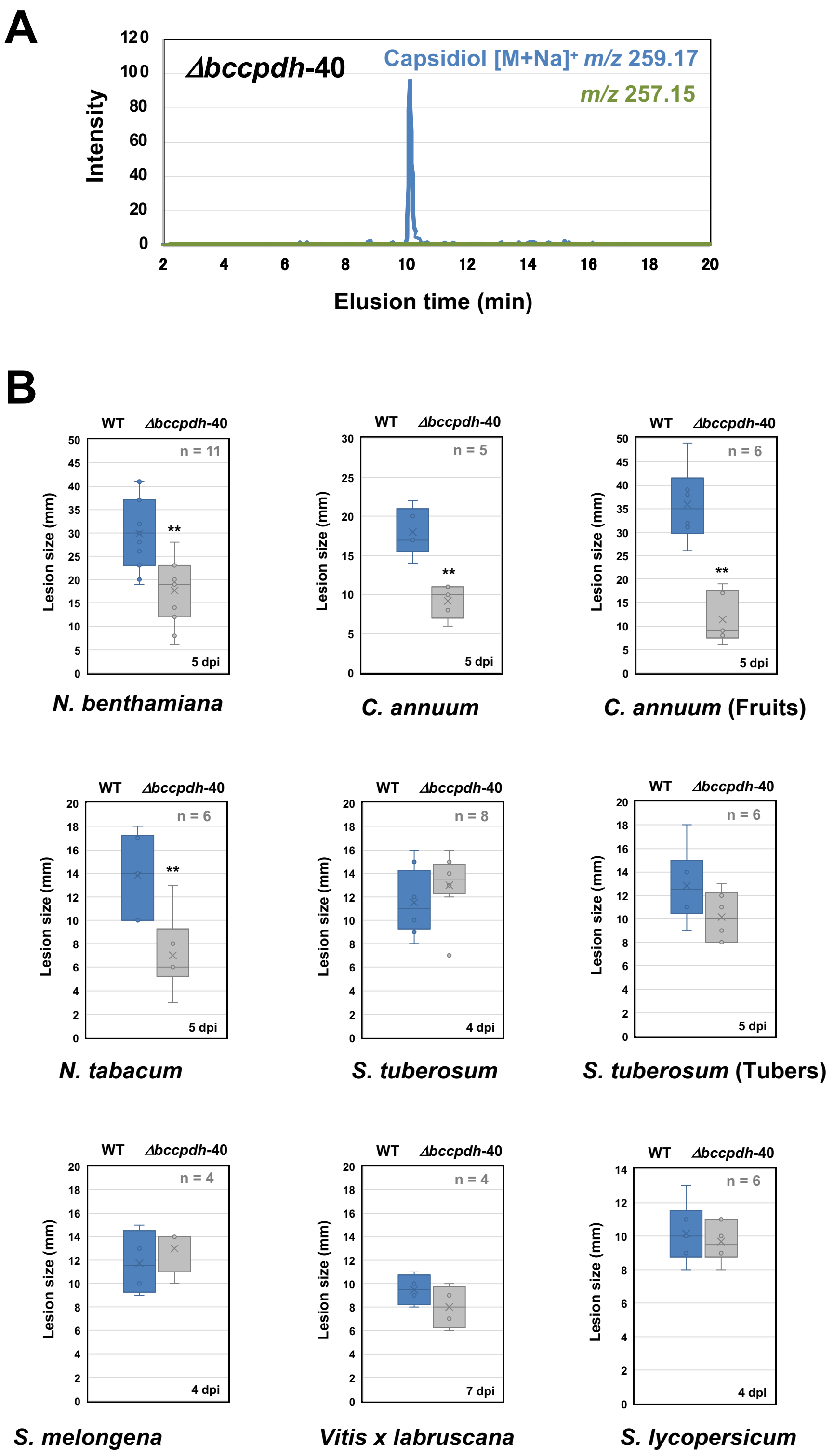


**Fig. S18. (A)** Mycelial blocks (approx. 1 mm$^3$) of *B. cinerea Bccpdh* KO strain (*Δbccpdh*-40) were incubated in 50 µl of 100 µM capsidiol for 4 days and capsidiol, but not capsenone, was detected by LC/MS. **(B)** Indicated plants were inoculated with a mycelial block (5 mm$^3$) of wild type (WT) or *Δbccpdh*-40 and lesion size was measured at 4 to 7 days after the inoculation (dpi). Asterisks indicate a significant difference from WT as assessed by two-tailed Student's *t*-test. $^{**}P < 0.01$. Lines and crosses (x) in the columns indicate the median and mean values, respectively.

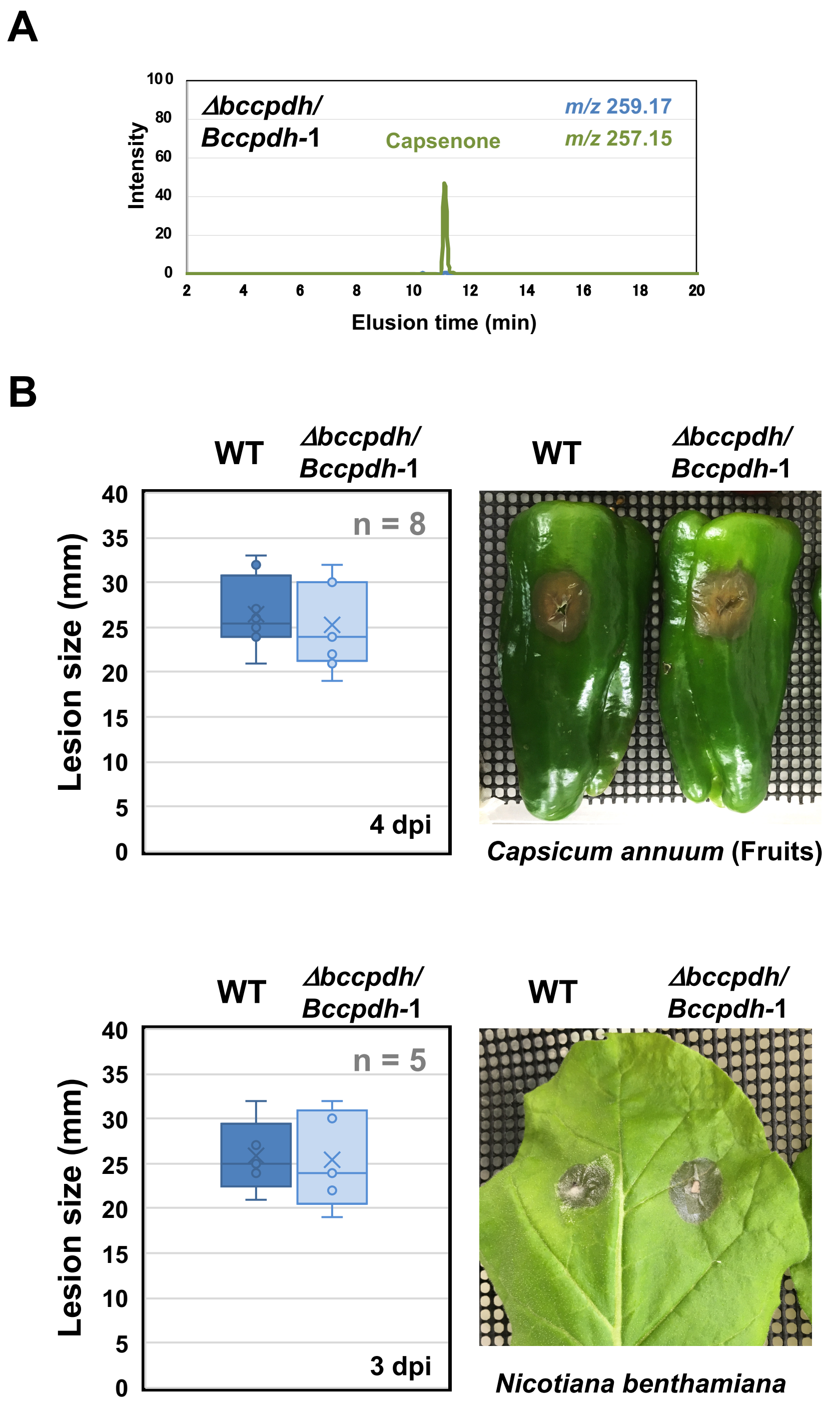


**Fig. S19. (A)** Mycelial blocks (approx. 1 mm$^3$) of *B. cinerea Bccpdh* complemented strain (*Δbccpdh/Bccpdh*-1) were incubated in 50 µl of 100 µM capsidiol for 4 days and capsenone was detected by LC/MS. **(B)** *C. annuum* fruits and *N. benthamiana* leaves were inoculated with a mycelial block (5 mm$^3$) of *B. cinerea* wild type (WT) or *Δbccpdh/Bccpdh*-1 and lesion size was measured at 4 or 3 days after the inoculation (dpi). Lines and crosses (x) in the columns indicate the median and mean values, respectively.

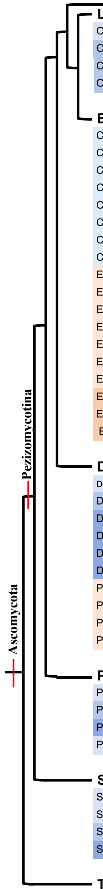


| Taxonomic group | Species | Type | Strain | Orthologue (Gene ID) | Cluster type |
|---|---|---|---|---|---|
| **Lecanoromycetes** | | | | | |
| OSLEUM clade; Umbilicariomycetidae; Umbilicariales | *Lasallia pustulata* | L | A1-1 | | |
| OSLEUM clade; Lecanoromycetidae; Teloschistales | *Xanthoria parietina* | L | 46-1 | | |
| OSLEUM clade; Lecanoromycetidae; Lecanorales | *Usnea florida* | L | ATCC18376 | | |
| OSLEUM clade; Lecanoromycetidae; Lecanorales | *Cladonia grayi* | L | Cgr/DA2myc/ss | | |
| **Eurotiomycetes** | | | | | |
| Chaetothyriomycetidae; Chaetothyriales; Herpotrichiellaceae | *Cladophialophora bantiana* | A | CBS 173.52 | | |
| Chaetothyriomycetidae; Chaetothyriales; Herpotrichiellaceae | *Cladophialophora psammophila* | S | CBS 110553 | A1O5_11311 | A |
| Chaetothyriomycetidae; Chaetothyriales; Herpotrichiellaceae | *Exophiala aquamarina* | A | CBS 119918 | A1O9_12300 | A |
| Chaetothyriomycetidae; Chaetothyriales; Herpotrichiellaceae | *Exophiala spinifera* | A | CBS 89968 | PV08_08948 | A |
| Chaetothyriomycetidae; Chaetothyriales; Herpotrichiellaceae | *Fonsecaea nubica* | A | CBS 269.64 | AYO20_01287 | B |
| Chaetothyriomycetidae; Chaetothyriales; Herpotrichiellaceae | *Fonsecaea monophore* | A | CBS 269.37 | AYO21_01771 | B |
| Chaetothyriomycetidae; Chaetothyriales; Herpotrichiellaceae | *Fonsecaea pedrosoi* | A | CBS 271.37 | Z517_08798 Z517_04127 | A B |
| Chaetothyriomycetidae; Chaetothyriales; Herpotrichiellaceae | *Fonsecaea multimorphosa* | A | CBS 102226 | Z520_02000 | C |
| Eurotiomycetidae; Eurotiales; Aspergillaceae | *Aspergillus nidulans* | S | FGSC A4 | | |
| Eurotiomycetidae; Eurotiales; Aspergillaceae | *Aspergillus fumigatus* | A | A1163 | | |
| Eurotiomycetidae; Eurotiales; Aspergillaceae | *Aspergillus turcosus* | A | HMR AF 23 | CDV55_101651 | D |
| Eurotiomycetidae; Eurotiales; Aspergillaceae | *Penicillium solitum* | P | IBT 29525 | | |
| Eurotiomycetidae; Eurotiales; Aspergillaceae | *Penicillium vulpinum* | S | IBT 29486 | PENVUL_c003G09183 | E |
| Eurotiomycetidae; Eurotiales; Aspergillaceae | *Penicillium coprophilum* | S | IBT 31321 | PENCOP_c001G07223 | E |
| Eurotiomycetidae; Eurotiales; Trichocomaceae | *Talaromyces atroroseus* | S | IBT 11181 | | |
| Eurotiomycetidae; Onygenales; Ajellomycetaceae | *Histoplasma capsulatum* | A | G186AR | | |
| Eurotiomycetidae; Onygenales; Arthrodermataceae | *Trichophyton rubrum* | A | CBS 288.86 | | |
| Eurotiomycetidae; Onygenales; Onygenaceae | *Coccidioides immitis* | A | H538.4 | | |
| **Dothideomycetes** | | | | | |
| Dothideomycetes incertae sedis; Botryosphaeriales; Botryosphaeriaceae | *Diplodia seriata* | P | F98.1 | | |
| Dothideomycetidae; Dothideales; Saccotheciaceae | *Aureobasidium pullulans* | E | EXF-150 | | |
| Dothideomycetidae; Mycosphaerellales; Mycosphaerellaceae | *Cercospora berteroae* | P | CBS538.71 | | |
| Dothideomycetidae; Mycosphaerellales; Mycosphaerellaceae; | *Pseudocercospora fijiensis* | P | CIRAD86 | | |
| Dothideomycetidae; Mycosphaerellales; Mycosphaerellaceae | *Zymoseptoria tritici* | P | IPO323 | | |
| Dothideomycetidae; Mycosphaerellales; Teratosphaeriaceae | *Teratosphaeria nubilosa* | P | CBS 116005 | EJ03DRAFT_364263 | F |
| Pleosporomycetidae; Pleosporales; Pleosporineae | *Alternaria alternata* | P | SRC1lrK2f | | |
| Pleosporomycetidae; Pleosporales; Pleosporineae | *Bipolaris maydis* | P | C5 | | |
| Pleosporomycetidae; Pleosporales; Pleosporineae | *Pyrenophora tritici-repentis* | P | Pt-1C-BFP | | |
| Pleosporomycetidae; Pleosporales; Pleosporineae | *Stemphylium lycopersici* | P | CIDEFI 216 | | |
| **Pezizomycetes** | | | | | |
| Pezizales; Ascobolaceae; Ascobolus | *Ascobolus immersus* | S | RN42 | | |
| Pezizales; Tuberaceae; Tuber | *Tuber melanosporum* | M | Mel28 | | |
| Pezizales; Morchellaceae; Morchella | *Morchella conica* | M | CCBAS932 | | |
| Pezizales; Pyronemataceae; Pyronema | *Pyronema omphalodes* | S | CBS 100304 | | |
| **Saccharomycotina** | | | | | |
| Saccharomycetales; Saccharomycetaceae | *Saccharomyces cerevisiae* | S | EC1118 | | |
| Saccharomycetales; Saccharomycetaceae | *Eremothecium gossypii* | P | ATCC 10895 | | |
| Saccharomycetales; Debaryomycetaceae | *Candida albicans* | A | 12C | | |
| Saccharomycetales; Dipodascaceae | *Yarrowia lipolytica* | S | CLIB122 | | |
| **Taphrinomycotina** | | | | | |
| Schizosaccharomycetes; Schizosaccharomycetales; Schizosaccharomycetaceae | *Schizosaccharomyces pombe* | S | 972h- | | |
| Taphrinomycetes; Taphrinales; Taphrinaceae | *Taphrina wiesneri* | P | JCM 22204 | | |
| Pneumocystidomycetes; Pneumocystidales; Pneumocystidaceae | *Pneumocystis carinii* | A | B80 | | |

| Taxonomic group | Species | Type | Strain | Orthologue (Gene ID) | Cluster type |
|---|---|---|---|---|---|
| **Sordariomycetes** | | | | | |
| Hypocreomycetidae; Glomerellales; Glomerellaceae | *Colletotrichum higginsianum* | P | IMI 349063 | | |
| Hypocreomycetidae; Glomerellales; Glomerellaceae | *Colletotrichum orbiculare* | P | MAFF 240422 | | |
| Hypocreomycetidae; Glomerellales; Plectosphaerellaceae | *Verticillium dahliae* | P | JR2 | | |
| Hypocreomycetidae; Hypocreales; Clavicipitaceae | *Claviceps purpurea* | P | 20.1 | | |
| Hypocreomycetidae; Hypocreales; Clavicipitaceae | *Metarhizium acridum* | I | CQMa 102 | | |
| Hypocreomycetidae; Hypocreales; Clavicipitaceae | *Torrubiella hemipterigena* | I | BCC 1449 | VHEMI04522 | G |
| Hypocreomycetidae; Hypocreales; Clavicipitaceae | *Epichloe festucae* | E | Fl1 | | |
| Hypocreomycetidae; Hypocreales; Cordycipitaceae | *Beauveria bassiana* | I | ARSEF 2860 | | |
| Hypocreomycetidae; Hypocreales; Cordycipitaceae | *Akanthomyces lecanii* | I | RCEF 1005 | LEL_05488 | H |
| Hypocreomycetidae; Hypocreales; Cordycipitaceae | *Cordyceps sp.* | I | RAO-2017 | CDD83_9032 | I |
| Hypocreomycetidae; Hypocreales; Cordycipitaceae | *Cordyceps militaris* | I | CM01 | | |
| Hypocreomycetidae; Hypocreales; Hypocreaceae | *Trichoderma harzianum* | M | T6776 | | |
| Hypocreomycetidae; Hypocreales; Nectriaceae; Fusarium fujikuroi species complex | *Fusarium fujikuroi* | P | EF1 | FFUJ_06845 | A |
| Hypocreomycetidae; Hypocreales; Nectriaceae; Fusarium fujikuroi species complex | *Fusarium verticillioides* | P | 7600 | | |
| Hypocreomycetidae; Hypocreales; Nectriaceae; Fusarium fujikuroi species complex | *Fusarium proliferatum* | P | ET1 | FPRO_07385 | A |
| Hypocreomycetidae; Hypocreales; Nectriaceae; Fusarium fujikuroi species complex | *Fusarium nygamai* | P | CS10214 | FNYG_05967 | A |
| Hypocreomycetidae; Hypocreales; Nectriaceae; Fusarium sambucinum species complex | *Fusarium graminearum* | P | PH-1 | | |
| Hypocreomycetidae; Hypocreales; Nectriaceae; Fusarium oxysporum species complex | *Fusarium oxysporum* f. sp. *melonis* | P | 26406 | | |
| Hypocreomycetidae; Hypocreales; Nectriaceae; Fusarium oxysporum species complex | *Fusarium oxysporum* f. sp. *pisi* | P | HDV247 | FOVG_05603 | J |
| Hypocreomycetidae; Hypocreales; Nectriaceae; Fusarium oxysporum species complex | *Fusarium oxysporum* f. sp. *raphani* | P | 54005 | FOQG_14214 | J |
| Hypocreomycetidae; Hypocreales; Nectriaceae; Fusarium solani species complex | *Fusarium solani* (*F. vanettenii*) | P | 77-13-4 | NechaG7417 | K |
| Hypocreomycetidae; Hypocreales; Nectriaceae; Fusarium solani species complex | *Fusarium kuroshium* | P, IS | UCR3666 | CDV36_000277 | K |
| Hypocreomycetidae; Hypocreales; Nectriaceae; Fusarium solani species complex | *Fusarium ambrosium* | P, IS | NRRL 20438 | CDV31_000895 | K |
| Hypocreomycetidae; Hypocreales; Nectriaceae; Fusarium solani species complex | *Fusarium euwallaceae* | P, IS | UCR1854 | BHE90_006019 | K |
| Hypocreomycetidae; Hypocreales; Ophiocordycipitaceae | *Hirsutella minnesotensis* | I | 3608 | HIM_09377 | I |
| Hypocreomycetidae; Hypocreales; Ophiocordycipitaceae | *Ophiocordyceps sinensis* | I | CO18 | OCS_02677 | I |
| Hypocreomycetidae; Hypocreales; Ophiocordycipitaceae | *Purpureocillium lilacinum* | A | Sc_YJM1199 | VFPBJ_07474 | I |
| Hypocreomycetidae; Hypocreales; Ophiocordycipitaceae | *Tolypocladium paradoxum* | I | NRBC 100945 | TPAR_01618 | A |
| Hypocreomycetidae; Hypocreales; Ophiocordycipitaceae | *Tolypocladium capitatum* | I | CBS 113982 | | |
| Sordariomycetidae; Magnaporthales; Pyriculariaceae | *Pyricularia oryzae* | P | 70-15 | | |
| Sordariomycetidae; Sordariales; Podosporaceae | *Podospora anserina* | S | S mat+ | | |
| Sordariomycetidae; Sordariales; Sordariaceae | *Neurospora crassa* | S | OR74A | | |
| Sordariomycetidae; Sordariales; Sordariaceae | *Sordaria macrospora* | S | k-hell | | |
| Xylariomycetidae; Xylariales; Xylariaceae | *Rosellinia necatrix* | P | W97 | | |
| **Leotiomycetes** | | | | | |
| Erysiphales; Erysiphaceae | *Blumeria graminis* f. sp. *hordei* | P | DH14 | | |
| Erysiphales; Erysiphaceae | *Blumeria graminis* f. sp. *tritici* | P | 96224 | | |
| Erysiphales; Erysiphaceae | *Erysiphe necator* | P | C | | |
| Helotiales; Drepanopezizaceae | *Diplocarpon rosae* | P | DortE4 | | |
| Helotiales; Mollisiaceae | *Phialocephala scopiformis* | E | CBS 120377 | | |
| Helotiales; Ploettnerulaceae | *Rhynchosporium agropyri* | P | 04CH-RAC-A.6.1 | | |
| Helotiales; Hyaloscyphaceae | *Hyaloscypha bicolor* | M | E | | |
| Helotiales; Sclerotiniaceae | *Botrytis cinerea* | P | B05.10 | Bcin08g00930 | L |
| Helotiales; Sclerotiniaceae | *Botrytis tulipae* | P | Bt9001 | | |
| Helotiales; Sclerotiniaceae | *Botrytis elliptica* | P | Be9601 | | |
| Helotiales; Sclerotiniaceae | *Botrytis porri* | P | MUCL3349 | | |
| Helotiales; Sclerotiniaceae | *Botrytis paeoniae* | P | Bp0003 | | |
| Helotiales; Sclerotiniaceae | *Botrytis hyacinthi* | P | Bh0001 | | |
| Helotiales; Sclerotiniaceae | *Botrytis galanthina* | P | MUCL435 | | |
| Helotiales; Sclerotiniaceae | *Sclerotinia sclerotiorum* | P | 1980 UF-70 | | |
| Helotiales; Sclerotiniaceae | *Sclerotinia borealis* | P | F-4128 | | |
| Helotiales; Sclerotiniaceae | *Monilinia fructicola* | P | Mfrc123 | | |
| Helotiales; Amorphothecaceae | *Amorphotheca resinae* | S | ATCC 22711 | | |

**Fig. S20** Distribution of *B. cinerea* CPDH orthologues in Ascomycota fungi. Cluster types were classified based on the conservation of genes around CPDH orthologs in the genome (See Figs. S24 and 25). The types of fungi were categorized as follows. L, Lichen; A, Animal pathogen; S, Saprophyte; P, Plant pathogen, E, Endophyte; M, Mycorrhiza; I, Insect pathogen, IS, Insect symbiont.

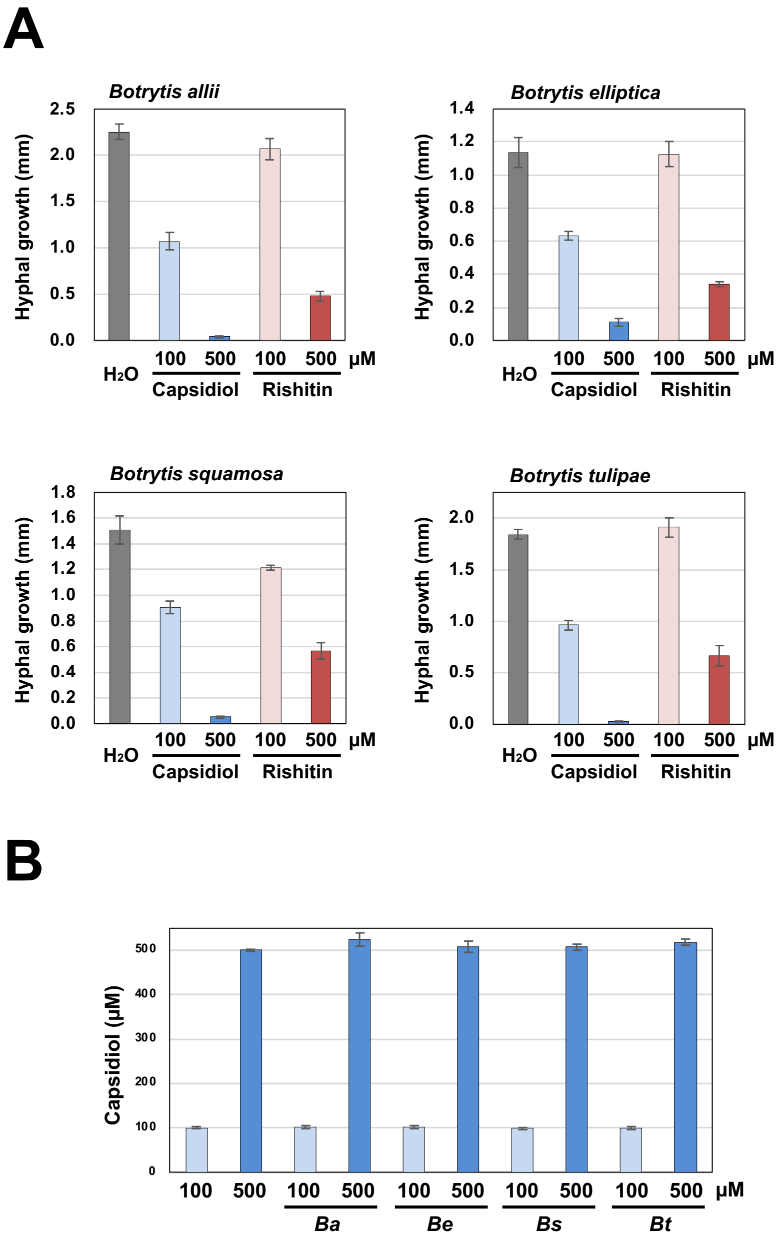


**Fig. S21.** Sensitivity and metabolic capacity of sesquiterpenoid phytoalexins in *Botrytis* species. **(A)** Mycelial blocks (approx. 1 mm$^3$) of the indicated pathogen were incubated in 50 µl water, 100 or 500 µM capsidiol or rishitin. Growth of hyphae from the mycelial block was measured after 24 h incubation (n = 6). **(B)** Residual capsidiol was quantified after 48 h incubation (n = 3). *Ba*, *B. allii* (isolated from onion); *Be*, *B. elliptica* (*Lilium* sp.); *Bs*, *Botrytis squamosa* (Chinese chive); *Bt, B. tulipae* (tulip).

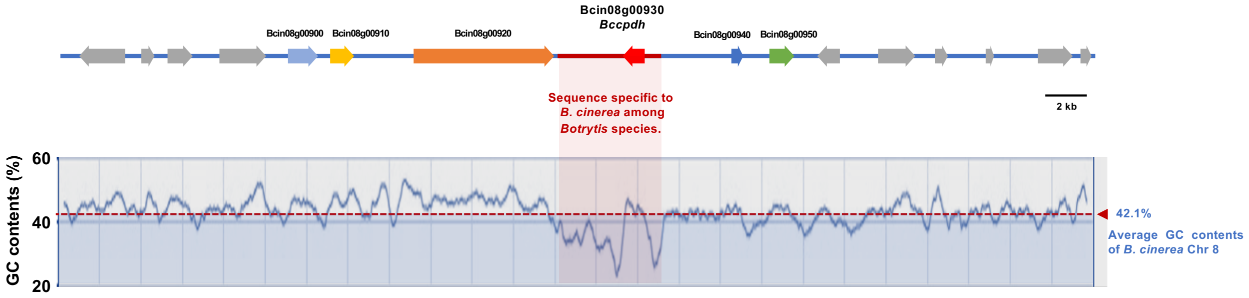


**Fig. S22** GC content plot (window size 500 bp) of *B. cinerea* genomic region surrounding *Bccpdh* gene (Bcin08g00930) in chromosome 8. The average GC content of chromosome 8 (42.1%) is indicated by a dotted red line.

### Supplementary Note 6

#### *CPDH* orthologs in the fungal kingdom

*CPDH* orthologs were found in some Ascomycota fungi. Based on the genes surrounding the *CPDH* orthologs, conserved synteny of the loci was found among different species. Phylogenetic analysis of *CPDH* orthologs indicates that sequence similarity did not necessarily correlate with the taxonomic relationship. Rather, *CPDH* orthologs of the same cluster type tend to form a clade in the phylogenetic tree, which might indicate *CPDH* orthologs (and surrounding genes) were transferred via multiple horizontal gene transfer (HGT) events. For *Fusarium* species, *CPDH* orthologs were detected in species of three species complexes (*F. fujikuroi*, *F. oxysporum* and *F. solani* species complexes), consistent with Stoessl et al. (1973) that reported *F. oxysporum* and *F. solani* can metabolize capsidiol to capsenone. However, cluster types of three species complexes are different, which may indicate that these *Fusarium* species complexes obtained *CPDH* orthologs by independent HGT events. *Bccpdh* locus in *B. cinerea* doesn’t show similarity with other *cpdh* clusters, suggesting that *B. cinerea* might obtain ancestral *Bccpdh* independently from an unidentified organism.

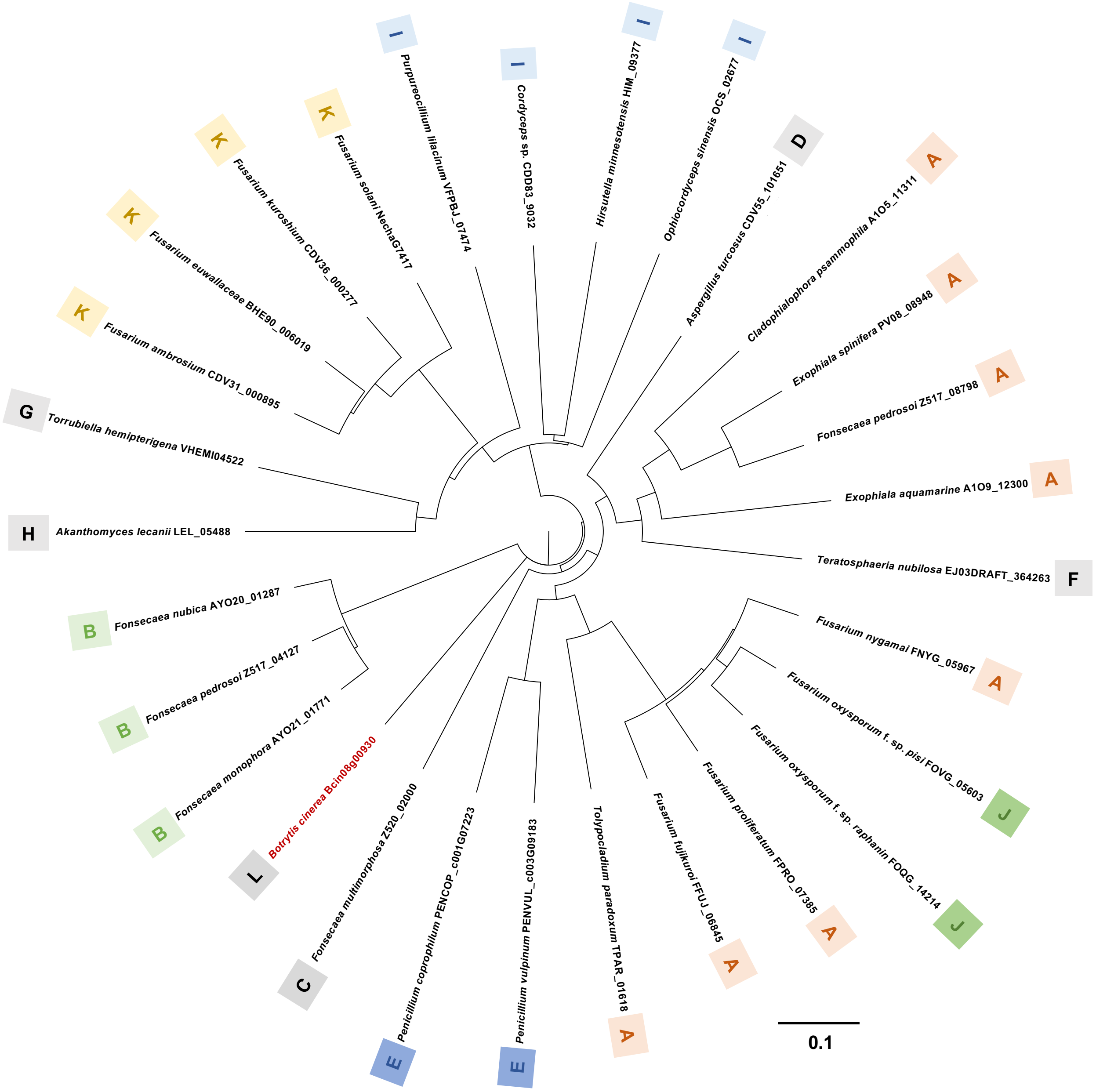


**Fig. S23.** A phylogenetic tree of BcCPDH orthologs from Ascomycota fungi. The deduced amino acid sequences of CPDH orthologs were aligned by ClustalW (Thompson et al., 1994), and the phylogenetic tree was constructed using the neighbor-joining (NJ) method (Saitou and Nei, 1987). The scale bar corresponds to 0.1 estimated amino acid substitutions per site. Cluster types (A to L) classified based on the conservation of genes around CPDH orthologs in the genome are indicated (See Figs. S20, 24 and 25).

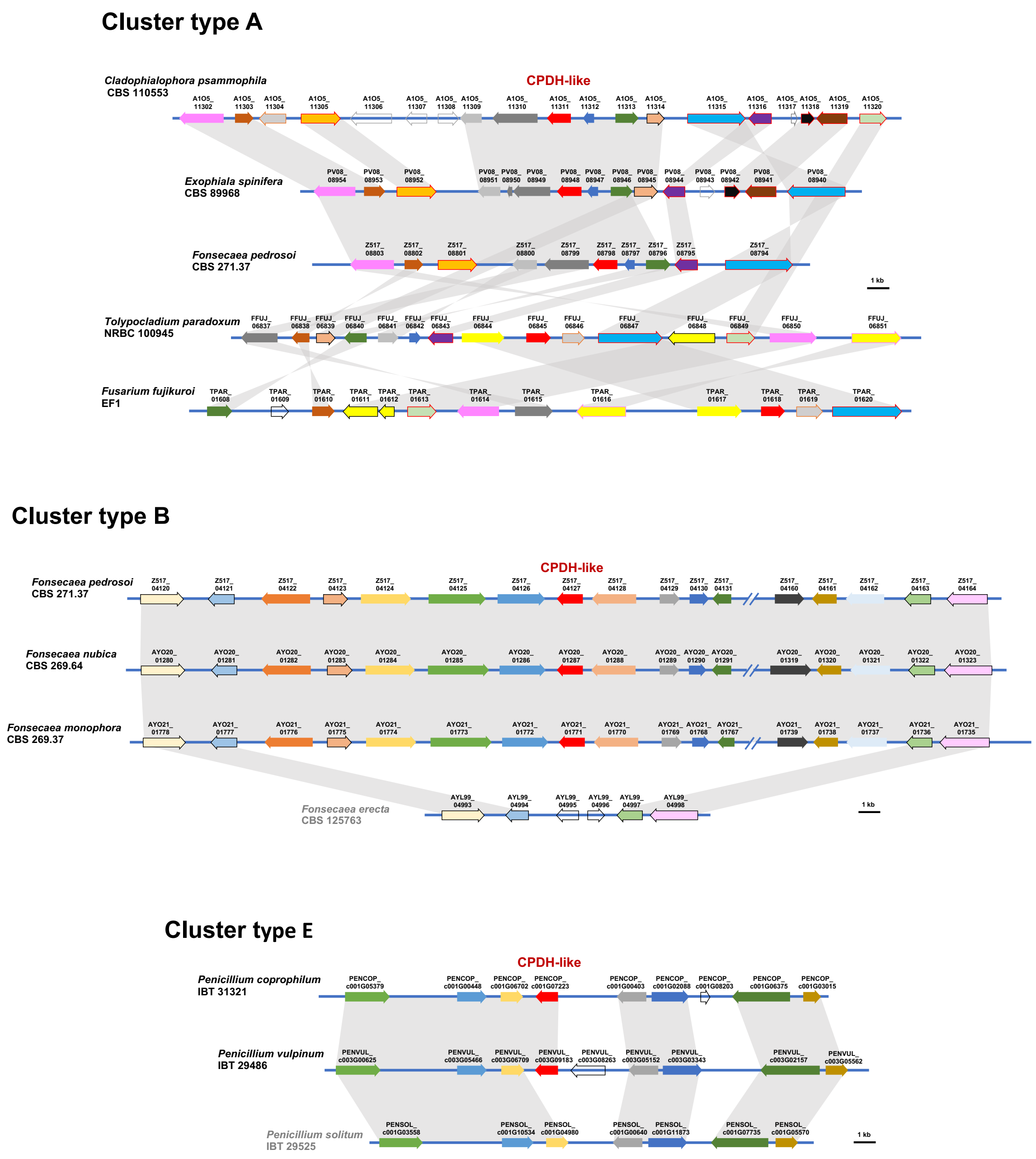


**Fig. S24.** Conserved synteny of the loci containing fugal CPDH orthologues in cluster types A, B and E (See Fig. S20). The matching colors in each cluster type indicate orthologous genes. Genes encoding CPDH orthologues are shown as red arrows. Scale bars = 1 kb.

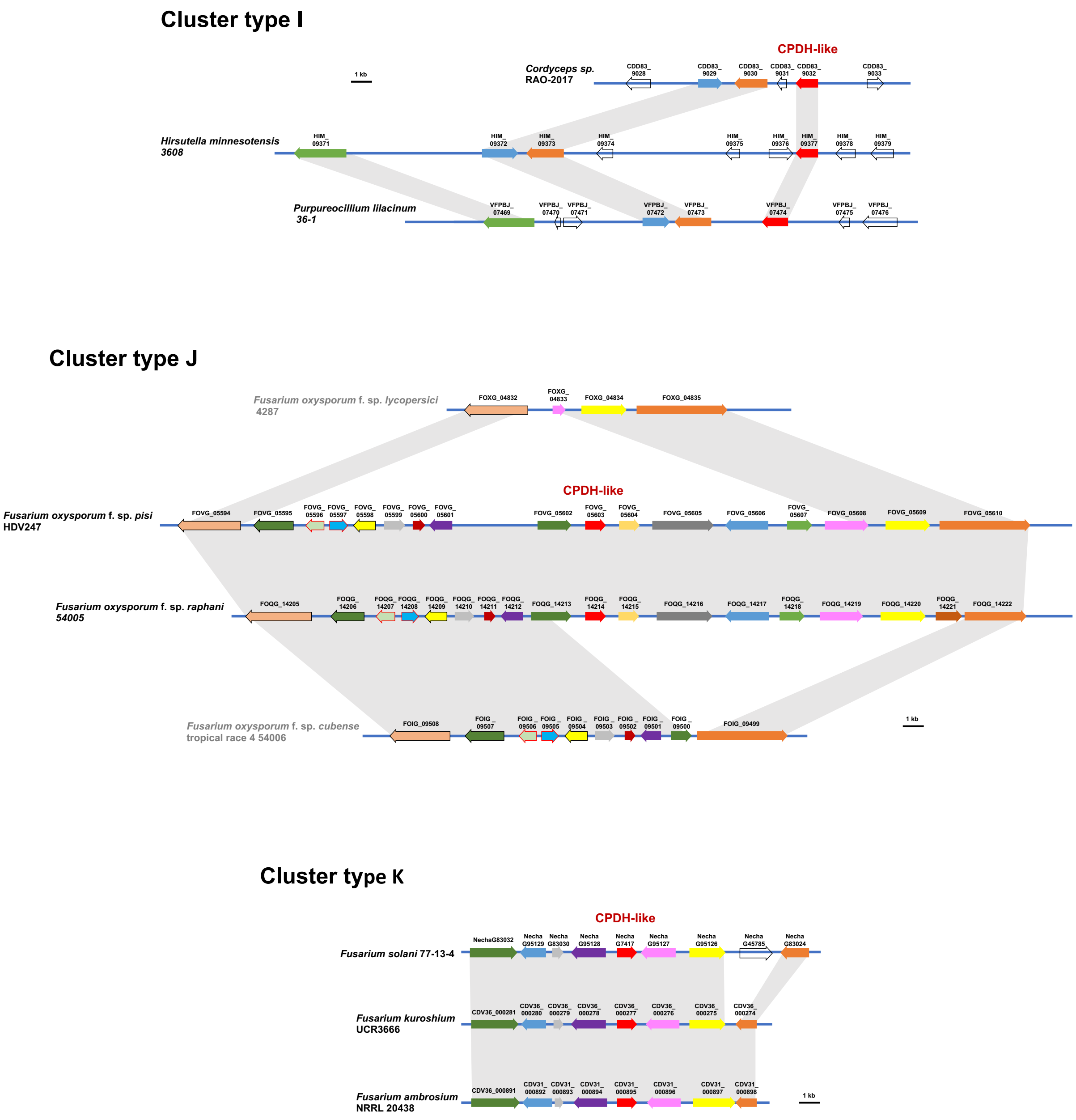


**Fig. S25.** Conserved synteny of the loci containing fugal CPDH orthologues in cluster types I, J and K (See Fig. S20). The matching colors in each cluster type indicate orthologous genes. Genes encoding CPDH orthologues are shown as red arrows. Scale bars = 1 kb.

**Table S5.** CPDH activity in *B. cinerea* strains isolated form different plant species.

| Isolated from | Strain name | Isolated | | CPDH activity |
|---|---|---|---|---|
| | | Location* | Year | |
| Tomato | 2019052 | Mie | 2019 | + |
| Tomato | 2018107 | Mie | 2018 | + |
| Tomato | 2017017 | Mie | 2017 | + |
| Tomato | NBc1 | Aichi | 2009 | + |
| Eggplant | 2019001 | Mie | 2019 | + |
| Eggplant | 2018018 | Mie | 2018 | + |
| Eggplant | 2017017 | Mie | 2017 | + |
| Strawberry | AI18 | Mie | 2018 | + |
| Strawberry | 2019086 | Mie | 2019 | + |
| Strawberry | 2018105 | Mie | 2018 | + |
| Strawberry | 2017100 | Mie | 2017 | + |
| Cucumber | 2019101 | Mie | 2019 | + |
| Cucumber | 2018152 | Mie | 2018 | + |
| Cucumber | 2017149 | Mie | 2017 | + |
| Asparagus | HSB3 | Hokkaido | 2011 | + |
| Lettuce | KBC-2 | Kawaga | 2012 | + |
| Pea | T.K-26-1 | Ibaraki | 1995 | + |
| Rose | T.K-31-3 | Niigata | 1996 | + |
| Barley | TAC96-O1 | Toyama | 1996 | + |
| Bitter orange | S-1-1 | Wakayama | 2003 | + |
| Flowering dogwood | LFP-BB-6 | Fukuoka | 1982 | + |
| Okra | Okurami-2 | Mie | 2007 | + |
| *Morus* sp. | Y-1 | Yamagata | 1980 | + |
| *Vitis* sp. | 4519-1 | Akita | 1986 | + |

*Prefecture name in Japan.

**Table S6.** CPDH activity in *Fusarium oxysporum* strains isolated form different plant species.

| forma specialis | Host plant | Strain name | Isolated | | CPDH activity |
|---|---|---|---|---|---|
| | | | Location* | Year | |
| *cucumerinum* | Cucumber | Cu:8-1 | Toyama | 1990 | + |
| *cucumerinum* | Cucumber | KF-13 | Aomori | 1981 | + |
| *cucumerinum* | Cucumber | 1-19 | Kyoto | 1988 | - |
| *cucumerinum* | Cucumber | JPPAC 10 | Fukuoka | 1978 | + |
| *cucumerinum* | Cucumber | MG1126 | Miyagi | unknown | + |
| *melonis* | Melon | 2-10 | Nagasaki | 1991 | - |
| *melonis* | Melon | B-1 | Kanagawa | 1990 | + |
| *melonis* | Melon | Mel02010 | Nagasaki | unknown | + |
| *melonis* | Hami melon | 2-32 | Yamagata | unknown | - |
| *lagenariae* | Calabash | KF-01 | Aomori | 1978 | + |
| *lagenariae* | Calabash | Lag:4-1 | Wakayama | unknown | + |
| *lagenariae* | Calabash | Lag:6-1 | Kumamoto | 1979 | - |
| *lycopersici* | Tomato | 9859-1 | Aichi | unknown | + |
| *niveum* | Watermelon | 03-05543 | Shizuoka | 1963 | - |
| *niveum* | Watermelon | Niv:1-0 | Fukuoka | unknown | - |
| *niveum* | Watermelon | 80WF-2 | Kagoshima | 1988 | - |
| *momordicae* | Bitter melon | 90NF1-2 | Kagoshima | unknown | - |
| *momordicae* | Bitter melon | 24-11 | Kagoshima | 1994 | - |
| *raphani* | Daikon radish | 03-05123 | unknown | unknown | + |

*Prefecture name in Japan.

### Supplementary Note 7

### Materials and Methods

#### Biological material, growth conditions and incubation in phytoalexins.

Fungal and oomycete strains used in this study were listed in Tables S7, S8 and S9. They were grown on potato dextrose agar (PDA), rye media or V8 agar as indicated in the Tables at 23°C. For the incubation of fungal or oomycete strains in phytoalexins, mycelia blocks (approx. 1 $mm^3$) were excised from the growing edge of the colony on indicated media using a dissection microscope (Stemi DV4 Stereo Microscope, Carl Zeiss, Oberkochen, Germany) and submerged in 50 µl of water or indicated phytoalexin in a sealed 96 well clear plate. The plate was incubated at 23˚C for the indicated time and outgrowth of hyphae was monitored under light microscope BX51 (Olympus, Tokyo, Japan) and measured using ImageJ software (Schneider et al., 2012). Capsidiol, capsidiol 3-acetate and debneyol were purified from *Nicotiana tabacum* as previously reported (Matsukawa *et al.*, 2013) and synthesized rishitin (Murai *et al.* 1975) was provided from former Prof. Akira Murai (Hokkaido University, Japan). Resveratrol and scraleol are obtained from Sigma-Aldrich (Burlington, MA, USA).

**Table S7.** Oomycete and fungal strains used in this study.

| Oomycete and fungal species | Strain | Origin | Media | References |
|---|---|---|---|---|
| **Oomycete species** | | | | |
| *Phytophthora infestans* | 08YB1 | Potato | Rye | Shibata *et al.* 2010 |
| *Phytophthora nicotianae* | Pn96 | Tobacco | V8 | MAFF305940* |
| *Phytophthora capsici* | CH01CMP1 | Green pepper | V8 | MAFF242869* |
| *Phytophthora cryptogea* | CH88-18 | Nipplefruit | V8 | MAFF306435* |
| **Fungal species** | | | | |
| *Alternaria solani* | KL1 | Potato | PDA | MAFF244036* |
| *Colletotrichum coccodes* | PTK1 | Potato | PDA | MAFF243012* |
| *Fusarium coeruleum* | K. Kita 37 | Potato | PDA | MAFF235977* |
| *Gibellulopsis nigrescens* | Kita44 | Potato | PDA | MAFF235985* |
| *Rhizoctonia solani* | NR19 | Potato | PDA | MAFF237435* |
| *Sclerotinia sclerotiorum* | SU-1 | Eggplant | PDA | MAFF744080* |
| *Stemphylium lycopersici* | KuNBY1 | Tobacco | PDA | MAFF306895* |
| *Cercospora nicotianae* | CTC5 | Tobacco | PDA | MAFF243736* |
| *Alternaria brassicicola* | BA31 | Broccoli | PDA | MAFF242993* |
| *Fusarium graminearum* s. str | 407011 | Wheat | PDA | Suga *et al.* 2016 |
| *Fusarium verticillioides* | Maize L-2 | Maize | PDA | MAFF240086* |
| *Botrytis allii* | Yuki11-1 | Onion | PDA | MAFF307143* |
| *Botrytis tulipae* | 4-3 | Tulip | PDA | MAFF245230* |
| *Botrytis squamosa* | 5ND4 | Chinese chive | PDA | MAFF244973* |
| *Botrytis elliptica* | S0210 | *Lilium* sp. | PDA | MAFF306626* |

*MAFF No. of strains obtained from stock center of Ministry of Agriculture, Forestry and Fisheries (MAFF), Japan.

**Table S8.** *Botrytis cinerea* strains used in this study.

| Fungal species | Strain | Host plant | Media | References |
|---|---|---|---|---|
| *Botrytis cinerea* | NBc1 | Tomato | PDA | Tsuge unpublished |
| | 2019052 | Tomato | PDA | Kawakami unpublished |
| | 2018107 | Tomato | PDA | Kawakami *et al.* 2019 |
| | 2017017 | Tomato | PDA | Kawakami *et al.* 2019 |
| | 2019001 | Eggplant | PDA | Kawakami unpublished |
| | 2018018 | Eggplant | PDA | Kawakami *et al.* 2019 |
| | 2017017 | Eggplant | PDA | Kawakami *et al.* 2019 |
| | AI18 | Strawberry | PDA | This study |
| | 2019086 | Strawberry | PDA | Kawakami unpublished |
| | 2018105 | Strawberry | PDA | Kawakami *et al.* 2019 |
| | 2017100 | Strawberry | PDA | Kawakami *et al.* 2019 |
| | 2019101 | Cucumber | PDA | Kawakami unpublished |
| | 2018152 | Cucumber | PDA | Kawakami *et al.* 2019 |
| | 2017149 | Cucumber | PDA | Kawakami *et al.* 2019 |
| | HSB3 | Asparagus | PDA | MAFF243107* |
| | KBC-2 | Lettuce | PDA | MAFF307162* |
| | T.K-26-1 | Pea | PDA | MAFF237249* |
| | T.K-31-3 | Rose | PDA | MAFF237516* |
| | TAC96-O1 | Barley | PDA | MAFF237696* |
| | S-1-1 | Bitter orange | PDA | MAFF306809* |
| | LFP-BB-6 | Flowering dogwood | PDA | MAFF410003* |
| | Okurami-2 | Okra | PDA | MAFF731108* |
| | Y-1 | *Morus* sp. | PDA | MAFF840049* |
| | 4519-1 | Vitis sp. | PDA | MAFF615005* |

*MAFF No. of strains obtained from stock center of Ministry of Agriculture, Forestry and Fisheries (MAFF), Japan.

**Table S9.** *Fusarium oxysporum* strains used in this study.

| Fungal species | forma specialis | Strain | Host plant | Media | References |
|---|---|---|---|---|---|
| *Fusarium oxysporum* | *cucumerinum* | Cu:8-1 | Cucumber | PDA | Namiki *et al.* 1994 |
| | *cucumerinum* | KF-13 | Cucumber | PDA | Kuwata unpublished |
| | *cucumerinum* | 1-19 | Cucumber | PDA | Fukunishi unpublished |
| | *cucumerinum* | JPPAC 10 | Cucumber | PDA | Kiso unpublished |
| | *cucumerinum* | MG1126 | Cucumber | PDA | Honkura unpublished |
| | *melonis* | 2-10 | Melon | PDA | Sakaguchi unpublished |
| | *melonis* | B-1 | Melon | PDA | Namiki *et al.* 1994 |
| | *melonis* | Mel02010 | Melon | PDA | Namiki *et al.* 1994 |
| | *melonis* | 2-32 | Hami melon | PDA | Yuki unpublished |
| | *lagenariae* | KF-01 | Calabash | PDA | Kuwata unpublished |
| | *lagenariae* | Lag:4-1 | Calabash | PDA | Kobayashi unpublished |
| | *lagenariae* | Lag:6-1 | Calabash | PDA | Kobayashi unpublished |
| | *lycopersici* | 9859-1 | Tomato | PDA | Matsusaki unpublished |
| | *niveum* | 03-05543 | Watermelon | PDA | Namiki *et al.* 1994 |
| | *niveum* | Niv:1-0 | Watermelon | PDA | Namiki *et al.* 1994 |
| | *niveum* | 80WF-2 | Watermelon | PDA | Namiki *et al.* 1994 |
| | *momordicae* | 90NF1-2 | Bitter melon | PDA | Namiki *et al.* 1994 |
| | *momordicae* | 24-11 | Bitter melon | PDA | Yamaguchi unpublished |
| | *raphani* | 03-05123 | Daikon radish | PDA | Namiki *et al.* 1994 |

**Quantitative analysis of phytoalexins by GCMS.**

For the quantification of phytoalexins after the incubation with pathogens, the supernatant (50 µl) was collected, mixed with 50 µl ethyl acetate by vortexing for 1 min, and phytoalexin extracted in the organic solvent were collected and quantified by GC/MS using an Agilent Technologies 7890A GC System with a DuraBond Ultra Inert column (length 30 m; diameter 0.25 mm; film 0.25 µm, Agilent Technologies, Santa Clara, CA, USA) as previously described (Camagna *et al.* 2020). Pure capsidiol and rishitin were used for quantitative standards.

**Detection of phytoalexins and their metabolites using LC/MS.**

For the detection of phytoalexins and their metabolites after the incubation with pathogens, the supernatant (50 µl) was collected, mixed with 50 µl acetonitrile and measured by LC/MS (Accurate-Mass Q-TOF LC/MS 6520, Agilent Technologies) with ODS column Cadenza CD-C18, 75 x 2 mm (Imtakt, Kyoto, Japan).

**Extraction of RNA and RNAseq analysis.**

Mycelial blocks (approx. 1 $mm^3$, 100 pieces) were incubated in 10 ml of CM media [1 g $Ca(NO_3)^2$, 0.2 g $KH_2PO_4$, 0.25 g $MgSO_4$, 0.15 g NaCl, 500 µl Micronutrient solution (Sanderson and Srb 1965), 1 g yeast extract, 1 g peptone /1L] with or without indicated concentration of phytoalexins at 23°C for 24 h with gentle shaking (100 rpm) and frozen in liquid nitrogen. The frozen mycelia were ground using mortar and pestle, and the total RNA was extracted using the RNeasy Plant Mini Kit (QIAGEN, Hilden, Germany), according to the manufacturer's instructions. The quality and quantity of isolated RNA were evaluated using Qubit RNA HS Assay Kit (Thermo Fisher Scientific, Waltham, MA, USA). The mRNA was purified with NEBNext Poly(A) mRNA magnetic isolation module (New England Biolabs, Ipswich, MA, USA) and used for the construction of cDNA libraries using the NEBNext Ultra II RNA library prep kit for Illumina and NEBNext Multiplex oligos for Illumina (New England Biolabs) according to the manufacturer's instructions. RNA-Seq libraries were sequenced using Illumina NextSeq 500 (Illumina, San Diego, CA, USA) with single-read mode. The nucleotides of each read with less than 13 quality value were masked and reads less than 50 bp in length were discarded before mapping. The filtered reads were mapped to annotated cDNA sequences for *B. cinerea* (Botrytis_cinerea.ASM83294v1.cdna.all.fa, http://fungi.ensembl.org/Botrytis _cinerea/Info/Index) using Bowtie software (Langmead *et al.* 2009) and the number of reads mapping to each annotated cDNA was counted. For each gene, the relative fragments per kilobase of transcript per million mapped reads (FPKM) values were calculated and significant difference from the control was assessed by the two-tailed Student's *t*-test. RNA-seq data reported in this work are available in GenBank under the accession numbers DRA013980.

**Extraction of genomic DNA, PCR and construction of vectors**

Genomic DNA of *E. festucae* and *B. cinerea* was isolated from fungal mycelium grown in potato dextrose broth (PDB) as described previously (Byrd *et al.,* 1990) or using DNeasy Plant Mini Kit (QIAGEN). PCR amplification from genomic and plasmid DNA templates was performed using PrimeStar Max DNA polymerase (Takara Bio, Kusatsu, Japan) or GoTaq Master Mix (Promega, Madison, WI, USA). Vectors for heterologous expression, detection of promoter activity, gene knock out and complementation used in this study are listed in Table S10. Sequences of primers used for the construction of vectors and PCR to confirm the gene knockout are listed in Table S11.

**Table S10.** Plasmids used in this study

| Vector name | Base vector | Restriction sites used | Insert | Primers used to amplify insert | References | Note |
|---|---|---|---|---|---|---|
| **Base vectors** | | | | | | |
| pPN94 | - | - | - | - | Takemoto *et al.* 2006 | Base vector for gene expression under TEF promoter, $Amp^R/Hyg^R$ |
| pNPP150 | - | - | - | - | Niones and Takemoto 2015 | Base vector for gene knockout (*HSVtk* marker), $Amp^R/Gen^R$ |
| pNPP170 | pNPP150 | *Eco* RI/ *Sma* I | P_*BcactA* (Bcin16g02020) | IF-BcPact-F, IF-BcPact-R2 | This study | Base vector for expression of gene under the control of *BcactA* promoter. Base vector for the construction of pNPP170-BcGFP, $Amp^R/Hyg^R$ |
| pNPP170-BcGFP | pNPP170 | *Not* I | BcGFP[1] | IF pNPP170-BcGFP-F, IF pNPP170-BcGFP-R | This study | Base vector for the promoter analysis using BcGFP marker, $Amp^R/Hyg^R$ |
| pNPP210 | pNPP170-BcGFP | *Eco* RI | OflucBK[2] | IF-OflucBK-F IF-OflucBK-R | This study | Base vector for promoter analysis using Luciferase marker, $Amp^R/Hyg^R$ |
| pSF17.1 | - | | - | - | Tanaka *et al.* 2008 | Base vector for complementation, $Amp^R/Gen^R$ |
| **Plasmids for heterologous expression of *B. cinerea* gene in *E. festucae*** | | | | | | |
| pNPP196 (pPN94-Bcin08g00930) | pPN94 | *Bam* HI/ *Not* I | Bcin08g00930 | pPN94-Bc08g00930-F, pPN94-Bc08g00930-R | This study | Constitutive expression of Bcin08g00930 (*Bccpdh*) |
| pNPP197 (pPN94-Bcin12g01750) | pPN94 | *Bam* HI/ *Not* I | Bcin12g01750 | pPN94-Bc12g01750-F, pPN94-Bc12g01750-R | This study | Constitutive expression of Bcin12g01750 |
| pNPP201 (pPN94-Bcin16g01490) | pPN94 | *Bam* HI/ *Not* I | Bcin16g01490 | pPN94-p450-Ch16-F, pPN94-p450-Ch16-R | This study | Constitutive expression of Bcin16g01490 |
| **Plasmids for promoter analysis in *B. cinerea*** | | | | | | |
| pNPP202 (pNPP170-P_Bcin08g00930-BcGFP) | pNPP170-BcGFP | *Sac* I/ *Sma* I | P_*Bccpdh* (1 kb) | pNPP170-P_8g00930-F, pNPP170-P_8g00930-R | This study | Expression of GFP under the control of 1 kb *Bccpdh* promoter |
| pNPP203 (pNPP170-P_Bccpdh(750)-BcGFP) | pNPP170-BcGFP | *Sac* I/ *Sma* I | P_*Bccpdh* (750 bp) | pNPP170-P_8g00930-750F, pNPP170-P_8g00930-R | This study | Expression of GFP under the control of 750 bp *Bccpdh* promoter |
| pNPP204 (pNPP170-P_Bccpdh(500)-BcGFP) | pNPP170-BcGFP | *Sac* I/ *Sma* I | P_*Bccpdh* (500 bp) | pNPP170-P_8g00930-500F, pNPP170-P_8g00930-R | This study | Expression of GFP under the control of 500 bp *Bccpdh* promoter |
| pNPP205 (pNPP170-P_Bccpdh(250)-BcGFP) | pNPP170-BcGFP | *Sac* I/ *Sma* I | P_*Bccpdh* (250 bp) | pNPP170-P_8g00930-250F, pNPP170-P_8g00930-R | This study | Expression of GFP under the control of 250 bp *Bccpdh* promoter |
| pNPP206 (pNPP170-P_Bccpdh(200)-BcGFP) | pNPP170-BcGFP | *Sac* I/ *Sma* I | P_*Bccpdh* (200 bp) | pNPP170-P_8g00930-200F, pNPP170-P_8g00930-R | This study | Expression of GFP under the control of 200 bp *Bccpdh* promoter |
| pNPP207 (pNPP170-P_Bccpdh(100)-BcGFP) | pNPP170-BcGFP | *Sac* I/ *Sma* I | P_*Bccpdh* (100 bp) | pNPP170-P_8g00930-100F, pNPP170-P_8g00930-R | This study | Expression of GFP under the control of 100 bp *Bccpdh* promoter |
| pNPP211 (pNPP210-P_Bccpdh(250)-Luc) | pNPP210 | *Eco* RI | P_*Bccpdh* (250 bp) | pNPP210-P_8g00930-250F, pNPP210-P_8g00930-R | This study | Expression of Luciferase under the control of 250 bp *Bccpdh* promoter |
| **Plasmids for gene knockout and complimentation of *Bccpdh*** | | | | | | |
| pNPP198 (pNPP150-Bccpdh-KOver5) | pNPP150 | *Sal* I/ EcoRI | 5'*Bccpdh*-PtrpC-hph-3'*Bccpdh* | cpdh-KO5-Fo, cpdh-KO5-Ro, cpdh-KO3-F5, cpdh-KO3-R5 | This study | Knockout vector for *Bccpdh* |
| pNPP199 (pSF17-*Bccpdh*) | pSF17.1 | *Eco* RV | *Bccpdh* locus (2 kb) | pSF17-BcCPDH-F pSF17-BcCPDH-R | This study | Complimentation vector of *Bccpdh* |

[1] *GFP* gene synthesized to optimize codon usage for *B. cinerea* (Leroch *et al.* 2011).

[2] *Luciferase* gene synthesized to optimize codon usage for fungi (Gooch *et al.* 2008). Modified to remove a restriction site (Murata *et al.* unpublished).

**Table S11.** Primers used in this study.

| Primer name | Sequence 5'→3' |
|---|---|
| **Primers for sequencing and comformation of gene knockout** | |
| pII99-3 | GGCTGGCTTAACTATGCG |
| PtrpC-2 | CAAATTTTGTGCTCACCG |
| hph-seqR | ACTTCGAGCGGAGGCATC |
| Ptef-seq | TAACCTCTCTTCAGAAAG |
| TtrpC-seq | TCTGGAAGAGGTAAACCCG |
| BcGFP-seqR | CTTATGGCCATTGACGTCAC |
| cpdh-RC-F | CAGCACTTTGAGCTGATACG |
| cpdh-RC-R | AGTTCCTAAAGTTGTAAAGCC |
| cpdh-CF3 | GGCTCTCATCAAGGATATCC |
| **Primers for construction of Base vectors** | |
| IF-BcPact-F | TACCGAGCTCGAATTCGATGTGCGTCCTCTTCTGC |
| IF-BcPact-R2 | ACGTTAAGTGCGGCCGCGGTTGATAAATTAAGACG |
| IF pNPP170-BcGFP-F | TTATCAACCGCGGCCCCCGGGTTCACCATGGTTTC |
| IF pNPP170-BcGFP-R | ACGTTAAGTGCGGCCGAATTCCTATTTGTAAAGTT |
| IF-OflucBK-F | TACCGAGCTCGAATTCATGGAGGACGCCAAGAACA |
| IF-OflucBK-R | AAGTGCGGCCGAATTTCAGAGCTTGGACTTGCCGC |
| **Primers for construction of vectors for heterologous expression** | |
| pPN94-Bc08g00930-F | AACCTCTAGAGGATCATGGCAGCACTATCACTCAA |
| pPN94-Bc08g00930-R | ACGTTAAGTGCGGCCCTAAAGTTGTAAAGCCTGAA |
| pPN94-Bc12g01750-F | AACCTCTAGAGGATCCGATGAACTCCATTACAGCT |
| pPN94-Bc12g01750-R | ACGTTAAGTGCGGCCTCATGAAGTTCTCAACGTCC |
| pPN94-p450-Ch16-F | AACCTCTAGAGGATCATGTCGCCAGCACTCTTCGA |
| pPN94-p450-Ch16-R | ACGTTAAGTGCGGCCCTCCTCTCCAACTTTTAGGC |
| **Primers for construction of vectors for promoter analysis** | |
| pNPP170-P_8g00930-F | CCAAGCTGGGTACCGCTAGACACCTTCTTGGAACA |
| pNPP170-P_8g00930-R | AACCATGGTGAACCCTTCGATTGCTTTCAATAGTG |
| pNPP170-P_8g00930-750F | CCAAGCTGGGTACCGTATAAAAAAAGTATGAATTG |
| pNPP170-P_8g00930-500F | CCAAGCTGGGTACCGGCTTCCCTCTAAATGCTTCA |
| pNPP170-P_8g00930-250F | CCAAGCTGGGTACCGGTGATAACTTATGATTAAGT |
| pNPP170-P_8g00930-200F | CCAAGCTGGGTACCGGACCGCCAAGAAGTAGACAT |
| pNPP170-P_8g00930-100F | CCAAGCTGGGTACCGCCATAATATCTTATGAGTTT |
| pNPP210-P_8g00930-250F | TACCGAGCTCGAATTGTGATAACTTATGATTAAGT |
| pNPP210-P_8g00930-R | CGTCCTCCATGAATTTTCGATTGCTTTCAATAGTG |
| **Primers for construction of knockout and complimentation vectors** | |
| cpdh-KO5-Fo | ATGCCTGCAGGTCGAAGGATAATAGCGGCGTATGA |
| cpdh-KO5-Ro | ATCCTCTAGAGTCGAATCTAAGCCCCCAAGCTTCT |
| cpdh-KO3-F5 | TACCGAGCTCGAATTGTCGTAACTTCCTCAAACTC |
| cpdh-KO3-R5 | TATCATCGATGAATTTGTAAAGCCTGAACAGGAGC |
| pSF17-BcCPDH-F | GAATTCATCGATGATCTAGACACCTTCTTGGAACA |
| pSF17-BcCPDH-R | ACCGGCAGATCTGATTCCTAAAGTTGTAAAGCCTG |

Extension sequence for in-fusion reaction are shown in red letters.

**Fungal transformation**

Protoplasts of *E. festucae* were prepared as follows. Mycelial blocks of *E. festucae* (approx. 1 $mm^3$, 100 pieces) were added to 50 ml PDB media in 100 ml Erlenmeyer flask and shaken for 3 to 4 days at 23˚C, 100 rpm. Mycelia from 3 flasks were collected by centrifugation at 3,000 x g for 10 min and suspended in 30 ml of OM buffer [1.2 mM $MgSO^4$, 10 mM phosphate buffer, pH5.8]. Mycelia were then collected by filtration using an 80-mesh nylon cloth, and suspended in 10 ml of enzyme solution [10 mg/ml lysing Enzymes (Sigma-Aldrich), 5 mg/ml Kitalase (Wako Pure Chemicals) in OM buffer] in 50 ml falcon tube and shaken at 28°C, 80 rpm for approx. 3 h. After removing undigested mycelia by filtration with a 200 mesh nylon cloth, 30 ml of 0.7 M NaCl was added and the protoplasts were precipitated by centrifugation at 3,000 x g for 5 min. The precipitated protoplasts were suspended in 20 ml of STC [1 M sorbitol, 50 mM Tris-HCl (pH 8.0), 50 mM $CaCl_2$] and the solution was centrifuged at 3,000 x g for 5 min. The precipitated protoplasts were resuspended in the STC solution to approx. 2.5 × $10^8$ protoplasts/ml and mixed with 40% PEG solution [40% polyethylene glycol 4000 (Wako Pure Chemicals), 1 M sorbitol, 50 mM Tris-HCl (pH 8.0), 50 mM $CaCl_2$,] at 4:1. Aliquoted protoplast solution (100 µl, 2 x $10^8$ /ml) was stored at -80°C until use.

Protoplasts of *B. cinerea* were prepared as follows. To induce the sporulation of *B. cinerea*, colonies grown on PDA in 90 mm Petri dishes were exposed to BLB blacklight (Peak wavelength 352 nm) for approx. 2 weeks. Sterile water (5-10 ml) was added to the Petri dishes and spores were released using a spreader from the mycelial surface. Spores (approx. 2 x $10^6$) were added to 50 ml PDB media in 100 ml Erlenmeyer flask and shaken for 16 h at 23˚C, 100 rpm. Germinated hyphae were collected by centrifugation at 3,000 x g for 5 min, suspended in 20 ml of 0.7 M NaCl, and centrifuged at 3,000 x g for 5 min. The collected hyphae from 2 flasks were suspended in 5 ml of enzyme solution [10 mg/ml lysing Enzymes (Sigma-Aldrich), 5 mg/ml Kitalase (Wako Pure Chemicals) in 0.7 M NaCl] and shaken at 28°C, 80 rpm for approx. 3 h. After removing undigested mycelia by filtration with a 200 mesh nylon cloth, 15 ml of 0.7 M NaCl was added and the protoplasts were precipitated by centrifugation at 3,000 x g for 5 min. The protoplasts were suspended in 20 ml of STC and the solution was centrifuged at 3,000 x g for 5 min. The precipitated protoplasts were resuspended in the STC solution to approx. 2.5 × $10^8$ (or lower) protoplasts/ml and mixed with 40% PEG solution at 4:1. Aliquoted protoplast solution (100 µl, 2 x $10^8$ /ml or lower) was stored at -80°C until use.

Protoplasts of *E. festucae* or *B. cinerea* (100 µl) were mixed with 5 µg of either circular or linear (for gene KO) plasmids (<100 µl) and incubated on ice for 30 min. The mixture of protoplasts and plasmid DNA was gently mixed with 900 µl of PEG solution and further incubated on ice for 20 min. Aliquots (100 µl) of the protoplast suspension were mixed with 3 ml of 0.8% YPSA media [0.1% yeast extract, 0.1% tryptone, 34.2% sucrose, 0.8% agar] melted and warmed to 50°C, and immediately poured into 90 mm Petri dishes containing approx. 10 ml of YPSA media (1.8% agar). Plates were incubated overnight at 23°C and overlaid with melted (and then cooled to 50°C) PDA containing 150 µg/ml (for *E. festucae*) or 75 µg/ml (for *B. cinerea*) hygromycin B or 400 µg geneticin (for *B. cinerea*). Plates were incubated at 23°C until colonies emerged, which were sub-cultured on PDA containing appropriate antibiotics.

For the isolation of *B. cinerea* knockout strains, candidate colonies were exposed to BLB blacklight for the induction of sporulation. Single spore isolation was performed to obtain purified knockout strains. Note that *Δbccpdh*-40 and -52 were isolated from separate transformation experiments. Transformants of *E. festucae* and *B. cinerea* used in this study are listed in Table S12.

**Table S12.** Transformants used in this study.

| Strains | **Relevant characteristics** | **References** |
|---|---|---|
| ***Epichloë festucae*** | | |
| WT-DsRed | Fl1/pNPP94-DsRed; $Hyg^R$ | Kayano et al., 2013 |
| Ef-Bcin08g00930 | Fl1/pNPP196 ; $Hyg^R$ | This study |
| Ef-Bcin12g01750 | Fl1/pNPP197 ; $Hyg^R$ | This study |
| Ef-Bcin16g01490 | Fl1/pNPP201 ; $Hyg^R$ | This study |
| ***Botrytis cinerea*** | | |
| P_*Bccpdh:GFP* | AI18/pNPP202 ; $Hyg^R$ | This study |
| P_*Bccpdh* (750)*:GFP* | AI18/pNPP203 ; $Hyg^R$ | This study |
| P_*Bccpdh* (500)*:GFP* | AI18/pNPP204 ; $Hyg^R$ | This study |
| P_*Bccpdh* (250)*:GFP* | AI18/pNPP205 ; $Hyg^R$ | This study |
| P_*Bccpdh* (200)*:GFP* | AI18/pNPP206 ; $Hyg^R$ | This study |
| P_*Bccpdh* (100)*:GFP* | AI18/pNPP207 ; $Hyg^R$ | This study |
| P_*Bccpdh* (250)*:Luc* | AI18/pNPP211 ; $Hyg^R$ | This study |
| *Δbccpdh* -52 | Fl1/*Δbccpdh* ::P*trpC* -*hph* ; $Hyg^R$ | This study |
| *Δbccpdh* -40 | Fl1/*Δbccpdh* ::P*trpC* -*hph* ; $Hyg^R$ | This study |
| *Δbccpdh/Bccpdh* -1 | *Δbccpdh* -52/pNPP199; $Hyg^R$, $Gen^R$ | This study |

**Pathogen inoculation**

Leaves, fruits (bell pepper) or tuber (potato) of plant species were kept in moistened and sealed in a plastic chamber. Leaves detached from the plant were covered with a wet Kimwipes at the cut end of the stem. Mycelial blocks (approx. 5 $mm^3$) of *B. cinerea* were excised from the growing edge of the colony grown on PDA and placed on the downside of the leaf or on the fruit and tuber and covered with wet lens paper. For the inoculation on bell pepper fruits, the surface of the fruits was injured by a needle beneath the placed mycelial block. For the inoculation on *N. benthamiana*, mycelial blocks of *B. cinerea* were placed on the upside of leaves attached to the plant body and the plant was kept at high humidity at 23 ˚C for 1 day after the inoculation, and then moved to a growth room at 23 ˚C.

For the inoculation of *B. cinerea* spores (Fig. 4), *B. cinerea* spore suspension (1 x $10^4$/ml) in glucose-phosphate solution (10 mM glucose, 10 mM $NaH_2PO_4$) was placed on the downside of *N. benthamiana* leaves in sealed plastic chambers, covered with wet lens paper, and incubated at 23 ˚C for indicated time.

**Microscopy**

Images of *B. cinerea* strains expressing GFP or hyphae stained with Calcofluor white (Sigma-Aldrich) were captured using a confocal laser scanning microscope FV1000-D (Olympus, Japan). The laser for detection of GFP was used as the excitation source at 488 nm, and GFP fluorescence was recorded between 515 and 545 nm. The laser for detection of Calcofluor white was used as the excitation source at 405 nm, and fluorescence was recorded between 425 nm and 475 nm.

**Detection of luciferase activity of *B. cinerea* P_*Bccpdh*:*Luc* transformant**

*B. cinerea* P_*Bccpdh*:*Luc* transformant was grown on PDA at 23°C. Three mycelia blocks (approx. 2 $mm^3$) were excised from the growing edge of the colony and submerged in 50 µl of water or indicated phytoalexin containing 50 µM D-luciferin in a sealed 96-well microplate (Nunc 96F microwell white polystyrene plate, Thermo Fisher Scientific, Waltham, MA, USA). Changes in luminescence intensity were measured over time with Mithras LB 940 (Berthold Technologies, Bad Wildbad, Germany).

**DNA sequencing and Bioinformatics**

DNA fragments were sequenced by the dideoxynucleotide chain termination method using Big-Dye ver. 3 chemistry (Applied Biosystems, USA). Products were separated on an ABI 3130 analyzer (Applied Biosystems). Sequence data was analyzed and annotated using MacVector (version 18.2 or earlier; MacVector Inc., Apec, NC, USA). Draft genome sequences of fungal species used for the analysis shown in Fig. 6, S20, S22-25 were obtained from Ensembl Genomes project (Ensembl Fungi, http://fungi.ensembl.org/index.html).

For phylogenetic analysis (Fig. S23), the deduced amino acid sequences were aligned by ClustalW (Thompson et al., 1994), and the phylogenetic tree was constructed using the neighbor-joining method (Saitou and Nei, 1987), and drawn using FigTree v1.4.4 (http://tree.bio.ed.ac.uk/software/figtree/).
